# The multiverse of data preprocessing and analysis in graph-based fMRI: A systematic literature review of analytical choices fed into a decision support tool for informed analysis

**DOI:** 10.1101/2024.01.14.575565

**Authors:** Daniel Kristanto, Micha Burkhardt, Christiane Thiel, Stefan Debener, Carsten Gießing, Andrea Hildebrandt

## Abstract

The large number of different analytical choices researchers use may be partly responsible for the replication challenge in neuroimaging studies. For robustness analysis, knowledge of the full space of options is essential. We conducted a systematic literature review to identify the analytical decisions in functional neuroimaging data preprocessing and analysis in the emerging field of cognitive network neuroscience. We found 61 different steps, with 17 of them having debatable options. Scrubbing, global signal regression, and spatial smoothing are among the controversial steps. There is no standardized order in which different steps are applied, and the options within several steps vary widely across studies. By aggregating the pipelines across studies, we propose three taxonomic levels to categorize analytical choices: 1) inclusion or exclusion of specific steps, 2) distinct sequencing of steps, and 3) parameter tuning within steps. To facilitate access to the data, we developed a decision support app with high educational value called METEOR, which allows researchers to explore the space of choices as reference for well-informed robustness (multiverse) analysis.

**Highlights:**

1. Data analysis variability in neuroimaging hinders replicability.
2. Analysis across multiple defensible options examines the robustness of results.
3. We conducted a systematic literature review to identify analytical options.
4. We identified 61 steps and 102 options in performing graph-fMRI analysis.
5. Interactive visualization of these steps and options is available as a Shiny app.

## 1. Introduction

Recent neuroimaging studies have shown that the organization of functional brain networks plays a crucial role in cognitive processing (e.g., Barabási et al., 2023; Bassett & Bullmore, 2017). Many previous studies have used graph theory or network science as an analytical framework to infer global and local properties of brain network topology and to investigate the association of brain network topology with cognitive abilities (for individual studies, see Gracia-Tabuenca et al., 2023; Kristanto et al., 2023; Van Den Heuvel et al., 2009; for a review, see Barabási et al., 2023; Blanken et al., 2021). Network science provides a comprehensive and integrated approach to understanding brain function by considering the complex interactions among multiple brain regions, and thus, the use of network science to study how the brain is organized to control cognitive processing is growing. There is, however, strong evidence that many empirical findings are highly dependent on the specific data preprocessing and analysis pipeline used. This calls into question the robustness of many reported findings, which has been discussed in the context of the replication challenge (Braun et al., 2012; Liang et al., 2012; S. Lin et al., 2018; Welton et al., 2015). A recent study involving 70 different teams analyzing the same dataset and testing the same hypotheses demonstrated the fundamental variability of the analysis pipelines used and the conclusions reached by each team (Botvinik-Nezer et al., 2020).

Potential solutions to the replication challenge, which may arise at least in part from the large number of degrees of freedom in data analysis, have been suggested. First, preregistration, which requires researchers to specify the details of their study design and planned analysis before the study is conducted, has been shown to improve credibility by reducing data-driven adjustments when creating the analysis pipeline (Nosek et al., 2019). Preregistration does, however, not allow researchers to draw conclusions for the entire choices of analytical approaches, although it is similarly important for a truly cumulative science to ensure the robustness of findings across equally defensible analytical options. Second, multiverse analysis has been proposed as an alternative to preregistration to mitigate the replication challenge, in part due to the large number of degrees of freedom in data preprocessing and analysis (Steegen et al., 2016). Multiverse analysis emphasizes the execution of all defensible analysis pipelines, including preprocessing steps, to bring transparency to the black box of exploration. This approach is also known as forking paths analysis (Gelman & Loken, 2013). For example, a forking path means that when testing a hypothesis about the association between brain network segregation and a behavioral outcome, researchers can consider several candidates for graph measures indicating network segregation, such as modularity or the clustering coefficient. A full multiverse analysis should then be followed by a full report of the results from all possible combinations of forking paths. This report might be, for example a *p*-value distribution, a specification curve analysis and plot, or a vibration of effects plot (for review, see Hall et al., 2022). A joint conclusion can then be drawn by integrating the results across the multiverse of options (Simonsohn et al., 2020) or by reporting a multiverse variability index that quantifies the robustness of the results across the forking paths (Olsson-collentine et al., 2023).

Multiverse analysis is a promising approach, but in neuroimaging coupled with the implementation of network science, it seems impossible for individual researchers to grasp the entire multiverse of defensible analysis pipelines to perform such analysis. Various aspects of the data contribute to the variability of the pipelines applied, creating forking paths that lead to potential differences in results across studies. First, functional magnetic resonance images (fMRI) are heavily affected by noise caused by the MRI machine, participant motion, and psychological processes unrelated to the measurement intentions that may also be active (see review by T. T. Liu, 2016). Therefore, data preprocessing steps aim to minimize the effects of noise prior to statistical analysis of brain-behavior associations. With the increasing number of computational methods developed for temporal and spatial preprocessing of fMRI data, a variety of analysis pipelines have been proposed (e.g., Caballero-Gaudes & Reynolds, 2017; Esteban et al., 2019; Glasser et al., 2013; Waller et al., 2022). Second, feature extraction from the brain network can be done in several ways. To give a simple example, the choice of the brain atlas that defines the nodes in the network has been shown to affect final conclusions (Bottino et al., 2022; Stanley et al., 2013). Several further complex decisions concern, for example, the chosen density of the brain networks, which will clearly influence the calculated graph metrics to be associated with psychological variables, and the resulting conclusions may vary depending on the corresponding analytical choices (Bordier et al., 2017; Dafflon et al., 2022).

There are several possible ways of identifying the complex multiverse of choices for an fMRI analysis pipeline. For example, expert workshops have been used to crowdsource analytical decisions (Paul et al., 2022). Multi-analyst project, such as the one published in the fMRI field by Botvinik-Nezer et al. (2020) is another option. However, both may be biased towards the participating experts and may not represent the entire literature on defensible decisions applied in a given research area. In the present work, we aim to define the multiverse of fMRI data preprocessing and analysis by focusing on network analysis using graph theory, through a systematic literature search and information extraction in order to better cover decisions made across the entire field. In these selected studies we focus on both the preprocessing of fMRI data to minimize the influence of noise, and the steps required to determine the brain network and parameterize its topological properties.

To our knowledge, the most recent systematic review of methodological decisions in fMRI preprocessing pipelines was published in 2012 (Carp, 2012). The main aim of that review was to document significant gaps in reporting practices across fMRI studies. We go beyond that scope here. Our study has a two-step approach: First, we perform a systematic review to investigate the multiverse in graph-based fMRI preprocessing and analysis. Second, we provide a decision support app that allows researchers to explore the space of choices and can serve as reference for well-informed robustness (multiverse) analysis. We envision that a decision support system can become an essential tool for individual researchers at all career stages to either design their own multiverse analysis pipeline, or to evaluate how similar their intended pipeline is to those already published. We also highlight the extent to which studies have not consistently reported their analytical decisions. Finally, we discuss how multiverse analysis can become a widely accepted und used analytical approach to infer not only the result itself, but also its robustness across analytical choices. We discuss the selection of defensible analytical decisions and the need of reporting guidelines.

## 2. Method

### 2.1. Specific aim of the systematic review

This review examines the variability in data preprocessing and analysis decisions employed by studies investigating the association between fMRI-based graph measures and behavioral outcomes. We limit our focus to the section of the pipeline from identifying the fMRI preprocessing software or toolbox to quantifying the graph measures, but do not focus on the methodological decisions that need to be made for the association models between brain-based graph metrics and behavioral measures.

In order to target a broad audience of the network neuroscience community working with fMRI data and to ensure the comprehensiveness, we conducted a two-stage literature review covering a wide range of potential fMRI preprocessing steps. We started with an initial literature search, aiming to include methodological papers addressing various technical aspects of fMRI preprocessing regardless of which kind of analysis of functional connectivity or local brain activity, and regardless of whether a brain-behavior association was investigated. The aim of the first review was to extract a complete picture of fMRI data preprocessing and to capture the most recent methodological advancements in techniques from general fMRI studies. The main output of the first review was the list of steps used in fMRI data preprocessing (see Results section 3.1). Next, we narrowed down our focus to network neuroscience, with the second literature review targeting fMRI studies that used graph measures and explored their relationship to cognitive ability. The aim of the second review was to examine how studies in cognitive network neuroscience used the steps identified in the first review to preprocess fMRI data. In particular, the second review also contributes to the list of steps beyond fMRI data preprocessing, namely steps related to the definition of the brain network for graph analysis. The first review will therefore be referred to in the remainder of the article as *Review 1* and the second review as *Review 2*.

### 2.2. Search strategy

To ensure transparency and adherence to established standards, we followed the Preferred Reporting Items for Systematic Reviews and Meta-Analyses (PRISMA) guidelines for reporting systematic reviews (Page et al., 2021). A visual representation of the entire search and selection process is provided in Figure 1, as shown in the respective flowchart.

**Figure 1.**
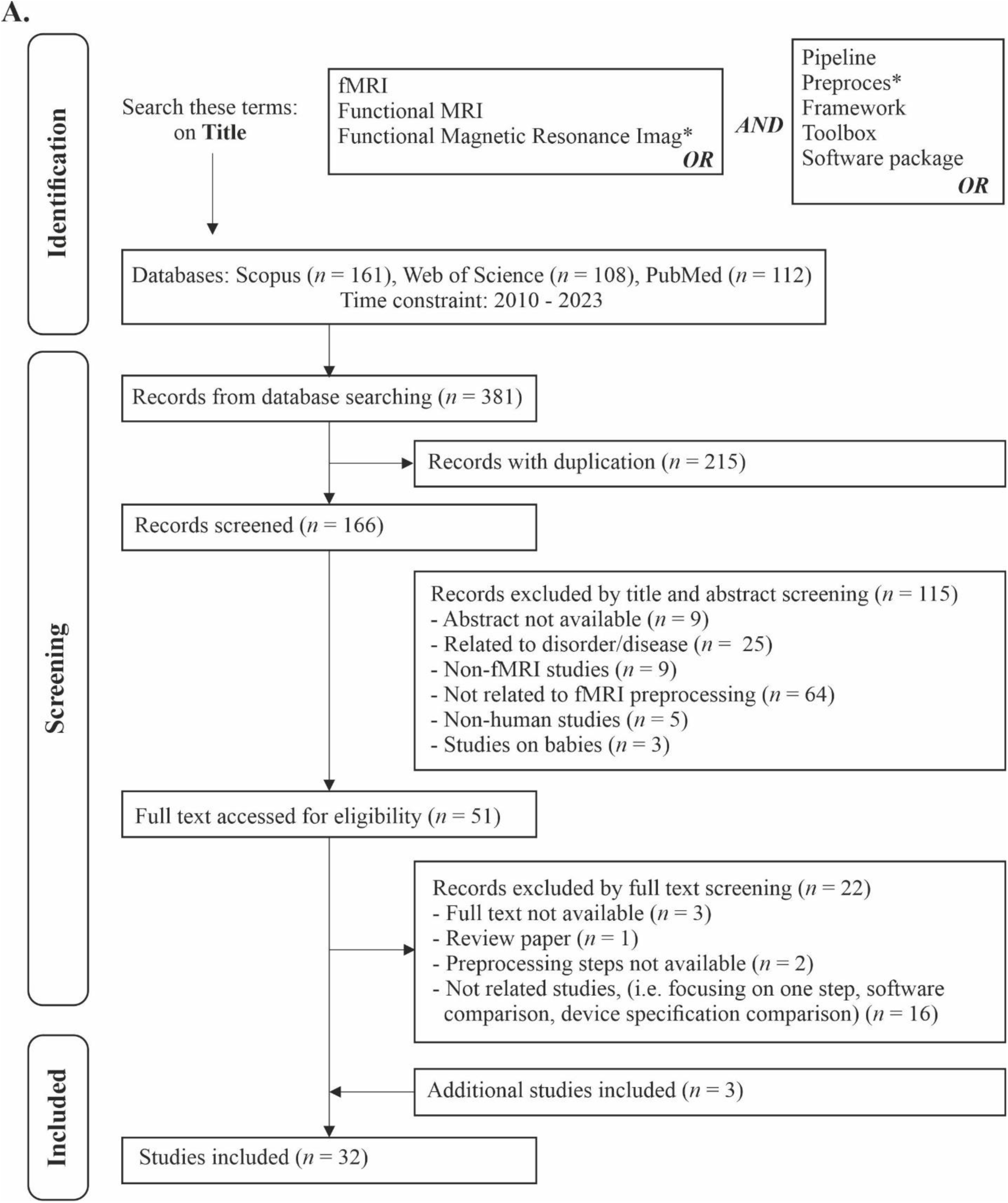
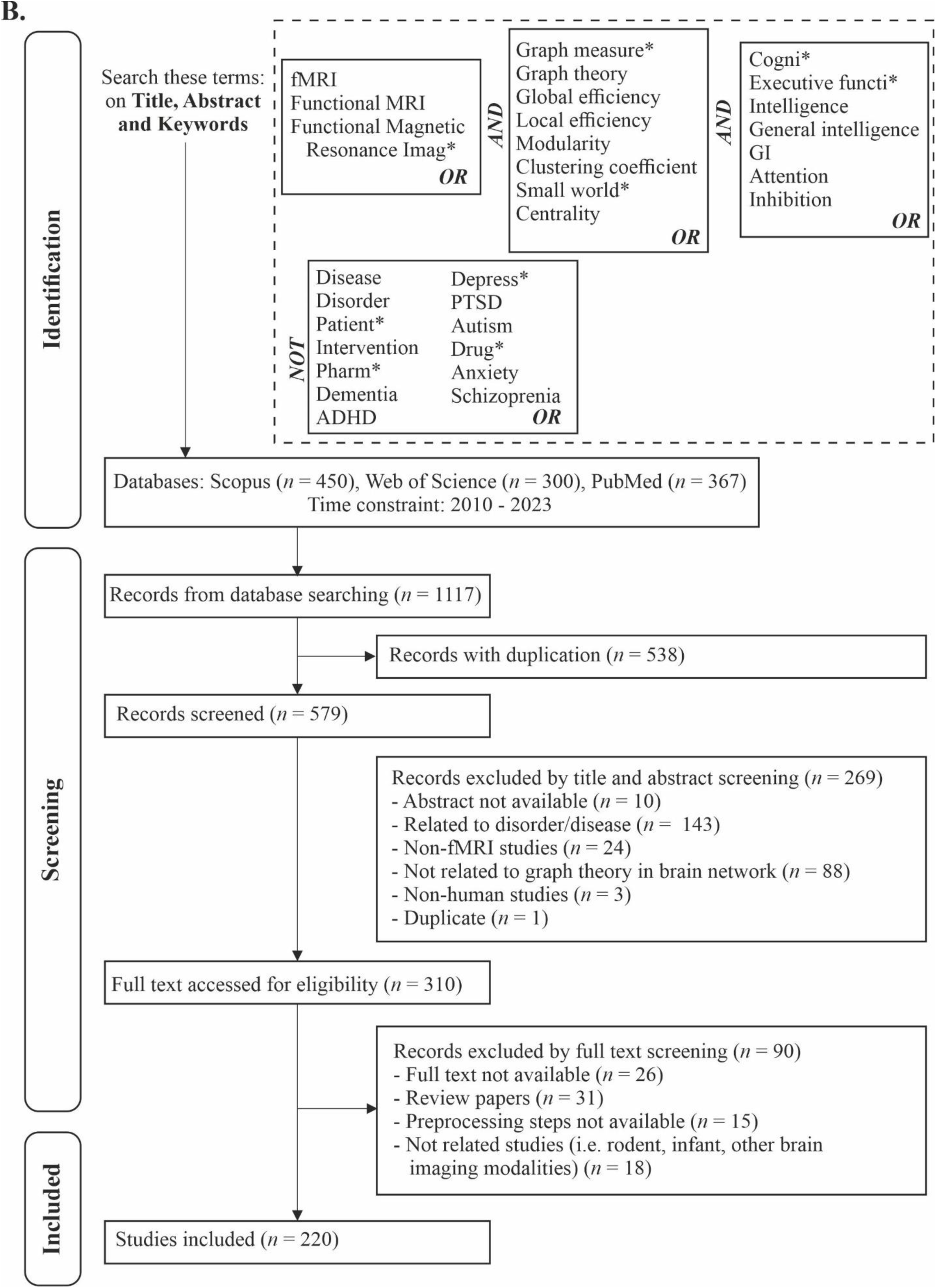
PRISMA diagrams to report the systematic review process of Reviews 1 and 2. Panel (A) shows a PRISMA diagram for Review 1 describing the identification, screening and selection process and panel (B) shows the same for Review 2.

We employed different search strategies for Review 1 and Review 2, as illustrated in Figure 1. For Review 1, the search focused on study titles containing various terms related to methodological aspects of fMRI research like ‘pipeline’ or ’preprocessing’. For Review 2, we extended the search to the title, abstract, or keywords fields and included more terms related to graph theory and cognitive ability terminology. The disparity in the scope of the search space (title for Review 1 and title, abstract, keywords for Review 2) resulted in a lower number of identified articles in Review 1 compared to Review 2, despite the use of broad search terms in Review 1. Furthermore, to maintain the focus on the healthy brain, we used a ’NOT’ Boolean gate to exclude studies related to brain or mental disorders. For both reviews, we applied a temporal constraint, limiting the search to studies published between 2010 and 2023, aiming to capture up-to-date methods in a dynamic and rapidly evolving methodological field. Furthermore, it is important to acknowledge that Review 2 is not a subset of Review 1 due to their distinct objectives. Review 1 primarily focuses on studies that introduced or proposed a set of preprocessing techniques for fMRI data, while Review 2 encompasses studies that exclusively employ these techniques, with the extension of graph-fMRI approach and the association with behavior.

The search was carried out in three databases: Scopus, Web of Science, and PubMed, on August 9^th^, 2023. To further refine the search results, we extracted the metadata containing the publication information of the articles in the form of a bibliography document (bib files) of the identified studies from each database. Using the RevTools (Westgate, 2019) package in the R Software for statistical computing (R Core Team, 2020), we undertook further preprocessing of the metadata. Initially, we concatenated the studies from different databases and eliminated duplicate entries by applying functions to identify duplicated items `(find_duplicates)` and to extract the unique items `(extract_unique_references)`. To identify such duplicates, both Digital Object Identifier (DOI)-based and title-based methods were employed. Thus, articles with similar DOI and/or titles were identified as duplicate and only one was retained. This resulted in 166 and 550 studies for Review 1 and Review 2 respectively, which were then subjected to further screening.

### 2.3. Screening phase and eligibility criteria

During the screening phase, we excluded studies that lacked relevance based on an assessment of their titles and abstracts. To facilitate this process, we used the `screen_abstract` function from RevTools, which provides a user-friendly interface for reading and annotating abstracts. A number of exclusion criteria were applied consistently for both Review 1 and Review 2, such as excluding studies involving brain or mental disorders, non-human subjects, and non-fMRI investigations. As a result of the screening phase, we were left with 51 and 296 studies, respectively, that met the initial eligibility criteria and were considered for further assessment.

In the subsequent eligibility assessment phase, we reviewed the full-text documents for the identified studies to determine their relevance. For this purpose, we used Zotero (www.zotero.org), a reference management software, to download the full text from the publishers. Specifically, we compiled a list of the DOIs associated with the studies and inserted them into Zotero to facilitate automated full-text retrieval. In cases where Zotero was unable to download certain papers, we obtained them manually from the publishers. During the eligibility assessment, for both Review 1 and Review 2, we focused exclusively on human studies. In addition, only studies that used fMRI-based graph measures and examined their association with behavioral variables were selected for Review 2. In both reviews studies using alternative imaging modalities other than fMRI and those conducted on non-human subjects were excluded. We also excluded studies involving infants because of potential differences in the approaches used to preprocess fMRI data for this population.

Finally, review articles and studies that did not report fMRI preprocessing steps were eliminated from review 1 and 2. We added three further studies on fMRI preprocessing pipelines from three fMRI research consortia (Human Connectome Project (HCP, Glasser et al., 2013), Enhancing Neuro Imaging Genetics Through Meta Analysis (ENIGMA, Waller et al., 2022), and UK Biobank (Alfaro-Almagro et al., 2018)) to Review 1 that were highly relevant to fMRI preprocessing. Those studies were not identified by the search because they did not contain either fMRI-related or preprocessing-related terms in the title. As a result, the final sample consisted of 32 studies for Review 1 and 220 studies for Review 2.

### 2.4. Data extraction procedure and quality assessment

As mentioned above, the aim of the present review was to extract methodological decisions on the pipeline from the identification of the fMRI preprocessing software/toolbox up to the quantification of the graph measures to be associated with behavioral variables at a later stage. The choices made in estimating the brain-behavior association model are not the subject of this review. To extract and categorize methodological decisions regarding the processing of the data up to the quantification of the graph measures, we used two different terms to denote distinct actions: Step(s) and Option(s). Steps refer to specific actions taken to preprocess or analyze the data, such as motion or temporal filtering. Options indicate when there were multiple choices or variations available within a particular Step, for example selecting different parameter options for motion regression (e.g., 6/12/24/36-parameters) or applying a low-pass, band-pass, or high-pass filter for temporal filtering. Notably, we coded only the Options of selected Steps that were frequently reported and varied across studies in the literature. It is important to note that the extracted Steps in Review 2 ended with the quantification of graph theory metrics, whereas Review 1 ended with the determination of the analysis path up to the calculation of a cleaned time-series fMRI signal.

The extraction of Steps and Options from the eligible studies was mainly carried out by the first author (DK). To estimate inter-rater consistency, two additional coders (co-authors MB and CG) performed an identical coding procedure on a subset of 25 randomly selected studies and compared their results with DK’s original coding. We refer here to each piece of information from a study as an item, including the preprocessing Steps taken by the studies and the corresponding Options. To estimate the inter-rater consistency, we calculated the number of items in which the coders differed. Out of a total of 652 and 689 coded items for authors DK and MB vs. DK and CG respectively, there were 53 (8.13%) vs. 32 (4.64%) discrepancies (see details in the Database.xlsx file at https://github.com/kristantodan12/fMRI_Multiverse/). To resolve these disagreements or differences in interpretation between the coders, a collaborative discussion was held to reach a consensus and determine the most appropriate coding. Finally, author DK re-checked the coding for all other papers after discussion with authors CG and MB. This approach ensured that the coding process was rigorous and aimed to minimize subjective bias, ultimately increasing the reliability and accuracy of the results.

### 2.5. Interactive app for data visualization

An interactive app was created using the *Shiny* toolbox (W. Chang et al., 2023) in the R Software for statistical computing. This allowed to visualize the Steps and Options adopted by each study selected for this review. Unlike static figures and tables, the Shiny application allows users to interactively select studies and explore their specific preprocessing and analysis pipelines. It also provides other features to facilitate a deeper exploration of the database of fMRI data preprocessing and analysis decisions that we created as a result of the systematic literature review. The app is available at https://meteor-oldenburg.shinyapps.io/fMRI_multiverse/. The code and the data to create the app are available at https://github.com/kristantodan12/fMRI_Multiverse/. In keeping with the title of the project this work is part of, we call the app ‘METEOR’ (MastEring ThE OppRessive number of forking paths unfolded by noisy and complex neural data).

METEOR consists of several tabs, each with a distinct function:

1. The initial tab, *Introduction*, provides an introduction to the application, an overview of its features, and information on how to use them.
2. The next tab, *Database*, lists all the data collected, organized into various subsections. These include the PRISMA diagrams (subsection *PRISMA Diagram*), a list of papers included in the Review 1 and Review 2 (subsection *List of Included Articles*), and lists of Steps (subsection *List of Steps*) and Options (subsection *List of Options*) identified during the analysis.
3. Next, users can explore the aggregated pipelines across studies in the *Steps* tab. This tab contains several subsections: *Aggregated Steps*, *Individual Step, Combination*, and *Order*. The subsection of *Aggregated Steps* visualizes the aggregated pipelines in a network fashion, where the nodes represent preprocessing or analysis Steps, and the edges represent the connection between two Steps. By hovering and clicking on the nodes/edges, the user gets information about the respective Steps and about how many papers used them. A definition of the Steps is additionally provided by clicking the nodes. There are also further settings on the left-hand dashboard to specify the Step to be displayed and the threshold for the number of papers that have used the given step, indicated by the edges. In addition, user can also explore the papers used a specific tab in *Individual Step* subsection. The *Combination* subsection allows users to explore pairs of functional preprocessing Steps used in combination. By selecting a specific Step, such as global signal regression for example, a “lollipop” plot is generated to indicate the frequency with which other Steps have been used in conjunction with global signal regression. Finally, the *Order* subsection allows users to examine the order of certain Steps relative to others. For example, if a user selects the motion regression step, a bar plot is generated to illustrate how many studies implemented other steps after motion regression.
4. The next tab, *Steps: Options*, allows the user to visualize the distribution of Options chosen by different studies. For example, users can examine the distribution of software used to preprocess fMRI data. Within this tab, one can also specify a particular Option, such as the software package SPM (Statistical Parametric Mapping), and obtain a list of studies that have used this Option.
5. Users can explore and visualize the Steps and Options taken by an individual study in the *Individual Paper* tab. It contains two subsections, *Step Visualization* and *Option Visualization*, which allow users to interactively explore the preprocessing and analysis pipeline and the specific options chosen for each study.
6. We have also incorporated an important feature called *Your Own Pipeline*, where users can enter their preferred pipeline for preprocessing fMRI data and associated options. As a result, the application will provide a count of the number of studies that have used the same pipeline and a list of those studies. Importantly, users can specify whether the order of Steps within their pipeline should be taken into account. If this feature is disabled, the algorithm will count the number of papers that have used the preprocessing pipeline specified by the user, regardless of the order in which the respective Steps were applied.
7. Finally, the last tab, labelled *About,* provides information about the project and the research team that created the resource.

## 3. Results

There are three subsections to the results section: 1) list of the coded Steps and Options, 2) variability in Steps, and 3) variability in Options. Given the primary focus of this review on network science implementation in fMRI data, the findings regarding the variability in preprocessing and analysis steps and options are derived from the preprocessing and analysis pipelines identified in Review 2. The aim is to provide a detailed picture of the multiverse of the fMRI data preprocessing and analysis pipelines, derived from the variability of Steps and Options as used in the literature. We also provide examples illustrating the use of the METEOR app to address controversial issues in the literature related to fMRI data analysis pipelines. In particular, some of the figures in the Results section are taken from the METEOR app. We invite readers to use the app while reading this paper to explore the findings in relation to other Steps and Options not directly illustrated here.

### 3.1. List of the coded Steps and Options

First, we identified the distinct Steps included in the different fMRI data preprocessing pipelines. In Review 1, we identified 49 unique Steps that can be grouped into three overarching domains: *structural image preprocessing*, *functional image preprocessing*, and *noise removal*. It is worth noting that, as mentioned above, the endpoint of these steps in Review 1 is the determination of a noise-free time-series fMRI signal. Furthermore, we emphasize that the Steps included in the preprocessing of structural images contain the procedures necessary to prepare the structural images for the subsequent standardization of the functional images to a common template. Thus, we excluded any structural image preprocessing steps aimed at calculating morphometric features of the brain, such as cortical thickness or brain volume measures which were not directly required to inform the analysis of functional brain data.

Expanding on Review 1, Review 2 contributed additional 12 Steps, grouped into the categories *functional connectivity definition* and *graph measure computation*. Notably, we also found a significant, additional Step used by certain studies that went beyond the quantification of graph measures. This particular step indicates how studies aggregate the results of different decisions made during data preprocessing and analysis, such as different thresholding techniques applied to the brain graph. Some studies chose to proceed with the most robust results across thresholding techniques, while others chose to average the results across different analytical decisions. Given the relevance of this step to multiverse analysis, we included it in the list of Steps, even though it goes one step beyond the computation of graph theory metrics.

In summary, the literature review yielded a total of 61 distinct Steps, which were grouped into five distinct domains: *structural image preprocessing*, *functional image preprocessing*, *noise removal*, *functional connectivity definition*, and *graph analysis*, as outlined in Table 1. Among these 61 Steps, we identified 17 Steps with multiple Options that have been frequently reported and varied across studies in literature. These Steps have a variable number of reported options, ranging from 2 to 20, resulting in a total of 102 Options distributed across the 17 Steps, as also summarized in Table 1. The lists of Steps and Options are also available in the METEOR app under the *Database* tab. Moreover, the list of studies used these Steps and Options is available in Table S1 of the supplementary material.

**Table 1.**
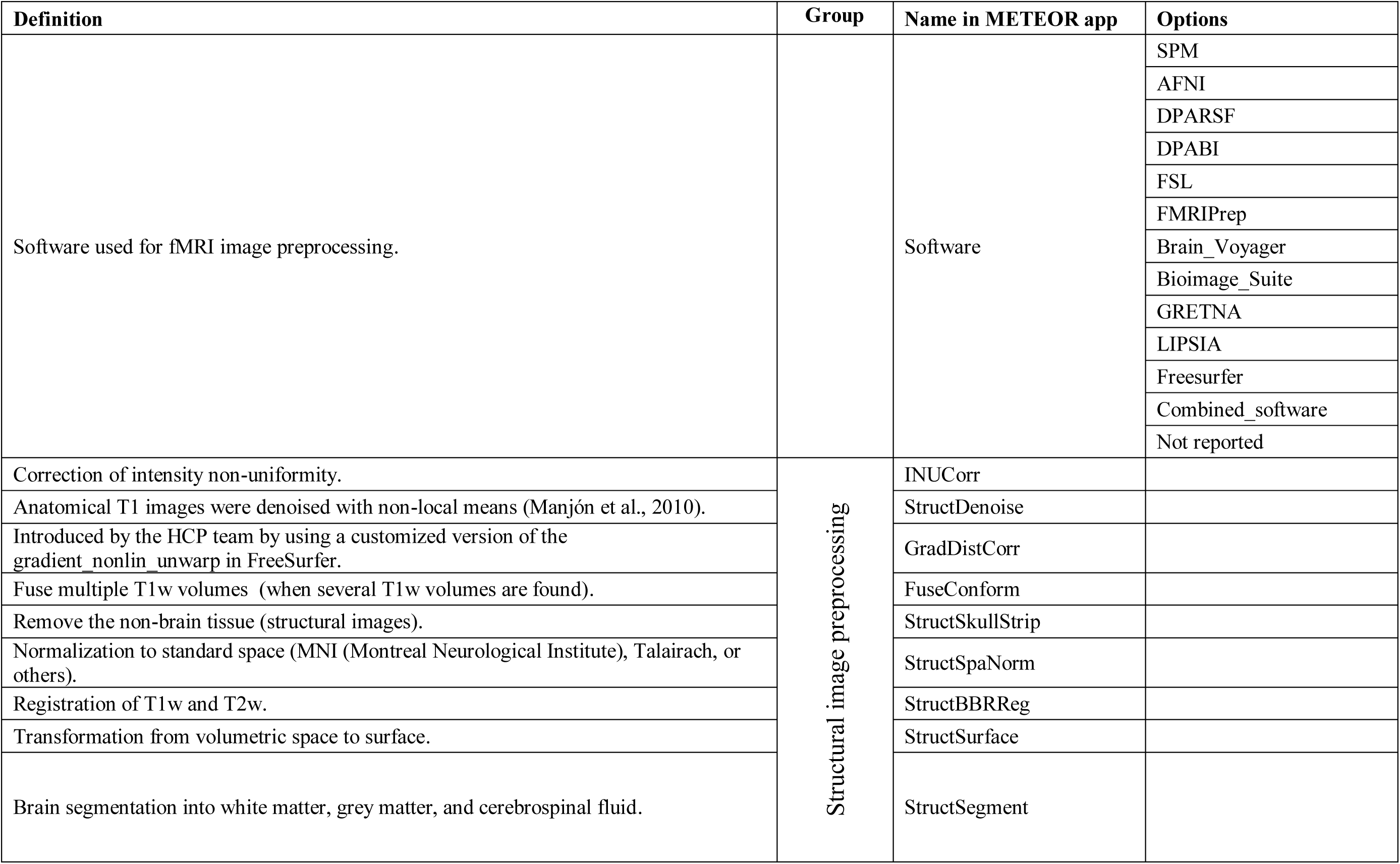
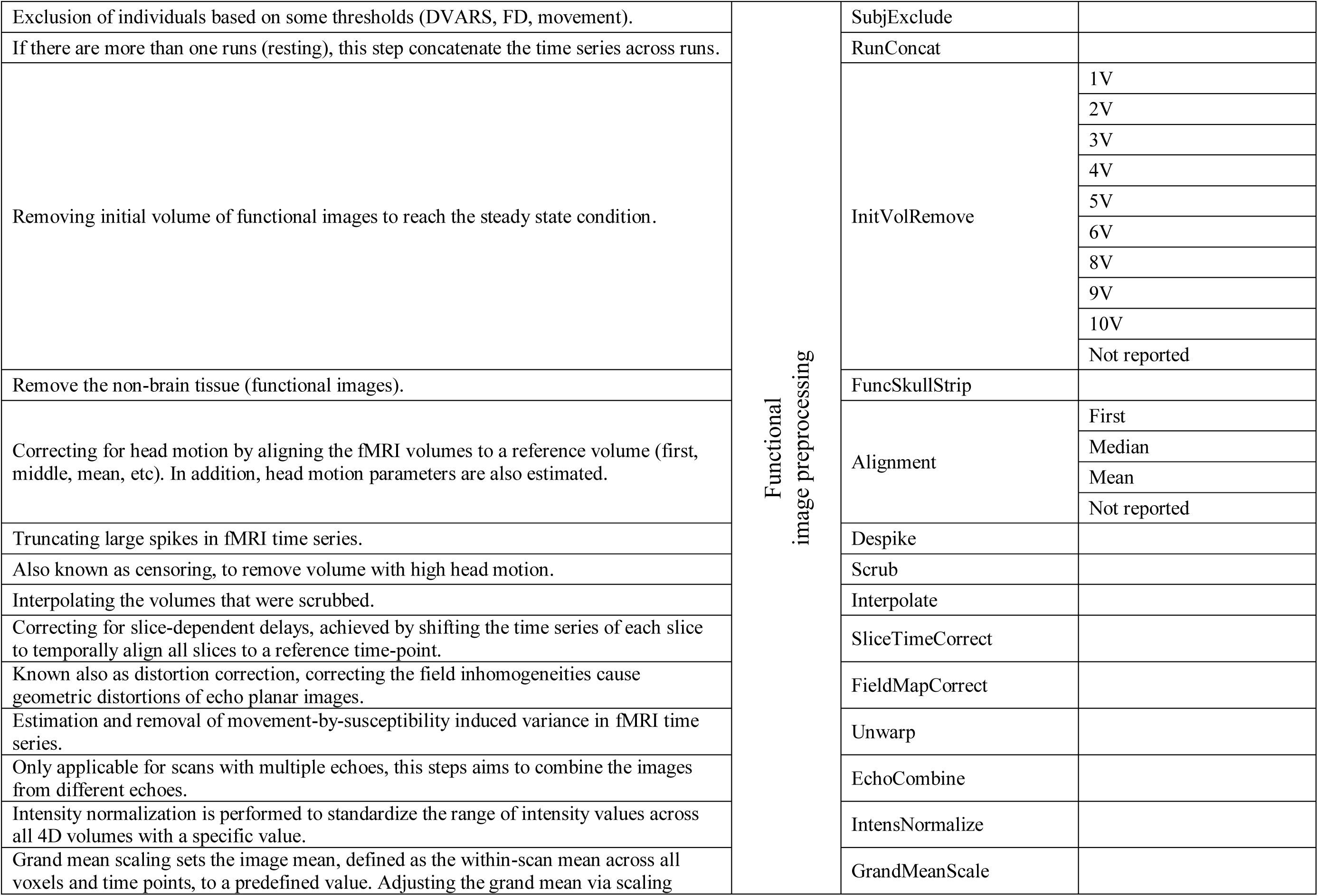
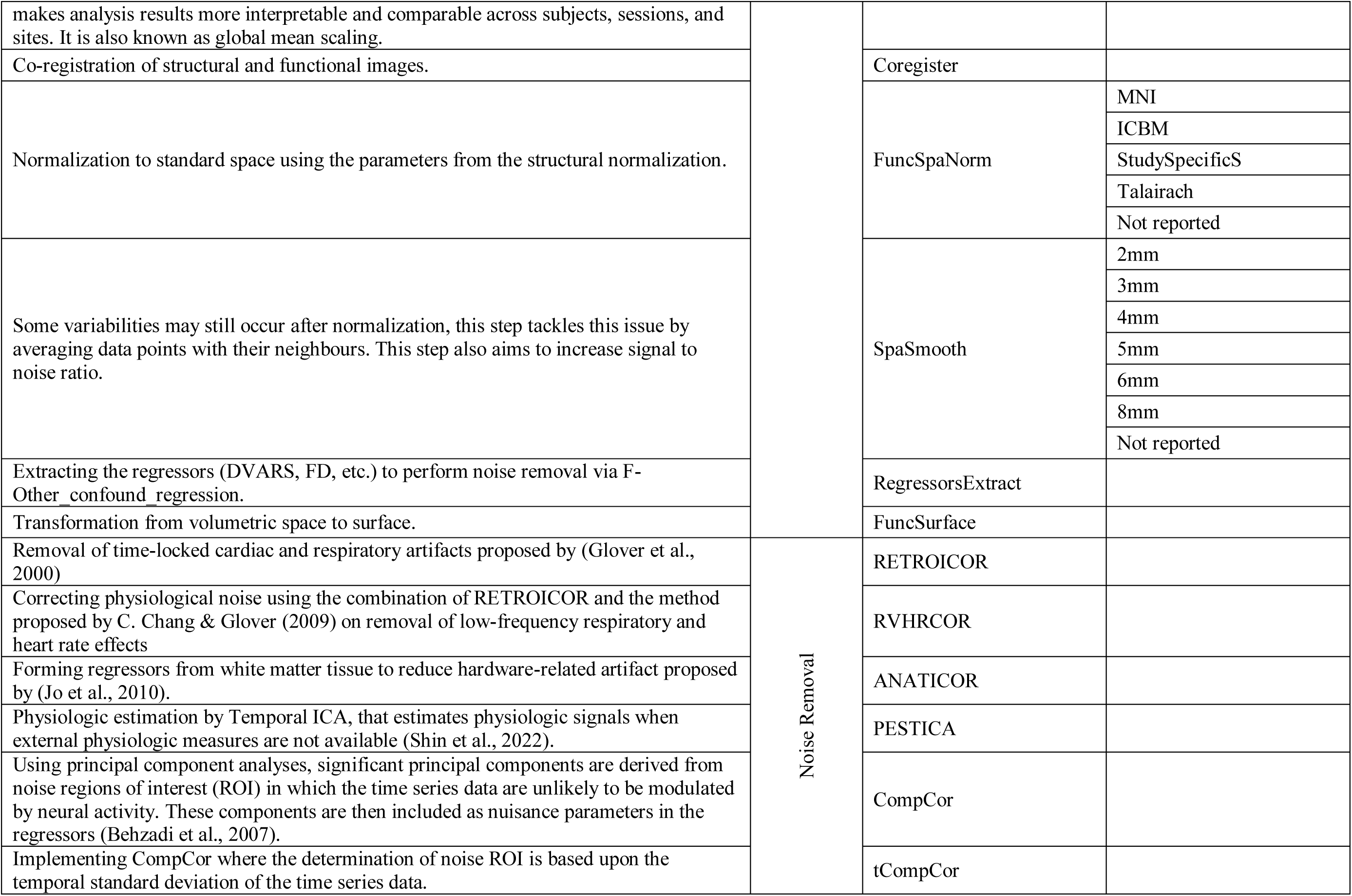
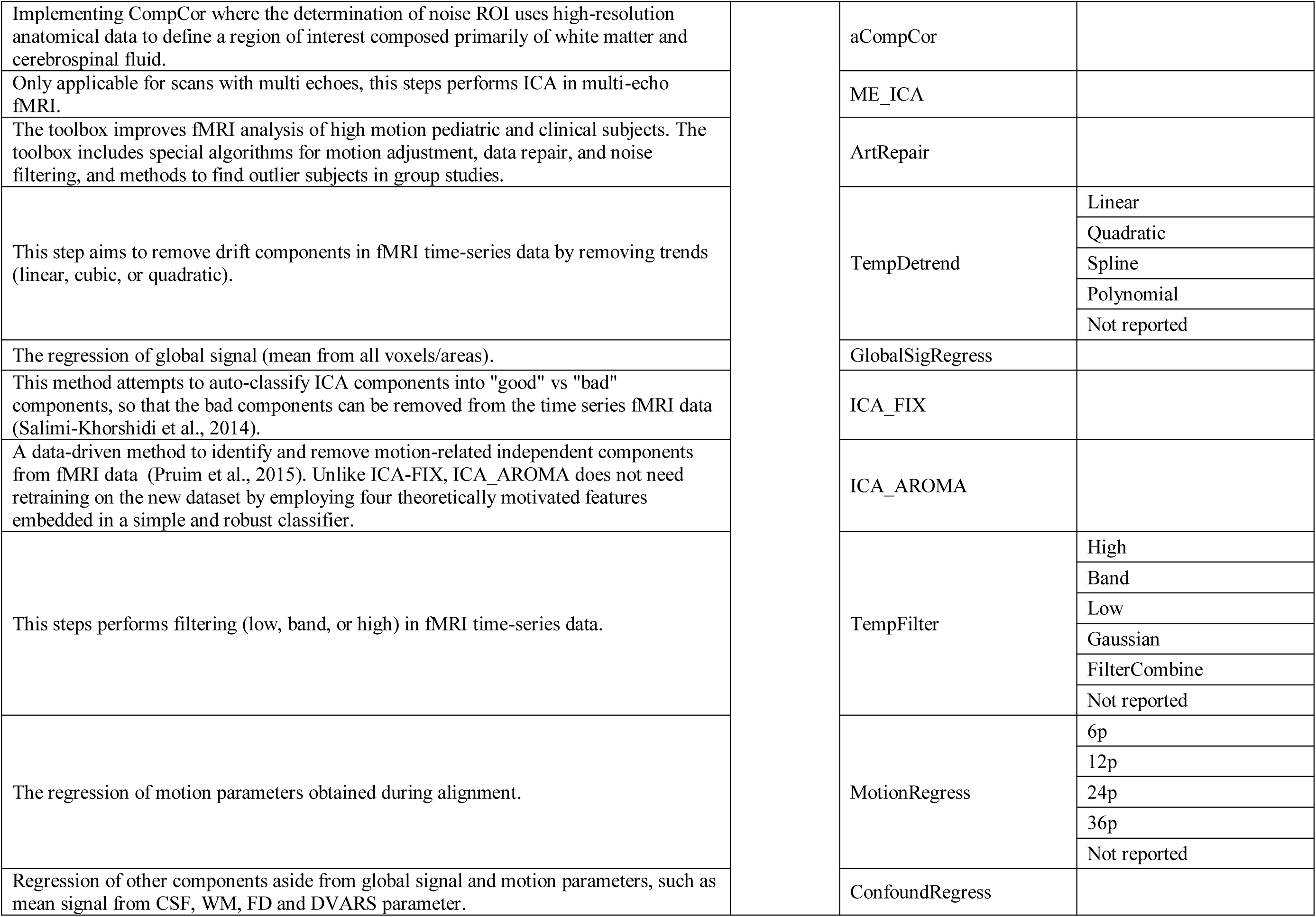
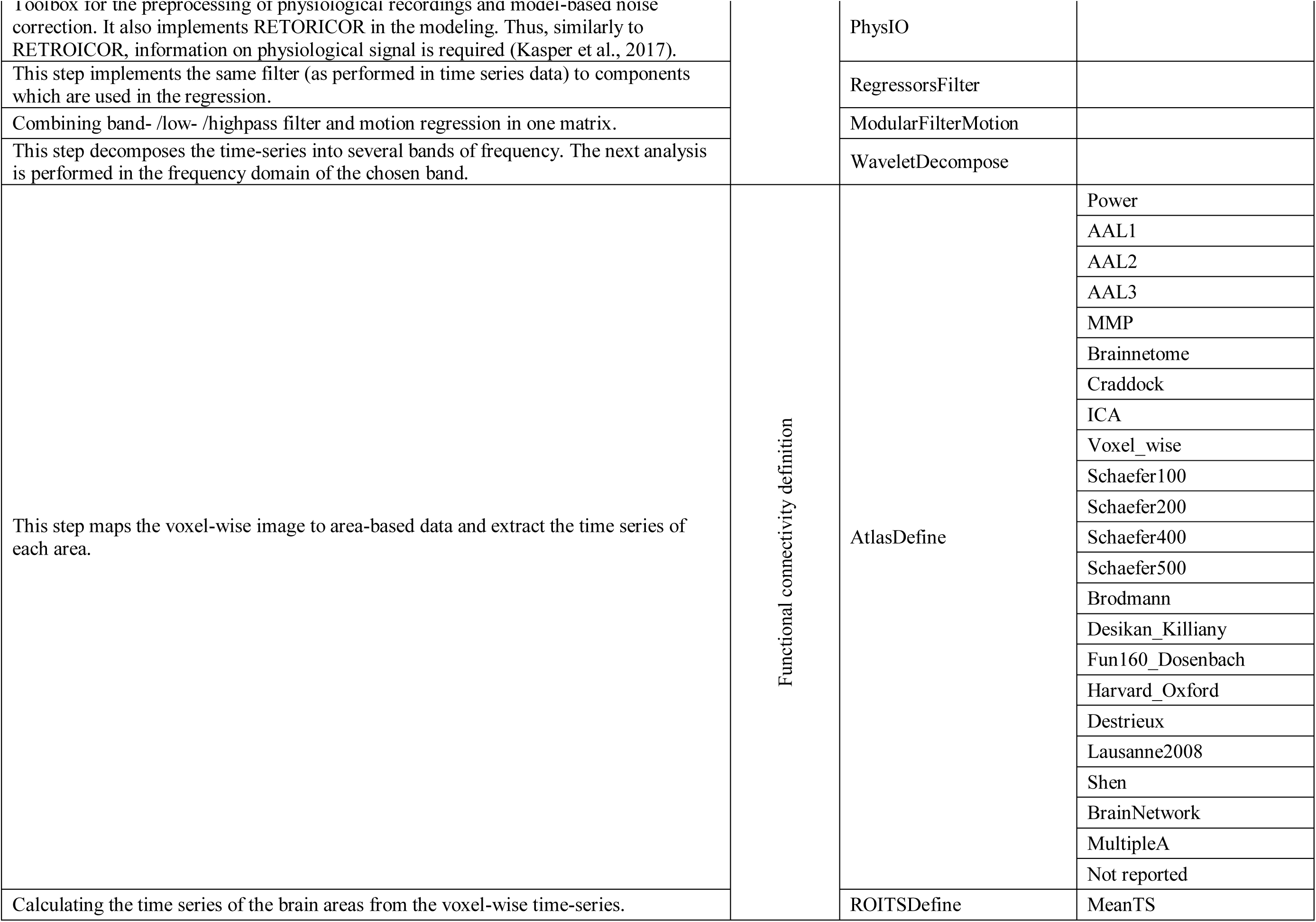
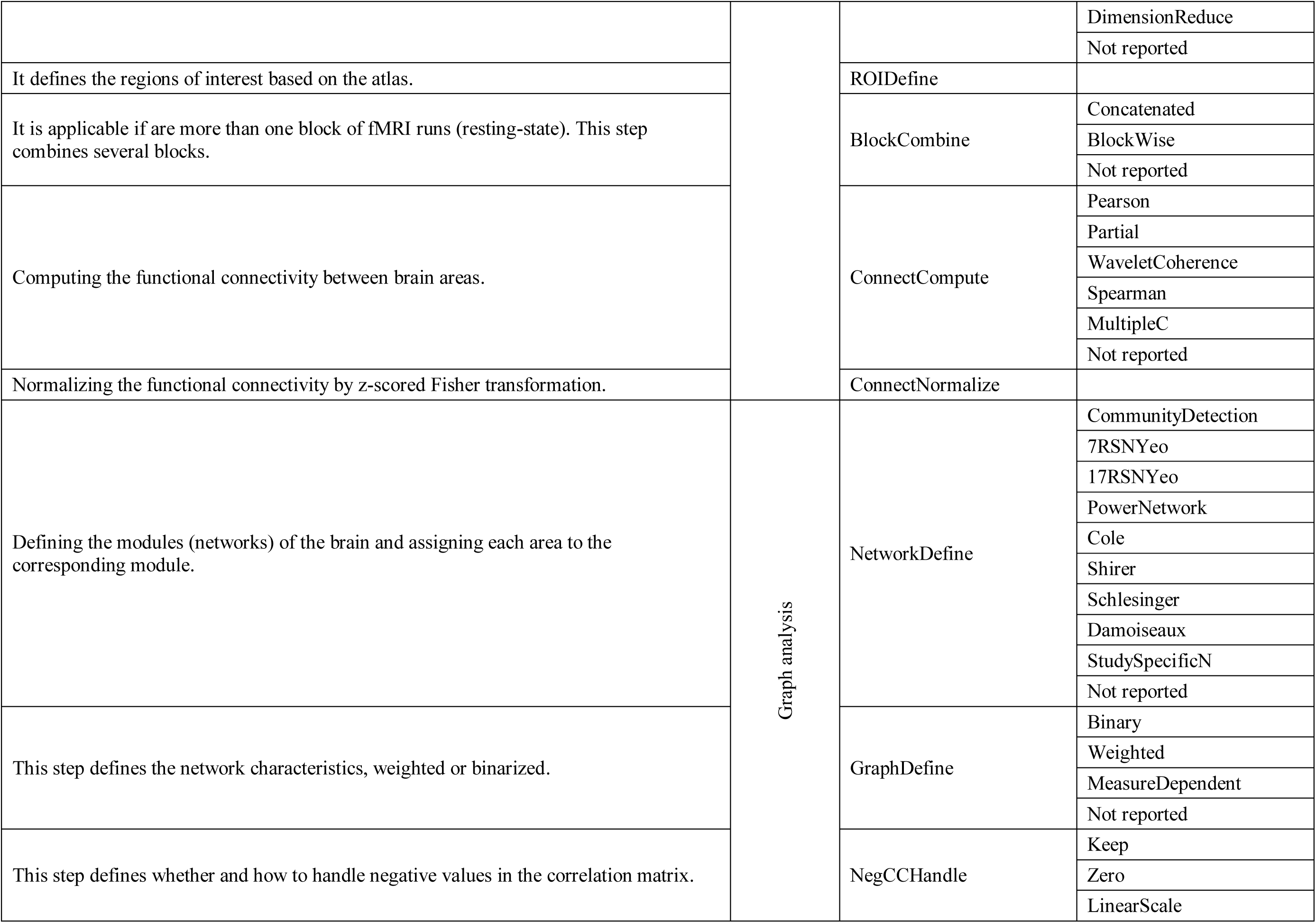
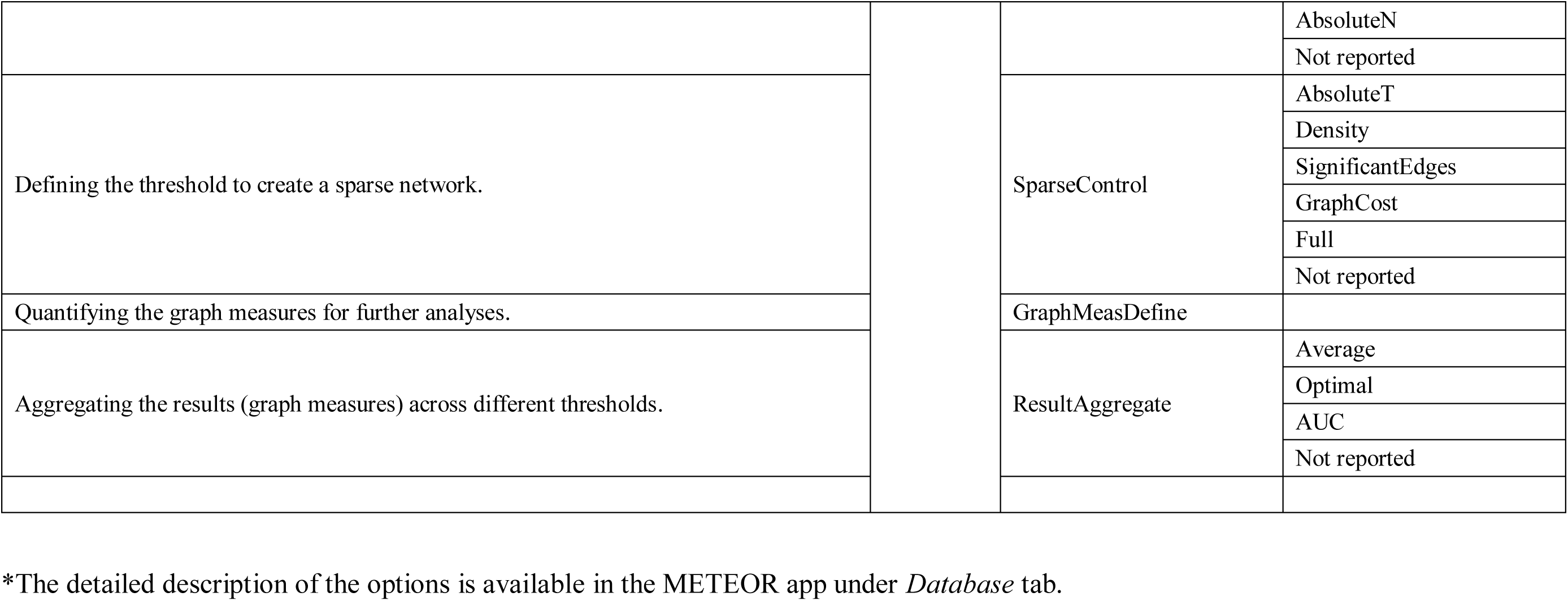
List of the identified Steps and Options.

### 3.2. Variability in Steps

#### 3.2.1. Aggregated pipeline across the studies

This section reports the results in terms of the variability of the steps taken by the studies from Review 2, as the focus of this review is on graph analysis of fMRI data. After coding the fMRI data preprocessing and analysis pipelines of each study and documenting them, there were a total of 220 pipelines from 220 studies in Review 2. This means that no two studies used exactly the same pipeline. As a first analysis, we aggregated these pipelines across studies to examine the most common decisions made in a subsequent Step, given a reference Step. This analysis also aimed to identify any deviations from these most common decisions. This aggregated pipeline is visualized in Figure 2. The nodes in the graph represent the individual Steps in the pipeline, while the directed edges (arrows) represent the connections between two Steps. For example, an arrow pointing from *ROIDefine* to *ConnectCompute* indicates that the *ConnectCompute* Step follows the *ROIDefine* Step in the pipeline. In the following, we will refer to the connections between the Steps as paths. Figure 2 includes certain features of graph visualization. First, the nodes are color-coded to represent their respective domains out of the five mentioned above. In addition, the size of the nodes corresponds to the number of studies that have performed the associated Step, with larger nodes indicating a higher frequency of occurrence. Similarly, the thickness of the edges is proportional to the number of studies that have followed the corresponding paths. The color of the edges is also coded to provide further information: grey edges represent paths taken by at least one study, blue edges represent paths used by more than 5 studies, and red edges represent paths taken by more than 20 studies. Note that the number of papers using a particular path/edge only takes the source of the edge (initial step) as a reference, regardless of the Steps (analytical decisions) taken before the source.

**Figure 2.**
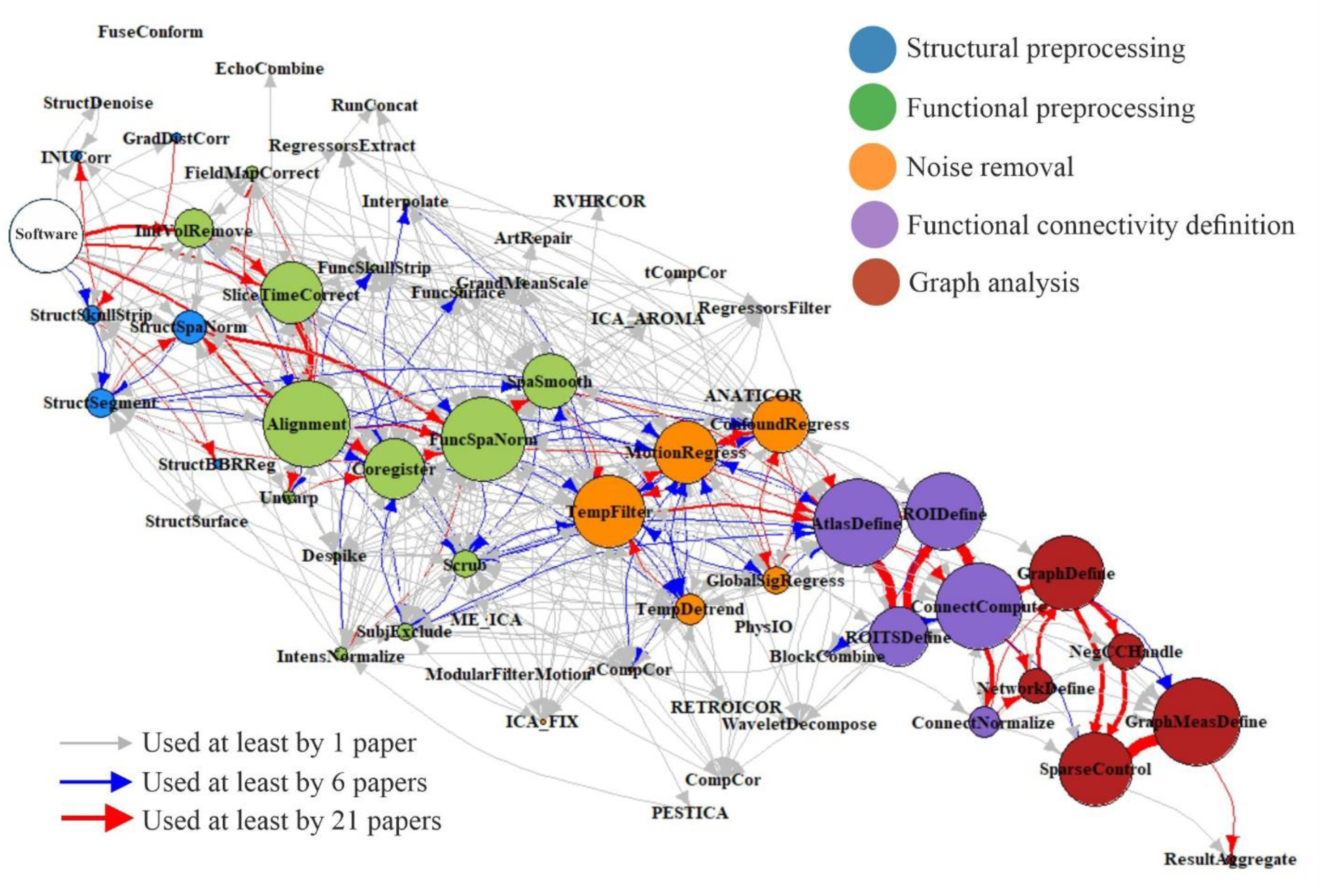
Aggregated pipeline across studies visualized as a graph. The fMRI data preprocessing and analysis pipelines extracted from the literature were aggregated across studies and visualized as a graph network. The nodes represent the Steps and the edges are the connections between the Steps. The size of the nodes and the thickness of the edges both scale linearly with the frequency of the Steps; larger nodes and thicker edges both indicating that the Steps were used in more studies. The color of the nodes (Steps) indicates the domain category of the node, while for the edges, black, blue, and red paths indicate that these connections have been implemented by at least 1 paper, more than 5 papers (6 or more), and more than 20 papers (21 or more), respectively.

Focusing on the red edges, which represent the most common paths connecting two steps, as they were followed by more than 20 studies, the analysis revealed a clear start-to-end pipeline for fMRI data preprocessing and analysis, with a high degree of agreement across studies. However, it must be noted that the number of papers associated with a specific path (pair of steps) was counted independently of the preceding or subsequent paths. In other words, the path shows only the most likely Step following a certain Step. The common path with red edges begins with software definition and removal of initial fMRI volumes to obtain a steady-state condition. The common path encompasses slice timing correction, alignment of functional images to a reference image, and structural image co-registration to normalize the functional images to a standard space. Once standardized functional images are obtained, the pipeline continues with noise removal steps, including spatial smoothing, motion influenced volume scrubbing, temporal filtering and regression of motion parameters and other confounders such as white matter (WM) and cerebrospinal fluid (CSF). At this stage, a cleaned fMRI time-series can be expected. Subsequent Steps are related to functional connectivity definition and graph analysis. The voxel level time-series data are then used to extract the area/region level time-series using a brain atlas. Functional connectivity is computed across pairs of brain areas and normalized across the whole brain to create the brain graph. Notably, most studies agree on several crucial analytical decisions to define the brain graph, such as controlling graph sparseness, handling negative edges, and determining edge weights (whether they are binary or weighted). Finally, graph measures are computed and aggregated across different analytic choices.

In addition to visualizing the most common paths connecting two Steps, Figure 2 also provides insight into how studies deviate from it, particularly through the grey and blue paths. It can be observed that discrepancies often occur in Steps related to the preprocessing of structural images, with some studies implementing and reporting more detailed approaches to preprocessing structural images before using them to normalize functional images. Another notable difference is in the noise removal Steps, with certain studies using more advanced techniques to distinguish noise from brain-related signals in fMRI time-series. These techniques include *CompCor* (Component Based Noise Correction Method) and its derivatives, RETORICOR (Retrospective Image Correction), ICA-FIX (Independent Component Analysis - FMRIB’s ICA-based X-noiseifier), and ICA-AROMA (ICA-based strategy for Automatic Removal of Motion Artifacts; see their more detailed description in the METEOR app). Other factors contributing to these deviations include grand mean scaling, global signal regression, and temporal detrending. Importantly, we did not find significant deviations in the Steps related to functional connectivity definition and graph analysis. This suggests a well-established consensus in these Steps of the data analysis pipeline.

Figure 2 also provides insight into the most controversial Steps among researchers. We defined the Steps as those performed by a moderate percentage of studies (approximately 30-70%, 60 to 150 studies across the 220 studies), indicating that they were not universally adopted but also not uncommon. These controversial steps include scrubbing, global signal regression, spatial smoothing, temporal detrending, motion and other confound regression, functional connectivity normalization, and the handling of negative correlations. Notably, these analytical decisions are highly debated in the literature (Braun et al., 2012; G. Chen et al., 2011; Liang et al., 2012).

#### 3.2.2. The estimated degree of the Steps

To further investigate the use of a particular Step relative to other Steps, we performed graph-based analysis on the pipeline summary extracted from the literature. We transformed the aggregated pipeline across all studies into a binary undirected graph, where nodes represent individual Steps and edges symbolize the connections between them, and computed the degree measure for each node to explore the variation in Steps connected to a particular node. Degree refers to the number of edges of a particular Step, where Steps with a higher degree coefficient indicate that they are connected to a more diverse set of Steps. Figure 3 shows the degrees of all the Steps. Interestingly, this analysis revealed that Steps associated with noise reduction techniques, such as temporal filtering, temporal detrending, motion and other confound regression, despiking, and spatial smoothing, had high degree coefficients. Similarly, functional image alignment and slice timing correction were Steps with notable degrees. This finding suggests that there is no clear agreement on which combination of these Steps should be performed. Conversely, lower degree Steps are predominantly associated with functional connectivity definition and graph analysis, where there is a commonly agreed combination of execution. Figure 3 also illustrates the steps connected to spatial smoothing of functional images (SpaSmooth) and temporal filtering (TempFilter) as two steps with high degree. The graph-based figures of these steps were taken from the METEOR app and can be created in the *Steps* tab. Notably, the visualization on the METEOR app also informs about the direction of the edges that contribute to the degree (in-degree: previous Step, out-degree: following Step). Readers can visualize the connections of other Steps (not only those shown in the static Figure 3) through the METEOR app by selecting the respective Step on the dashboard.

**Figure 3.**
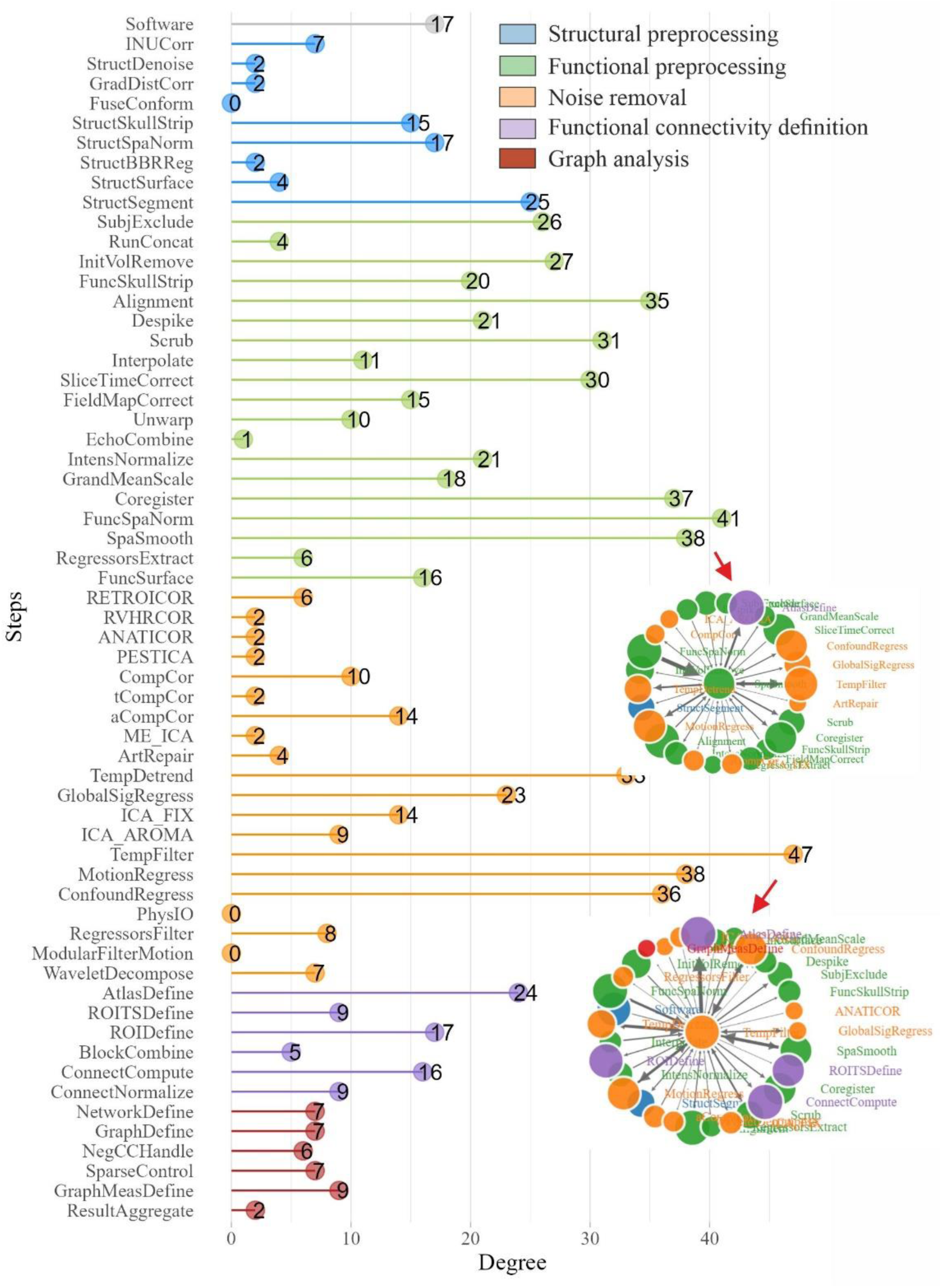
Degree coefficient of each Step of the preprocessing and analysis pipeline extracted from the literature. The lollipop plot shows the degree measure of each Step, where a higher degree indicates that the Step is connected to a high number of other Steps. Steps with the highest degree are mostly those associated with removal of noisy signal, such as spatial smoothing (SpaSmooth) and temporal filtering (TempFilter). The visualization on the METEOR app can also inform the direction of the edges that contribute to the degree (in-degree: previous Step, out-degree: following Step). Whereas high indegree means that these steps are followed by high variety of other Steps, in contrast high outdegree reflects that these steps follow high variety of other Steps.

#### 3.2.3. Combination and order of Steps

Next, we examined usage patterns of specific Steps in relation to other Steps. For this, we refer to the results of the *Combination* and *Order* subsections of the *Steps* tab in the METEOR app. The *Combination* section provides a visualization of the frequency with which two different Steps are used together. For example, we illustrate the use of global signal regression (GlobalSigRegress) as a reference Step, as shown in Figure 4A. The plot indicates that more than 50 studies performed global signal regression with other motion and confound regression. This is an interesting observation, given that the use of global signal regression combined with other confound regression techniques is highly debated (Parkes et al., 2018).

**Figure 4.**
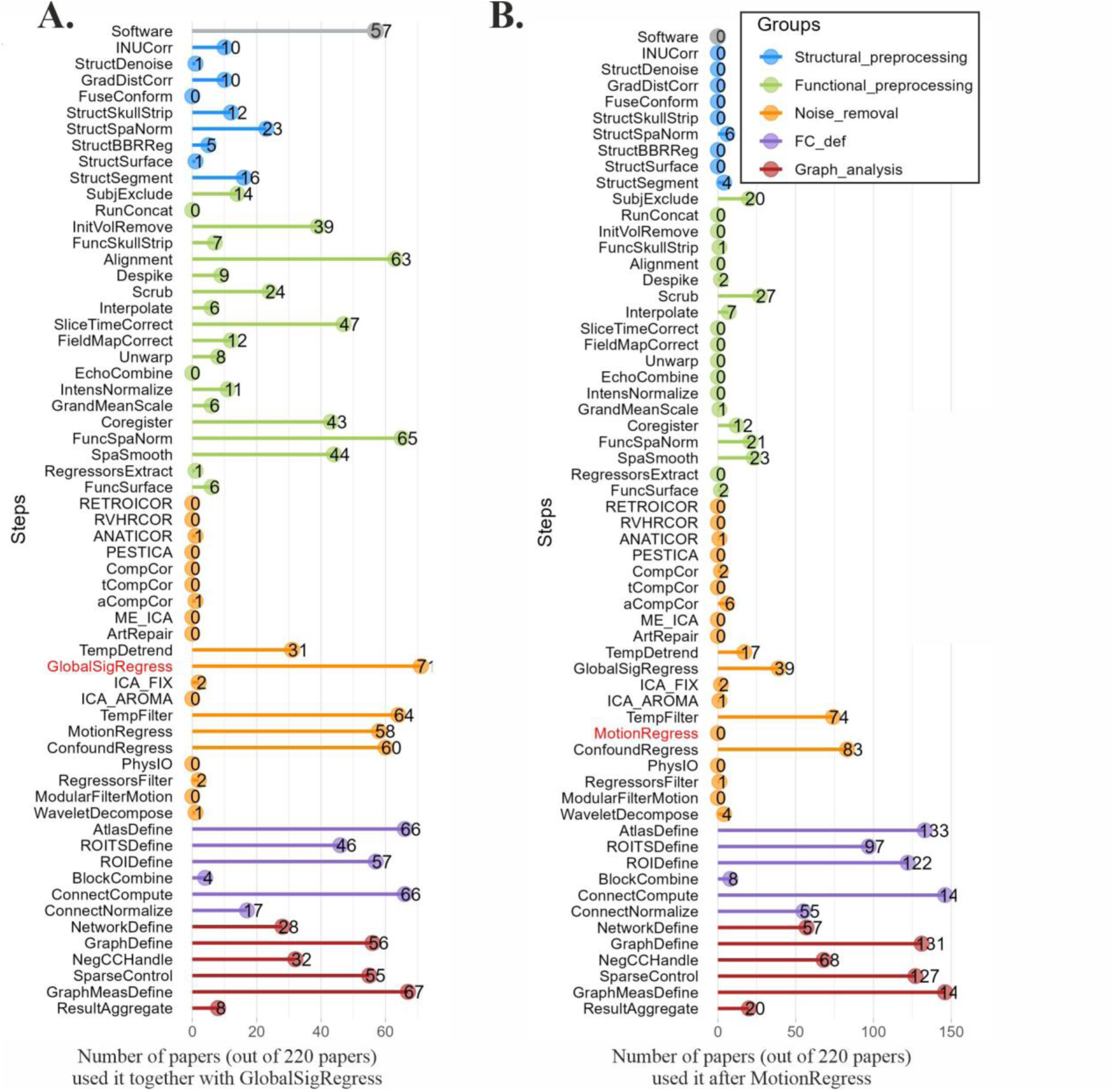
The occurrence of Steps relative to other Steps. Panel (A) visualizes the distribution of Steps which were used in combination with global signal regression (GlobalSigRegress, red label). It shows that there are many studies used global signal regression in combination with motion regression and/or other confound regression. The number of papers of the selected step indicates how many papers used this step. Panel (B) shows the Steps performed after implementing motion regression (MotionRegress, red label). The figure suggests that motion regression is typically performed at the end of the fMRI time-series extraction (low number of papers used any further step after motion regression) and before segmentation using the brain atlas. The number of papers of the selected step will always be zero.

The *Order* subsection shows the ordering of a particular Step relative to others. As an example, we examine the order of the motion regression Step (Figure 4B). The figure suggests that motion regression is typically performed at the end of the fMRI time-series extraction (low number of papers used any Steps after motion regression) and before segmentation using the brain atlas. Further exploration is possible via the METEOR app, using the *Combination* and *Order* tabs for all Steps extracted from the literature (C. Chang & Glover, 2009; Murphy & Fox, 2017; Power et al., 2014).

### 3.3. Variability in Options

#### 3.3.1. Variability in the Options of software and brain atlas

Out of the 61 Steps identified, we selected 17 Steps with the highest variability in the reported options. Figure 5 provides an overview of the distribution of Options for functional image preprocessing software and the brain atlases used to define regions of interest. As with the Steps, the distributions of Options for all Steps can also be accessed via the METEOR app that accompanies this paper. We also note that for all Steps with multiple Options, there are the Options of *not used* and *not reported*. The former refers to studies that did not mention the implementation of the corresponding Step, while the latter refers to studies that acknowledged the implementation of the Step but did not specify the Option chosen.

**Figure 5.**
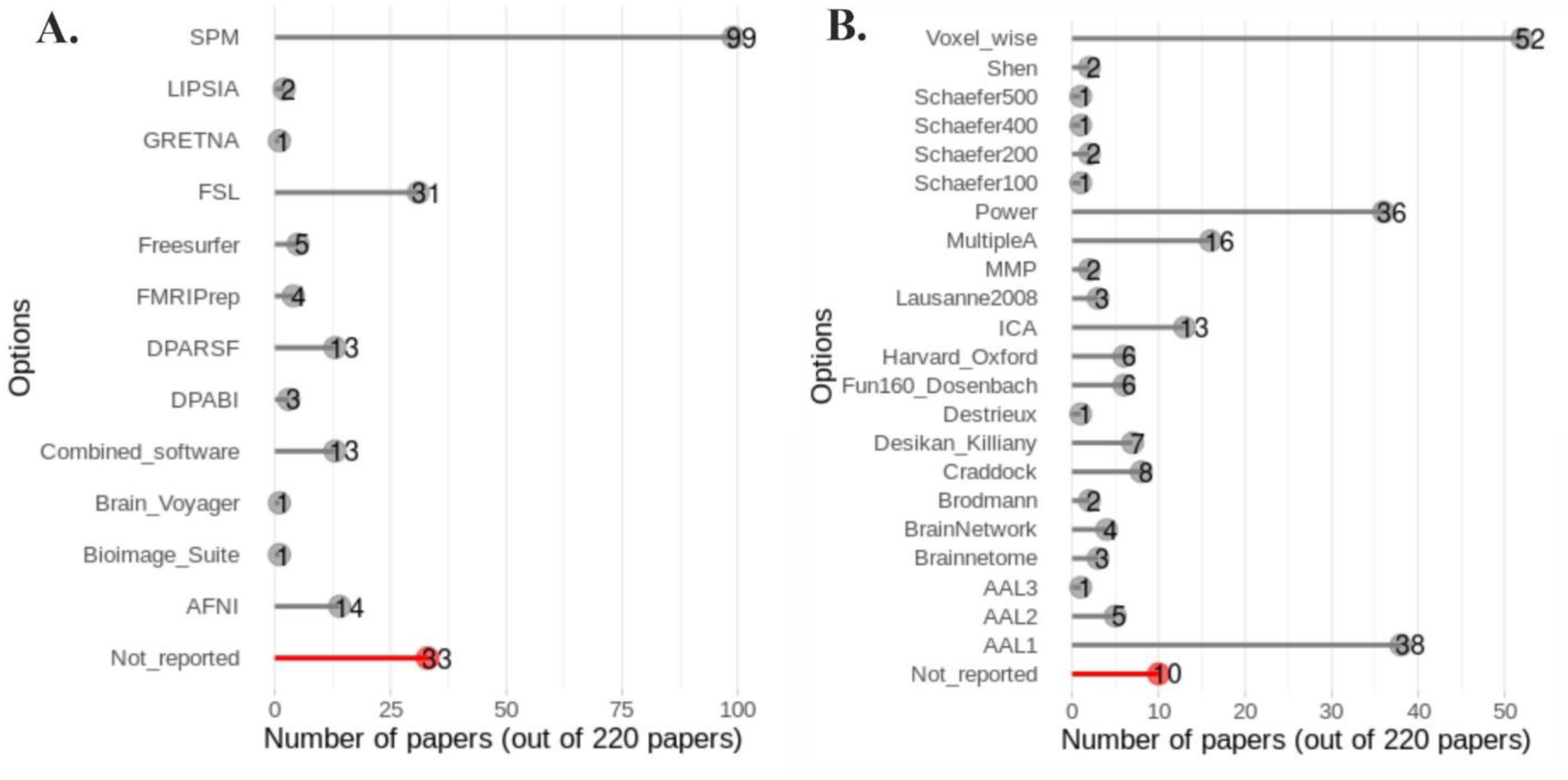
Distribution of Options. The distribution of Options in (A) the software and (B) the atlas definition Step. It is found that the software package SPM is the most popular software for fMRI data preprocessing. Moreover, it is noteworthy that a significant number of studies preferred voxel-based analysis by focusing on certain brain areas to region-based analysis.

Specifically, regarding the software used to preprocess the fMRI images, most studies used the software package SPM. The recently proposed toolbox to increase the reproducibility of fMRI preprocessing, fMRIPrep, was used by four studies identified as eligible for the present systematic review. Regarding the definition of brain atlases, numerous options were used, with the AAL16 (Automated Anatomical Labelling) and Power atlases being the most commonly used. However, it is noteworthy that a significant number of studies preferred voxel-based analysis (by focusing on certain areas) to region-based analysis (by focusing on the whole brain). Those studies were mostly interested in a particular brain area/subnetwork, thus, only limited number of voxels were included.

#### 3.3.2. Missing reports on the analysis Options applied

We selected 17 Steps where the Options are important and play a crucial role in the preprocessing and analysis of the data. However, a subset of studies did not report the Options they used, even though they indicated the use of these Steps. Figure 6 shows the number of studies that did not explicitly report Options for their respective Steps. The Step with the highest number of studies that did not report the corresponding Option was the alignment or motion correction of functional images. In addition, a number of studies did not report their chosen space for normalizing the functional images. Furthermore, for some Steps related to noise removal, there were cases where studies did not report their Options, such as the specific type of signal used for temporal detrending, the type of filter used for temporal filtering, and the number of parameters used in motion regression.

**Figure 6.**
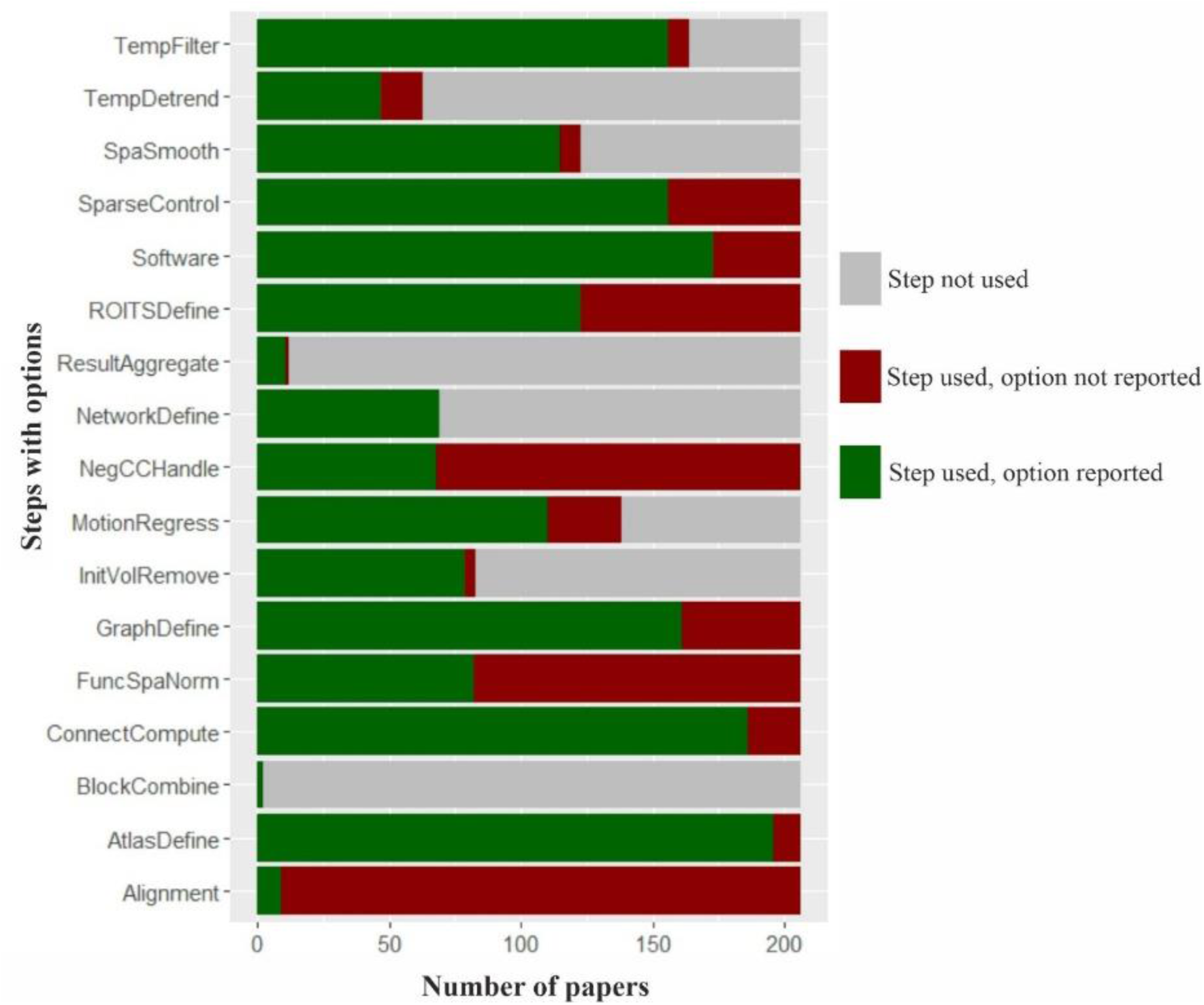
Variability in reporting the Options for fMRI preprocessing and analysis Steps. The plot visualizes the absolute frequency of the reported (green) and non-reported (red) Options. Many articles report that a particular Step in the analysis pipeline has been used without specifying the Option applied. It also shows that some studies did not perform a particular Step at all (grey).

## 4. Discussion

### 4.1. The multiverse

The present systematic review identified the multiverse of analytical decisions in the fMRI data preprocessing and analysis pipeline, with a particular focus on the emerging field of network neuroscience. Given concerns about the replicability of findings in this field of research, a clear description of the multiverse can provide valuable insights into defensible ways to assess robustness across analytical choices – a prerequisite for enhancing replicability.

We identified 61 distinct Steps employed in the preprocessing and analysis of fMRI data, 17 of which can vary significantly in terms of the main Options they can take. We refer to these Steps and Options as the ingredients of the multiverse. Researchers have considerable flexibility in constructing their unique pipelines by choosing a particular combination of these ingredients. Considering the vast array of ingredients, the multiverse of fMRI data preprocessing and analysis encompasses an extremely large number of forking paths, estimated to be in the trillions (i.e., by multiplying the number of Options, without even considering the exclusion and order of the Steps). To summarize, we established three taxonomic levels for characterizing the forking paths within the fMRI data preprocessing and analysis multiverse. These categorizations encompass: 1) inclusion or exclusion of specific Steps, 2) the sequence of executing Steps, and 3) parameter tuning (Options) of individual Steps. The first level only involves a single Step, occurring when researchers decided to use or not to use a particular Step in their analysis pipeline (e.g. global signal regression as one of the most debatable steps). Multiple Steps involved in the second level where the order of those Steps are altered. For instance, some studies perform temporal noise reduction at the voxel level prior to parcellation, while others prefer to implement it after brain region identification. The last level emerges when multiple Options are available for a particular Step, such as the choices of brain atlases for parcellation. It is important to note that these taxonomic levels do not necessarily reflect the relative influence of these forking paths on the final analysis results. In other words, the presence of a forking path at level 1 does not guarantee a lower impact on the findings compared to a path at level 3, and vice versa. A more comprehensive analysis, such as a sensitivity analysis, is required to assess the precise influence of these forking paths.

Another point worth to note is that the Steps specifically associated with the implementation of network science (i.e., computing functional connectivity (ConnectCompute), defining the graph edges (GraphDefine), defining the subnetworks (NetworkDefine), handling negative edges (NegCCHandle), controlling network sparsity (SparseControl), and aggregating results from different sparsity thresholds (ResultAggregate), account for a substantial proportion, approximately 30%, of all Options. The rapidly increasing application of network science in fMRI analysis stems from its capacity to investigate the complex association between brain function and behavior. This growing interest has expanded the number of options available in fMRI analysis pipelines. Further, the ongoing development of network science itself is likely to introduce even more options for graph-based fMRI data analysis (Srivastava et al., 2022). This expansion of available options reflects the field’s continuous development, and it does not necessarily imply a compromise in methodological rigor. Instead, it highlights the dynamic nature of fMRI analysis development and their potential for further advancements in understanding the functional connectivity patterns of the brain.

While the garden of the forking paths offers researchers greater flexibility and adaptability, it simultaneously poses challenges in terms of data analysis efficiency and reproducibility. Analyzing such an extensive number of paths is neither practical nor feasible. Furthermore, the results of this study show the analytical decisions and paths used in the literature to preprocess fMRI data, not the pipelines that should be used. The finding that there was no common pipeline used by more than one study also shows the considerable diversity in the pipeline and that there is no gold standard for preprocessing and parameterizing fMRI data for network neuroscience studies.

### 4.2. Narrowing the space by reducing the number of forking paths for future principled multiverse analysis

The very large number of potential forking paths raises an intriguing point for further exploration: How to reduce the number of potential pipelines for multiverse analysis applications, given the large number of options identified in this systematic review? The large number of analysis paths cannot be fully taken into account in a multiverse analysis due to the computational effort involved. A straightforward method may be to randomly sample from the entire multiverse extracted from the literature in the present work. However, this approach has limitations, primarily in terms of potentially limited coverage. Another approach was proposed with fMRIprep, an analysis-agnostic tool that automatically adapts the dataset to ensure optimal and high-quality preprocessing step (Esteban et al., 2019). However, finding an optimal or best pipeline to preprocess the data is not the main aim of multiverse analysis and identifying a best pipeline requires cross-validation approach in different samples or datasets.

Notably, the multiverse of data preprocessing and analysis decisions extracted here, may include non-equivalent decisions, i.e. one decision may be clearly better than another. However, this knowledge is not yet available in the knowledge space created by our systematic review and summarized in the METEOR app. A refined approach to multiverse analysis, called knowledge-based sampling, involves selecting forking paths based on their equivalence according to specific equivalence criteria. Del Giudice & Gangestad (2021) proposed a conceptual framework for assessing the equivalence of decisions/forking paths within the multiverse, emphasizing that only equivalent, and thus equally defensible analytical choices should be considered in a joint multiverse analysis. This assessment encompasses three aspects: measurement equivalence, effect equivalence, and power/precision equivalence. The outcome of this evaluation could determine whether a given forking path should be considered equivalent (principled equivalence), not equivalent (principled nonequivalence), or uncertain as compared with other steps and options in the path.

Notably, there is a challenge in deciding on the defensibility of analytical choices for fMRI graph theory studies according to the framework proposed by Del Giudice & Gangestad (2021). Given the highly complex data analysis procedure, no expert or novice in the field of behavioral neuroscience would have all the necessary knowledge to decide on the defensibility of analytical choices according to the criteria elaborated above. Our systematic review, together with the METEOR app, is a first step towards building up a systematic knowledge space covering defensible analytical solutions for network neuroscience approaches on brain-behavior associations. In the future, it may be better to describe analysis paths on a continuum as more or less comparable based on experts’ assessment supported by empirical studies. Moreover, the knowledge space will also need to contain information about how analytical choices affect insights into brain-behavior associations. Further, we envisage expansion of this knowledge space and automated updating of the space through literature mining and expertise crowdsourcing.

### 4.3. METEOR app as a decision support tool

This section aims to illustrate the utility of the interactive METEOR app in conducting multiverse studies, particularly in investigating debatable forking paths in fMRI data preprocessing and graph theory based analysis. It is important to emphasize that the METEOR app does not provide a definitive assessment of which forking path is superior to another; rather, it offers insights into the relative prevalence of these paths in the existing literature. Readers are highly encouraged to open the app and explore it alongside reading this discussion.

As a first example, let us focus on the question of combining ICA-AROMA with other regressions (i.e., motion regression, global signal regression, and white matter – cerebrospinal fluid (WM-CSF) signal regression). One study found that using only ICA-AROMA significantly improved resting-state network reproducibility (Pruim et al., 2015), while another study showed that incorporating ICA-AROMA with other regressions outperformed other approaches (using only motion regression and WM-CSF signal regression, and using only aCompCor) in terms of the residual relationship between in-scanner motion and functional connectivity in a high-motion dataset (Parkes et al., 2018). Using the METEOR app, we first examined the proportion of papers using ICA-AROMA in the *Steps* tab and the *Individual Step* subsection. We found 5 out of 220 studies. Next, in the *Combination* subsection, we noticed that none of these studies combined ICA-AROMA with global signal regression, while other studies combined ICA-AROMA with motion regression (three studies) and WM-CSF signal regression (four studies). Furthermore, a single study combined ICA-AROMA with ICA-FIX. In the *Orders* subsection, we can further investigate when in the analysis pipeline was ICA-AROMA performed. Motion regression and WM-CSF signal regression were mostly performed after ICA-AROMA, while one study that included ICA-FIX and ICA-AROMA decided to perform ICA-FIX before ICA-AROMA. In conclusion, we found that the majority of studies using ICA-AROMA still combined this step with other steps related to noise reduction, such as WM-CSF signal regression and ICA-AROMA, following the finding from Parkes et al., (2018).

Another example of potential exploration using the METEOR app is the use of Steps, which are intended to reduce noise but have been found to contribute to information loss, namely scrubbing (Aurich et al., 2015), spatial smoothing (Alahmadi, 2021), and global signal regression (Murphy & Fox, 2017). In the *Individual Step* subsection, we found that out of 220 papers, 67 used scrubbing, 134 used spatial smoothing, and 68 papers performed global signal regression. The combination of global signal regression and confound regression (motion and WM-CSF) is also an open discussion (Braun et al., 2012). In the *Combination* subsection of the *Steps* tab, we found that more than 75% of the papers that used global signal regression (68 papers) combined it with motion and WM-CSF signal regression. Finally, one can also explore the order of specific Steps, such as slice timing correction and motion regression, which have also been discussed previously (Parker & Razlighi, 2019). In the *Order* subsection, we found that all papers that used both steps (106 papers) performed slice timing correction before motion regression. Notably, previous study found that the optimal order between slice timing correction and motion regression depends on the level of motion and software used to perform fMRI preprocessing (Parker & Razlighi, 2019).

Similarly, the METEOR app can be used to explore the variability of the options, and the one that was commonly used, in the preprocessing steps and facilitate a more informed discussion. The above examples of the variability of the Options in the software and parcellation have also been discussed previously by Bowring et al. (2019) and Franco (2022). As an illustration, we examined the variability of Options in the size of the kernel in spatial smoothing, which was also addressed in a previous study by Alahmadi (2021). In the *Steps: Options* tab and by specifying the step as spatial smoothing, we found that the most common kernel size is 6 mm (52 papers), followed by 8 mm (42 papers) and 4 mm (18 papers). Another variability worth mentioning is the number of parameters in the motion regression. Using METEOR app, we found that the most common option was 6-parameter motion regression (3 rotation and 3 translation parameters), which was used in 53 papers that were selected for the present systematic review. However, a previous study by Satterthwaite et al. (2013) evaluated the effect of different numbers of motion regression parameters on the functional connectivity and showed evidence for less effect of motion on functional connectivity with higher number of parameters in motion regression (36-parameter).

Finally, the METEOR app can be used to design an own pipeline and compare it with the choices from the literature that have been fed into the current version of this knowledge space. As mentioned above, the user has the option to make the order of the steps relevant or irrelevant for this exploration. Especially for novices, such as PhD students, this option can be an invaluable learning aid.

### 4.4. The need of further standardization in reporting practices

This literature review also highlighted the continuing problem of some studies neglecting to report key decisions made during their analysis. Additionally, there were inconsistencies in reporting strategies across studies, such as using different terms to refer to identical steps or options, and a lack of uniformity in the level of detail provided regarding analytical decisions. These findings underscore the significance of implementing a comprehensive and standardized fMRI data preprocessing and analysis reporting template. A notable precedent for such an initiative is the “Guideline on How to Report an fMRI Study” proposed by Poldrack and colleagues in 2008 (Poldrack et al., 2008). Nevertheless, it is essential to note that this guideline lacks a specific template for reporting fMRI data preprocessing and analysis. An example of an existing reporting template that could serve as a model for this purpose is the “Agreed Reporting Template for EEG Methodology - International Standard (ARTEM-IS)” used for EEG studies (Kovic et al., 2023). Such a reporting template, that is also machine readable, would not only improve the clarity and transparency of the publications, but also contribute to the development of a dynamic database for the METEOR app accompanying this study and in general for the envisioned knowledge space discussed above. This dynamic database could be automatically updated with the latest information from recently published papers, enhancing the potential of the app as a valuable tool to support and facilitate further research efforts in the field and a better informed, more cumulative and robust network cognitive neuroscience.

### 4.5. Limitations

Our primary objective was to identify the multiverse in fMRI data preprocessing and analysis decisions, focusing specifically on network neuroscience. Consequently, the scope of this study was confined to graph-based analysis of fMRI data to explore the relationship between brain connectivity and behavior. However, by limiting the analysis to this specific domain, we may have overlooked potential analytical decisions obtained from other domains of fMRI analysis, for example the experimental setting and scanner parameters. We also limited our review to healthy individuals and the implementation of network science in fMRI data analysis. More general fMRI studies utilizing activation maps rather than functional connectivity analysis or studies involving patients may have different sets of analytical choices. Extending the analysis to these aspects would reveal an even more extensive multiverse. While the application we developed for this study is exclusively founded on the dataset derived from the literature review performed in this study, the functionality of the METEOR app could be easily expanded and the database continuously updated, possibly in an automated way, to incorporate other aspects of fMRI analysis not considered here.

### 4.6. Conclusion and outlook

This comprehensive literature review has identified the multitude of forking paths in graph-fMRI data preprocessing and analysis pipelines. Employing all possible combinations of these forking paths would be impractical and computationally intractable. To address this challenge, further research should focus on developing strategies for identifying defensible forking paths based on empirical studies, therefore reducing the computational burden associated with multiverse analysis. Additionally, the ongoing advancement of network science, with its growing array of methodological approaches, requires the consideration of potential forking paths that may emerge with future developments. While machine learning techniques could potentially offer computational assistance in conducting multiverse analysis in high-dimensional spaces (Dafflon et al., 2022), it is essential to recognize that these methods introduce their own set of forking paths due to the flexibility of model specification. Therefore, further studies are needed to carefully evaluate the trade-off between the flexibility of machine learning models and their utility in facilitating multiverse analysis. Finally, transparency remains important in mitigating the replication crisis. A lack of transparency in multiverse analysis could potentially introduce new sources of bias and reproducibility issues. Therefore, researchers must strive to maintain transparency throughout the multiverse analysis process, ensuring that all decisions and parameter choices are clearly documented and justified.

## Declaration of Interest

Declaration of interest: None.

## Acknowledgment

This work was supported by a grant from the German Research Foundation (DFG) to Andrea Hildebrandt (HI 1780/7-1), Carsten Gießing (GI 682/5-1), Stefan Debener (DE 779/8-1) and Christiane Thiel (TH 766/9-1) as part of the DFG priority program “META-REP: A Meta-scientific Programme to Analyse and Optimise Replicability in the Behavioural, Social, and Cognitive Sciences” (SPP 2317).

## Notes

https://meteor-oldenburg.shinyapps.io/fMRI_multiverse/

https://github.com/kristantodan12/fMRI_Multiverse/

### Competing Interest Statement

The authors have declared no competing interest.

