## Supplementary Table S1 for "The multiverse of data preprocessing and analysis in graph-based fMRI: A systematic literature review of analytical choices fed into a decision support tool for informed analysis"

### Supplementary Materials

Table S1. List of Identified Steps and Options. The last column indicates the studies used the Step or the Option.

| Name in METEOR app | Options | Articles |
|---|---|---|
| Software | SPM | (Shen et al., 2013); (Y. Wang et al., 2021); (Quante et al., 2018); (Zheng et al., 2021); (C. Yan & He, 2011); (Hayasaka, 2013); (G. Zhang & Liu, 2021); (X. Liu et al., 2020); (Gozdas et al., 2019); (Markett et al., 2018); (Tian et al., 2011); (L. Song et al., 2020); (Feng et al., 2015); (Le et al., 2020); (J. Wang et al., 2017); (Pan et al., 2018); (Kawagoe et al., 2017); (F. Fan et al., 2021); (Servaas et al., 2017); (Xia et al., 2019); (Westphal et al., 2017); (He et al., 2019); (Pamplona et al., 2015); (Iordan et al., 2018); (S. Zhang et al., 2022); (Jarrahi & Kollias, 2020); (Geib et al., 2017); (Deng et al., 2016); (Belden et al., 2020); (Braun et al., 2012); (Ding et al., 2011); (Finc et al., 2017); (Vatansever et al., 2015a); (Liang et al., 2012); (Geerligs et al., 2014); (Duan et al., 2014); (Farah & Horowitz-Kraus, 2019); (Neyland et al., 2021); (Meunier et al., 2014); (Alluri et al., 2017); (Breckel et al., 2013); (H. Zhang et al., 2012); (Santarnecchi et al., 2014); (Y. Fan et al., 2019); (Geerligs et al., 2015); (Alavash, Doebler, et al., 2015); (Markett et al., 2013); (Wu et al., 2013); (Smith et al., 2018); (P. Xu et al., 2014); (J. Liu et al., 2017); (Rubin et al., 2017); (Vatansever et al., 2015b); (Qin et al., 2016); (Xiao et al., 2016); (C. Wang et al., 2022); (Sreenivasan et al., 2017); (Suprano et al., 2019); (Jung et al., 2018); (Sun et al., 2017); (Y. Gu et al., 2022); (Göttlich et al., 2017); (Borchardt et al., 2015); (L. Wang et al., 2010); (Burdette et al., 2010); (Jin et al., 2020); (Moorthigari et al., 2020); (Ebrahimi et al., 2019); (Xi et al., 2019); (Agrawal et al., 2019); (Cao et al., 2019); (Brandl et al., 2018); (Huang et al., 2018); (Reddy et al., 2018); (Aggarwal et al., 2017); (Y. Liu et al., 2017); (Berroir et al., 2017); (Prčkovska et al., 2016); (de Paula et al., 2017); (Zhao et al., 2017); (File et al., 2016); (Alavash et al., 2016); (Du et al., 2015); (Alavash, Hilgetag, et al., 2015); (Wen et al., 2015); (X. Xu et al., 2015); (Taya et al., 2014); (Arnold et al., 2014); (Sami & Miall, 2013); (Spreng et al., 2013); (Messé et al., 2012); (Franzmeier et al., 2018); (Martial et al., 2023); (Dan et al., 2023); (X. Chen et al., 2023); (Yang et al., 2023); (H. Wang et al., 2023); (Invernizzi et al., 2023); (Lee et al., 2022) |
| | AFNI | (Tooley et al., 2020); (J. Song et al., 2014); (Danti et al., 2018); (Chong et al., 2019); (Yue et al., 2017); (Cohen & D'Esposito, 2016); (Cole et al., 2015); (Sheppard et al., 2011); (Koba et al., 2021); (Ketchabaw et al., 2022); (Anderson et al., 2017); (Liang et al., 2016); (Liang et al., 2013); (Han et al., 2023) |
| | DPARSF | (Bartholomew et al., 2019); (X. Liu et al., 2018); (Q. Li et al., 2019); (H. Yan et al., 2022); (Bueichekú et al., 2019); (X. Li et al., 2020); (C. Wang et al., 2020); (D. Liu et al., 2022); (X. Zhang et al., 2015); (J. Zhang et al., 2021); (Shang et al., 2017); (Cocchi et al., 2015); (Y. Gao et al., 2023) |
| | DPABI | (Zhou et al., 2021); (Farahani et al., 2022); (J. Lin et al., 2022) |
| | FSL | (Gracia-Tabuenca et al., 2021); (Foo et al., 2021); (Masuda et al., 2018); (Vriend et al., 2020); (Varangis et al., 2021); (Setton et al., 2022); (Madden et al., 2020); (Pezoulas et al., 2017); (Hilger et al., 2017b); (Bailey et al., 2018); (Monge et al., 2017); (Pindus et al., 2020); (Schlesinger et al., 2017); (Ray et al., 2020); (Orwig et al., 2021); (Lunsford-Avery et al., 2020); (Reineberg & Banich, 2016); (Kolskår et al., 2018); (Ginestet & Simmons, 2011); (Polanía et al., 2011); (Alnæs et al., 2015); (Jacob et al., 2016); (Ekman et al., 2012); (Huskey et al., 2018); (Zamroziewicz et al., 2017); (de Pasquale et al., 2017); (Moussa et al., 2012); (Satterthwaite et al., 2012); (Richards et al., 2018); (Breedt et al., 2022); (Ghiles et al., 2023) |

| | | |
|---|---|---|
| | FMRIPrep | (Malagurski et al., 2020); (Evensmoen et al., 2021); (Finc et al., 2020); (Tooley et al., 2022) |
| | Brain_Voyager | (Parhizi et al., 2018) |
| | Bioimage_Suite | (S. Gao et al., 2021) |
| | GRETNA | (X. Fan et al., 2021) |
| | LIPSIA | (Taruffi et al., 2017); (Koelsch & Skouras, 2014) |
| | Freesurfer | (Bottino et al., 2021); (Manza et al., 2020); (DeSalvo et al., 2014); (Amlien et al., 2019); (Mancini et al., 2017) |
| | Combined_software | (Kobayashi et al., 2020); (Su et al., 2021); (Sato et al., 2015); (Sheppard et al., 2012); (Gopinath et al., 2015); (Hearne et al., 2017); (Marek et al., 2015); (Yi et al., 2023); (Y. Fan et al., 2021); (Betzel et al., 2020); (Hilger et al., 2017a); (S. Gu et al., 2015); (Sheppard et al., 2011) |
| | Not reported | (Fujimoto et al., 2020); (Fukushima et al., 2017); (Zuo et al., 2018); (Ogawa, 2021); (Kruschwitz et al., 2018); (Yin et al., 2019); (Stevens et al., 2012); (Varangis et al., 2019); (Fukushima & Sporns, 2018); (Long et al., 2017); (C. Zhang et al., 2016); (Liao et al., 2017); (Rzucidlo et al., 2013); (T. Chen et al., 2016); (Huckins et al., 2019); (Zhong et al., 2014); (Lloyd, 2020); (Beaty et al., 2015); (Spielberg et al., 2015); (Crossley et al., 2013); (Shine et al., 2016); (Y. Zhang et al., 2022); (Tipnis et al., 2020); (Kim et al., 2018); (Knyazeva et al., 2018); (Ma & Zhang, 2017); (Bolt et al., 2017); (Gratton et al., 2016); (Najafi et al., 2016); (Thompson & Fransson, 2015); (Choi et al., 2023); (Khodaei et al., 2023); (Ryu et al., 2022) |
| INUCorr | | (Malagurski et al., 2020); (Evensmoen et al., 2021); (Tooley et al., 2022); (Betzel et al., 2020); (Gracia-Tabuenca et al., 2021); (Amlien et al., 2019); (S. Gao et al., 2021); (Fujimoto et al., 2020); (Fukushima et al., 2017); (Zuo et al., 2018); (Ogawa, 2021); (Kruschwitz et al., 2018); (Yin et al., 2019); (Pezoulas et al., 2017); (Fukushima & Sporns, 2018); (C. Zhang et al., 2016); (Liao et al., 2017); (T. Chen et al., 2016); (Lloyd, 2020); (Manza et al., 2020); (Jacob et al., 2016); (Y. Zhang et al., 2022); (Tipnis et al., 2020); (Kim et al., 2018); (Ma & Zhang, 2017); (Bolt et al., 2017); (Najafi et al., 2016); (Thompson & Fransson, 2015); (Khodaei et al., 2023) |
| StructDenoise | | (Gracia-Tabuenca et al., 2021) |
| GradDistCorr | | S. Gao et al., 2021); (Fujimoto et al., 2020); (Fukushima et al., 2017); (Foo et al., 2021); (Zuo et al., 2018); (Ogawa, 2021); (Kruschwitz et al., 2018); (Yin et al., 2019); (Pezoulas et al., 2017); (Fukushima & Sporns, 2018); (C. Zhang et al., 2016); (Liao et al., 2017); (T. Chen et al., 2016); (Lloyd, 2020); (Manza et al., 2020); (Jacob et al., 2016); (Y. Zhang et al., 2022); (Tipnis et al., 2020); (Kim et al., 2018); (Ma & Zhang, 2017); (Bolt et al., 2017); (Najafi et al., 2016); (Thompson & Fransson, 2015); (Khodaei et al., 2023); (S. Gao et al., 2021); (Fujimoto et al., 2020); (Fukushima et al., 2017); (Foo et al., 2021); (Zuo et al., 2018); (Ogawa, 2021); (Kruschwitz et al., 2018); (Yin et al., 2019); (Pezoulas et al., 2017); (Fukushima & Sporns, 2018); (C. Zhang et al., 2016); (Liao et al., 2017); (T. Chen et al., 2016); (Lloyd, 2020); (Manza et al., 2020); (Jacob et al., 2016); (Y. Zhang et al., 2022); (Tipnis et al., 2020); (Kim et al., 2018); (Ma & Zhang, 2017); (Bolt et al., 2017); (Najafi et al., 2016); (Thompson & Fransson, 2015); (Khodaei et al., 2023) |
| FuseConform | | |
| StructSkullStrip | | (Masuda et al., 2018); (Vriend et al., 2020); (Setton et al., 2022); (Long et al., 2017); (Monge et al., 2017); (Pindus et al., 2020); (Neyland et al., 2021); (Koba et al., 2021); (Orwig et al., 2021); (Zamroziewicz et al., 2017); (S. Gao et al., 2021); (Fujimoto et al., 2020); (Fukushima et al., 2017); (Foo et al., 2021); (Zuo et al., 2018); (Ogawa, |

| | | |
|---|---|---|
| | | 2021); (Kruschwitz et al., 2018); (Yin et al., 2019); (Malagurski et al., 2020); (Evensmoen et al., 2021); (Pezoulas et al., 2017); (Fukushima & Sporns, 2018); (C. Zhang et al., 2016); (Liao et al., 2017); (T. Chen et al., 2016); (Lloyd, 2020); (Hearne et al., 2017); (Tooley et al., 2022); (Manza et al., 2020); (Jacob et al., 2016); (Y. Zhang et al., 2022); (Betzel et al., 2020); (Tipnis et al., 2020); (Kim et al., 2018); (Ma & Zhang, 2017); (Bolt et al., 2017); (Najafi et al., 2016); (Thompson & Fransson, 2015); (Khodaei et al., 2023); (Gozdas et al., 2019); (Reineberg & Banich, 2016); (S. Gu et al., 2015); (Han et al., 2023); (J. Song et al., 2014); (Sato et al., 2015) |
| StructSpaNorm | | (Tooley et al., 2020); (Invernizzi et al., 2023); (Foo et al., 2021); (Malagurski et al., 2020); (Long et al., 2017); (Neyland et al., 2021); (Orwig et al., 2021); (Moorthigari et al., 2020); (Reddy et al., 2018); (Moussa et al., 2012); (Gozdas et al., 2019); (Feng et al., 2015); (Le et al., 2020); (X. Liu et al., 2018); (Finc et al., 2017); (Alluri et al., 2017); (Koba et al., 2021); (Crossley et al., 2013); (J. Zhang et al., 2021); (Betzel et al., 2020); (Amlien et al., 2019); (Huang et al., 2018); (Y. Liu et al., 2017); (de Paula et al., 2017); (File et al., 2016); (Quante et al., 2018); (Su et al., 2021); (J. Song et al., 2014); (Evensmoen et al., 2021); (J. Wang et al., 2017); (Yue et al., 2017); (Westphal et al., 2017); (Pamplona et al., 2015); (Iordan et al., 2018); (Vatansever et al., 2015b); (Hearne et al., 2017); (C. Wang et al., 2020); (Göttlich et al., 2017); (Borchardt et al., 2015); (Lin et al., 2022); (Berroir et al., 2017); (Gratton et al., 2016); (Franzmeier et al., 2018); (Pan et al., 2018); (S. Zhang et al., 2022); (Jarrahi & Kollias, 2020); (Deng et al., 2016); (Xiao et al., 2016); (C. Wang et al., 2022); (Ebrahimi et al., 2019); (Brandl et al., 2018); (Du et al., 2015); (Martial et al., 2023); (Dan et al., 2023); (Suprano et al., 2019); (Jung et al., 2018); (Y. Gu et al., 2022); (Jin et al., 2020); (de Pasquale et al., 2017); (Mancini et al., 2017); (X. Xu et al., 2015); (Han et al., 2023); (Varangis et al., 2019); (F. Fan et al., 2021); (Schlesinger et al., 2017); (Belden et al., 2020); (Y. Fan et al., 2019); (Reineberg & Banich, 2016); (Shang et al., 2017); (Liang et al., 2016); (S. Gu et al., 2015); (Liang et al., 2013); (Richards et al., 2018); (Ketchabaw et al., 2022); (Y. Gao et al., 2023); (Pindus et al., 2020); (Santarnecchi et al., 2014); (Zamroziewicz et al., 2017); (Hilger et al., 2017a); (Sato et al., 2015); (Satterthwaite et al., 2012) |
| StructBBRReg | | S. Gao et al., 2021); (Fujimoto et al., 2020); (Fukushima et al., 2017); (Zuo et al., 2018); (Ogawa, 2021); (Kruschwitz et al., 2018); (Yin et al., 2019); (Pezoulas et al., 2017); (Fukushima & Sporns, 2018); (C. Zhang et al., 2016); (Liao et al., 2017); (T. Chen et al., 2016); (Lloyd, 2020); (Manza et al., 2020); (Jacob et al., 2016); (Y. Zhang et al., 2022); (Tipnis et al., 2020); (Kim et al., 2018); (Ma & Zhang, 2017); (Bolt et al., 2017); (Najafi et al., 2016); (Thompson & Fransson, 2015); (Khodaei et al., 2023) |
| StructSurface | | (Evensmoen et al., 2021); (Tooley et al., 2022) |
| StructSegment | | (Xia et al., 2019); (Vriend et al., 2020); (Long et al., 2017); (Pindus et al., 2020); (Neyland et al., 2021); (Alluri et al., 2017); (Koba et al., 2021); (Orwig et al., 2021); (Moorthigari et al., 2020); (Zamroziewicz et al., 2017); (Moussa et al., 2012); (Invernizzi et al., 2023); (Su et al., 2021); (Tooley et al., 2022); (Marek et al., 2015); (Betzel et al., 2020); (Amlien et al., 2019); (de Paula et al., 2017); (File et al., 2016); (Foo et al., 2021); (Quante et al., 2018); (Malagurski et al., 2020); (Evensmoen et al., 2021); (Iordan et al., 2018); (Reineberg & Banich, 2016); (Vatansever et al., 2015b); (Hearne et al., 2017); (Polanía et al., 2011); (Borchardt et al., 2015); (Arnold et al., 2014); (C. Yan & He, 2011); (Gozdas et al., 2019); (Feng et al., 2015); (Le et al., 2020); (Cohen & D'Esposito, 2016); (X. Liu et al., 2018); (S. Zhang et al., 2022); (Jarrahi & Kollias, 2020); (Deng et al., 2016); (Finc et al., 2017); (Xiao et al., 2016); (C. Wang et al., 2022); (Sun et al., 2017); (Brandl et al., 2018); (Martial et al., 2023); (Dan et al., 2023); (H. Wang et al., 2023); (Varangis et al., 2021); (Varangis et al., 2019); (J. Wang et al., 2017); (Westphal et al., 2017); (Pamplona et al., 2015); (Suprano et al., 2019); (Y. Gu et al., 2022); (Jin et |

| | | |
|---|---|---|
| | | al., 2020); (Mancini et al., 2017); (X. Xu et al., 2015); (Pan et al., 2018); (F. Fan et al., 2021); (Y. Fan et al., 2019); (Du et al., 2015); (Richards et al., 2018); (Y. Gao et al., 2023); (Kobayashi et al., 2020); (Masuda et al., 2018); (J. Song et al., 2014); (Markett et al., 2018); (Belden et al., 2020); (Santarnecchi et al., 2014); (Ketchabaw et al., 2022); (Smith et al., 2018) |
| SubjExclude | | (Gozdas et al., 2019); (Pamplona et al., 2015); (Wu et al., 2013); (Farahani et al., 2022); (Pan et al., 2018); (J. Liu et al., 2017); (Y. Gu et al., 2022); (Jin et al., 2020); (Du et al., 2015); (X. Chen et al., 2023); (Pindus et al., 2020); (Belden et al., 2020); (Santarnecchi et al., 2014); (Suprano et al., 2019); (Vriend et al., 2020); (Jung et al., 2018); (Finc et al., 2017); (Zhou et al., 2021); (D. Liu et al., 2022); (G. Zhang & Liu, 2021); (Marek et al., 2015); (Servaas et al., 2017); (Geerligs et al., 2015); (Finc et al., 2020); (Sun et al., 2017); (Hearne et al., 2017); (X. Zhang et al., 2015); (Foo et al., 2021); (Xia et al., 2019); (Liao et al., 2017); (Xiao et al., 2016); (Borchardt et al., 2015); (Tooley et al., 2020); (Markett et al., 2018); (Dan et al., 2023); (F. Fan et al., 2021); (Najafi et al., 2016); (S. Gao et al., 2021); (Y. Fan et al., 2019); (Tooley et al., 2022); (Fukushima et al., 2017) |
| RunConcat | | (Madden et al., 2020); (Danti et al., 2018) |
| InitVolRemove | 1V | Sheppard et al., 2011); (Sheppard et al., 2012) |
| | 2V | (Beaty et al., 2015); (Jung et al., 2018); (Polanía et al., 2011); (Brandl et al., 2018) |
| | 3V | (J. Song et al., 2014); (Xiao et al., 2016); (Amlien et al., 2019); (Han et al., 2023) |
| | 4V | (Tooley et al., 2020); (Vriend et al., 2020); (Setton et al., 2022); (Hilger et al., 2017b); (Monge et al., 2017); (Sato et al., 2015); (Reineberg & Banich, 2016); (Göttlich et al., 2017); (Yi et al., 2023); (Y. Fan et al., 2021); (Mancini et al., 2017); (Hilger et al., 2017a); (Liang et al., 2016); (S. Gu et al., 2015); (DeSalvo et al., 2014); (Liang et al., 2013); (Spreng et al., 2013); (Satterthwaite et al., 2012); (Breedt et al., 2022) |
| | 5V | (Feng et al., 2015); (J. Wang et al., 2017); (Kawagoe et al., 2017); (Taruffi et al., 2017); (Cole et al., 2015); (Duan et al., 2014); (H. Zhang et al., 2012); (Santarnecchi et al., 2014); (Y. Fan et al., 2019); (J. Liu et al., 2017); (C. Wang et al., 2020); (J. Zhang et al., 2021); (Betzel et al., 2020); (Ebrahimi et al., 2019); (Du et al., 2015); (X. Chen et al., 2023); (Yang et al., 2023) |
| | 6V | (Vatansever et al., 2015a); (C. Wang et al., 2022); (Y. Gu et al., 2022); (Ekman et al., 2012) |
| | 8V | (Qin et al., 2016) |
| | 9V | (Alavash et al., 2016); (Alavash, Hilgetag, et al., 2015) |
| | 10V | (Foo et al., 2021); (Y. Wang et al., 2021); (C. Yan & He, 2011); (Yin et al., 2019); (X. Liu et al., 2020); (X. Fan et al., 2021); (Markett et al., 2018); (Tian et al., 2011); (Farahani et al., 2022); (L. Song et al., 2020); (Bartholomew et al., 2019); (Fukushima & Sporns, 2018); (Pan et al., 2018); (Long et al., 2017); (F. Fan et al., 2021); (Chong et al., 2019); (Xia et al., 2019); (He et al., 2019); (S. Zhang et al., 2022); (Deng et al., 2016); (Liang et al., 2012); (Q. Li et al., 2019); (Neyland et al., 2021); (Markett et al., 2013); (P. Xu et al., 2014); (X. Li et al., 2020); (Sun et al., 2017); (D. Liu et al., 2022); (Borchardt et al., 2015); (Shine et al., 2016); (Lin et al., 2022); (X. Zhang et al., 2015); (Jin et al., 2020); (Aggarwal et al., 2017); (Shang et al., 2017); (Wen et al., 2015); (Cocchi et al., 2015); (H. Wang et al., 2023) |
| | Not reported | (Fukushima et al., 2017); (Madden et al., 2020); (Wu et al., 2013); (X. Xu et al., 2015) |

| | | |
|---|---|---|
| FuncSkullStrip | | (Huskey et al., 2018); (Satterthwaite et al., 2012); (Breedt et al., 2022); (Setton et al., 2022); (Schlesinger et al., 2017); (Bottino et al., 2021); (Sato et al., 2015); (Lunsford-Avery et al., 2020); (Cohen & D'Esposito, 2016); (Zhong et al., 2014); (Ghiles et al., 2023); (Kolskår et al., 2018); (Gracia-Tabuenca et al., 2021); (Reineberg & Banich, 2016); (Alnæs et al., 2015); (Tooley et al., 2020); (Madden et al., 2020); (Betzel et al., 2020) |
| Alignment | First | (Tian et al., 2011); (L. Song et al., 2020); (Feng et al., 2015); (Pamplona et al., 2015); (Rzucidlo et al., 2013); (Geib et al., 2017); (Ding et al., 2011); (Bottino et al., 2021); (Q. Li et al., 2019); (Bueichekú et al., 2019); (Alavash, Doebler, et al., 2015); (Ginestet & Simmons, 2011); (Sun et al., 2017); (Y. Gu et al., 2022); (Ebrahimi et al., 2019); (Alavash et al., 2016); (Alavash, Hilgetag, et al., 2015); (Martial et al., 2023) |
| | Median | (Satterthwaite et al., 2012); (Ryu et al., 2022) |
| | Mean | (Gozdas et al., 2019); (Finc et al., 2017); (Santarnecchi et al., 2014) |
| | Not reported | (Shen et al., 2013); (S. Gao et al., 2021); (Fujimoto et al., 2020); (Kobayashi et al., 2020); (Fukushima et al., 2017); (Gracia-Tabuenca et al., 2021); (Foo et al., 2021); (Y. Wang et al., 2021); (Quante et al., 2018); (Su et al., 2021); (Tooley et al., 2020); (Zhou et al., 2021); (Zheng et al., 2021); (C. Yan & He, 2011); (Zuo et al., 2018); (Masuda et al., 2018); (Vriend et al., 2020); (Varangis et al., 2021); (Ogawa, 2021); (Hayasaka, 2013); (Kruschwitz et al., 2018); (Yin et al., 2019); (G. Zhang & Liu, 2021); (X. Liu et al., 2020); (X. Fan et al., 2021); (Malagurski et al., 2020); (J. Song et al., 2014); (Setton et al., 2022); (Madden et al., 2020); (Evensmoen et al., 2021); (Pezoulas et al., 2017); (Parhizi et al., 2018); (Stevens et al., 2012); (Markett et al., 2018); (Farahani et al., 2022); (Hilger et al., 2017b); (Varangis et al., 2019); (Bartholomew et al., 2019); (Danti et al., 2018); (Bailey et al., 2018); (Le et al., 2020); (Fukushima & Sporns, 2018); (J. Wang et al., 2017); (Pan et al., 2018); (Long et al., 2017); (Kawagoe et al., 2017); (F. Fan et al., 2021); (Servaas et al., 2017); (Chong et al., 2019); (Yue et al., 2017); (Cohen & D'Esposito, 2016); (Monge et al., 2017); (Xia et al., 2019); (X. Liu et al., 2018); (C. Zhang et al., 2016); (Liao et al., 2017); (Westphal et al., 2017); (He et al., 2019); (Pindus et al., 2020); (Iordan et al., 2018); (S. Zhang et al., 2022); (Jarrahi & Kollias, 2020); (Schlesinger et al., 2017); (T. Chen et al., 2016); (Deng et al., 2016); (Taruffi et al., 2017); (Belden et al., 2020); (Huckins et al., 2019); (Braun et al., 2012); (Cole et al., 2015); (Zhong et al., 2014); (Vatansever et al., 2015a); (Sheppard et al., 2011); (Lloyd, 2020); (Liang et al., 2012); (Geerligs et al., 2014); (Beaty et al., 2015); (Duan et al., 2014); (Farah & Horowitz-Kraus, 2019); (Neyland et al., 2021); (Meunier et al., 2014); (H. Yan et al., 2022); (Alluri et al., 2017); (Koba et al., 2021); (Ray et al., 2020); (Breckel et al., 2013); (H. Zhang et al., 2012); (Sato et al., 2015); (Y. Fan et al., 2019); (Sheppard et al., 2012); (Orwig et al., 2021); (Gopinath et al., 2015); (Lunsford-Avery et al., 2020); (Geerligs et al., 2015); (Reineberg & Banich, 2016); (Markett et al., 2013); (Wu et al., 2013); (Smith et al., 2018); (P. Xu et al., 2014); (J. Liu et al., 2017); (Kolskår et al., 2018); (Rubin et al., 2017); (Vatansever et al., 2015b); (Qin et al., 2016); (Xiao et al., 2016); (C. Wang et al., 2022); (Hearne et al., 2017); (Sreenivasan et al., 2017); (Suprano et al., 2019); (X. Li et al., 2020); (Jung et al., 2018); (Finc et al., 2020); (Spielberg et al., 2015); (Tooley et al., 2022); (Polanía et al., 2011); (Marek et al., 2015); (Crossley et al., 2013); (Manza et al., 2020); (Alnæs et al., 2015); (C. Wang et al., 2020); (D. Liu et al., 2022); (Jacob et al., 2016); (Ketchabaw et al., 2022); (Göttlich et al., 2017); (Borchardt et al., 2015); (Ekman et al., 2012); (L. Wang et al., 2010); (Burdette et al., 2010); (Shine et al., 2016); (Lin et al., 2022); (X. Zhang et al., 2015); (Y. Zhang et al., 2022); (Yi et al., 2023); (J. Zhang et al., 2021); (Y. Fan et al., 2021); (Betzel et al., 2020); (Jin et al., 2020); (Moorthigari et al., 2020); (Tipnis et al., 2020); (Amlien et al., 2019); (Xi et al., 2019); (Agrawal et al., 2019); (Cao et al., 2019); (Huskey et al., 2018); (Brandl et al., 2018); (Kim et al., 2018); (Huang et al., 2018); (Reddy et al., 2018); (Knyazeva et al., 2018); (Aggarwal et al., 2017); |

| | | |
|---|---|---|
| | | (Zamroziewicz et al., 2017); (de Pasquale et al., 2017); (Ma & Zhang, 2017); (Y. Liu et al., 2017); (Berroir et al., 2017); (Mancini et al., 2017); (Bolt et al., 2017); (Prčkovska et al., 2016); (de Paula et al., 2017); (Anderson et al., 2017); (Shang et al., 2017); (Zhao et al., 2017); (Hilger et al., 2017a); (Gratton et al., 2016); (File et al., 2016); (Najafi et al., 2016); (Liang et al., 2016); (S. Gu et al., 2015); (Thompson & Fransson, 2015); (Du et al., 2015); (Wen et al., 2015); (X. Xu et al., 2015); (Cocchi et al., 2015); (Taya et al., 2014); (DeSalvo et al., 2014); (Arnold et al., 2014); (Koelsch & Skouras, 2014); (Sami & Miall, 2013); (Liang et al., 2013); (Spreng et al., 2013); (Moussa et al., 2012); (Messé et al., 2012); (Sheppard et al., 2011); (Franzmeier et al., 2018); (Richards et al., 2018); (Dan et al., 2023); (Breedt et al., 2022); (Choi et al., 2023); (X. Chen et al., 2023); (Han et al., 2023); (Y. Gao et al., 2023); (Khodaei et al., 2023); (Ghiles et al., 2023); (Yang et al., 2023); (H. Wang et al., 2023); (Invernizzi et al., 2023); (Lee et al., 2022) |
| Despike | | J. Song et al., 2014); (Stevens et al., 2012); (Yue et al., 2017); (Marek et al., 2015); (Mancini et al., 2017); (Han et al., 2023); (Spielberg et al., 2015); (Ketchabaw et al., 2022); (Richards et al., 2018); (Hilger et al., 2017b); (Chong et al., 2019); (Sato et al., 2015); (Hilger et al., 2017a); (He et al., 2019); (Xi et al., 2019); (Quante et al., 2018); (Long et al., 2017); (Le et al., 2020); (Bolt et al., 2017) |
| Scrub | | (Varangis et al., 2021); (Varangis et al., 2019); (Qin et al., 2016); (X. Xu et al., 2015); (Taya et al., 2014); (Martial et al., 2023); (Ryu et al., 2022); (Duan et al., 2014); (Ray et al., 2020); (Suprano et al., 2019); (Alnæs et al., 2015); (Ebrahimi et al., 2019); (Mancini et al., 2017); (F. Fan et al., 2021); (Huckins et al., 2019); (Farah & Horowitz-Kraus, 2019); (Jung et al., 2018); (Kobayashi et al., 2020); (Bailey et al., 2018); (Monge et al., 2017); (Belden et al., 2020); (Geerligs et al., 2014); (Santarnecchi et al., 2014); (X. Li et al., 2020); (Marek et al., 2015); (Shine et al., 2016); (Cao et al., 2019); (File et al., 2016); (Iordan et al., 2018); (Lunsford-Avery et al., 2020); (Smith et al., 2018); (D. Liu et al., 2022); (Anderson et al., 2017); (Tooley et al., 2020); (Le et al., 2020); (C. Wang et al., 2020); (Dan et al., 2023); (L. Song et al., 2020); (Servaas et al., 2017); (Jarrahi & Kollias, 2020); (Q. Li et al., 2019); (Farah & Horowitz-Kraus, 2019); (Neyland et al., 2021); (Geerligs et al., 2015); (C. Wang et al., 2022); (Mancini et al., 2017); (Madden et al., 2020); (Gozdas et al., 2019); (Hearne et al., 2017); (Yue et al., 2017); (Xia et al., 2019); (Markett et al., 2018); (Fukushima et al., 2017); (Gracia-Tabuenca et al., 2021); (Fukushima & Sporns, 2018); (Long et al., 2017); (Manza et al., 2020); (Han et al., 2023); (Fujimoto et al., 2020); (J. Song et al., 2014); (Chong et al., 2019); (Y. Fan et al., 2019); (Orwig et al., 2021); (Najafi et al., 2016); (Tooley et al., 2022); (Kim et al., 2018); (Gratton et al., 2016); (Thompson & Fransson, 2015); (Sato et al., 2015) |
| Interpolate | | (Varangis et al., 2021); (Varangis et al., 2019); (Ryu et al., 2022); (Santarnecchi et al., 2014); (Anderson et al., 2017); (Tooley et al., 2020); (Huckins et al., 2019); (Shine et al., 2016); (Gratton et al., 2016); (Fukushima et al., 2017); (Fukushima & Sporns, 2018); (Y. Fan et al., 2019); (Orwig et al., 2021); (Thompson & Fransson, 2015) |
| SliceTimeCorrect | | (Su et al., 2021); (Zhou et al., 2021); (Zheng et al., 2021); (Varangis et al., 2021); (Hayasaka, 2013); (G. Zhang & Liu, 2021); (Varangis et al., 2019); (Danti et al., 2018); (Le et al., 2020); (X. Liu et al., 2018); (Westphal et al., 2017); (Iordan et al., 2018); (Jarrahi & Kollias, 2020); (Geib et al., 2017); (Taruffi et al., 2017); (Huckins et al., 2019); (Ding et al., 2011); (Finc et al., 2017); (Cole et al., 2015); (Zhong et al., 2014); (Farah & Horowitz-Kraus, 2019); (H. Yan et al., 2022); (Bueichekú et al., 2019); (Gopinath et al., 2015); (Smith et al., 2018); (Rubin et al., 2017); (Vatansever et al., 2015b); (Sreenivasan et al., 2017); (Suprano et al., 2019); (Finc et al., 2020); (Crossley et al., 2013); (Borchardt et al., 2015); (Ekman et al., 2012); (L. Wang et al., 2010); (Cao et al., 2019); (Huang et al., 2018); (Y. Liu et al., 2017); (Berroir et al., 2017); (Prčkovska et al., 2016); (Zhao et al., 2017); (Liang et al., 2016); (Taya et al., 2014); (Koelsch & Skouras, 2014); (Messé et al., 2012); (Martial et al., 2023); |

| | | |
|---|---|---|
| | | (Y. Gao et al., 2023); (Ryu et al., 2022); (Ghiles et al., 2023); (Shen et al., 2013); (Kobayashi et al., 2020); (Y. Wang et al., 2021); (Quante et al., 2018); (C. Yan & He, 2011); (X. Liu et al., 2020); (X. Fan et al., 2021); (Madden et al., 2020); (Parhizi et al., 2018); (Stevens et al., 2012); (Markett et al., 2018); (Tian et al., 2011); (Farahani et al., 2022); (Hilger et al., 2017b); (L. Song et al., 2020); (Bartholomew et al., 2019); (J. Wang et al., 2017); (Pan et al., 2018); (F. Fan et al., 2021); (Chong et al., 2019); (Yue et al., 2017); (Cohen & D'Esposito, 2016); (He et al., 2019); (S. Zhang et al., 2022); (Deng et al., 2016); (Braun et al., 2012); (Vatansever et al., 2015a); (Sheppard et al., 2011); (Liang et al., 2012); (Beaty et al., 2015); (Duan et al., 2014); (Q. Li et al., 2019); (Meunier et al., 2014); (Breckel et al., 2013); (Markett et al., 2013); (Wu et al., 2013); (P. Xu et al., 2014); (J. Liu et al., 2017); (Xiao et al., 2016); (C. Wang et al., 2022); (X. Li et al., 2020); (Jung et al., 2018); (Sun et al., 2017); (Y. Gu et al., 2022); (C. Wang et al., 2020); (D. Liu et al., 2022); (Göttlich et al., 2017); (Lin et al., 2022); (X. Zhang et al., 2015); (J. Zhang et al., 2021); (Jin et al., 2020); (Xi et al., 2019); (Agrawal et al., 2019); (Brandl et al., 2018); (Knyazeva et al., 2018); (Aggarwal et al., 2017); (Shang et al., 2017); (Hilger et al., 2017a); (File et al., 2016); (Du et al., 2015); (Wen et al., 2015); (Cocchi et al., 2015); (DeSalvo et al., 2014); (Liang et al., 2013); (Spreng et al., 2013); (Sheppard et al., 2011); (Franzmeier et al., 2018); (Choi et al., 2023); (X. Chen et al., 2023); (H. Wang et al., 2023); (Gracia-Tabuenca et al., 2021); (J. Song et al., 2014); (Kawagoe et al., 2017); (Monge et al., 2017); (Xia et al., 2019); (Pindus et al., 2020); (Santarnecchi et al., 2014); (Sheppard et al., 2012); (Polanía et al., 2011); (Yi et al., 2023); (Y. Fan et al., 2021); (Ebrahimi et al., 2019); (Zamroziewicz et al., 2017); (Mancini et al., 2017); (Gratton et al., 2016); (Satterthwaite et al., 2012); (Dan et al., 2023); (Han et al., 2023); (Lee et al., 2022); (Belden et al., 2020); (Y. Fan et al., 2019); (Orwig et al., 2021); (Qin et al., 2016); (X. Xu et al., 2015); (Richards et al., 2018); (Malagurski et al., 2020); (Long et al., 2017); (Neyland et al., 2021); (Koba et al., 2021); (Amlien et al., 2019) |
| FieldMapCorrect | | (Ray et al., 2020); (Knyazeva et al., 2018); (Lee et al., 2022); (Tooley et al., 2020); (Santarnecchi et al., 2014); (Spielberg et al., 2015); (F. Fan et al., 2021); (Geerligs et al., 2015); (Franzmeier et al., 2018); (Shine et al., 2016); (Amlien et al., 2019); (S. Gao et al., 2021); (Fujimoto et al., 2020); (Fukushima et al., 2017); (Zuo et al., 2018); (Ogawa, 2021); (Kruschwitz et al., 2018); (Yin et al., 2019); (Pezoulas et al., 2017); (Fukushima & Sporns, 2018); (C. Zhang et al., 2016); (Liao et al., 2017); (T. Chen et al., 2016); (Lloyd, 2020); (Manza et al., 2020); (Jacob et al., 2016); (Y. Zhang et al., 2022); (Betzel et al., 2020); (Tipnis et al., 2020); (Kim et al., 2018); (Ma & Zhang, 2017); (Bolt et al., 2017); (Najafi et al., 2016); (Thompson & Fransson, 2015); (Khodaei et al., 2023) |
| Unwarp | | (Ketchabaw et al., 2022); (Belden et al., 2020); (Dan et al., 2023); (Kobayashi et al., 2020); (Westphal et al., 2017); (Jarrahi & Kollias, 2020); (Smith et al., 2018); (Rubin et al., 2017); (Tooley et al., 2022); (Foo et al., 2021); (S. Gao et al., 2021); (Fujimoto et al., 2020); (Fukushima et al., 2017); (Zuo et al., 2018); (Ogawa, 2021); (Kruschwitz et al., 2018); (Yin et al., 2019); (Pezoulas et al., 2017); (Fukushima & Sporns, 2018); (C. Zhang et al., 2016); (Liao et al., 2017); (T. Chen et al., 2016); (Lloyd, 2020); (Manza et al., 2020); (Jacob et al., 2016); (Y. Zhang et al., 2022); (Tipnis et al., 2020); (Kim et al., 2018); (Ma & Zhang, 2017); (Bolt et al., 2017); (Najafi et al., 2016); (Thompson & Fransson, 2015); (Khodaei et al., 2023) |
| EchoCombine | | (Setton et al., 2022) |
| IntensNormalize | | (Gratton et al., 2016); (Huckins et al., 2019); (de Pasquale et al., 2017); (Stevens et al., 2012); (Gracia-Tabuenca et al., 2021); (Hilger et al., 2017b); (Orwig et al., 2021); (Reddy et al., 2018); (Hilger et al., 2017a); (Foo et al., 2021); (Amlien et al., 2019); (Manza et al., 2020); (S. Gao et al., 2021); (Fujimoto et al., 2020); (Fukushima et al., 2017); (Zuo et al., 2018); (Ogawa, 2021); (Kruschwitz et al., 2018); (Yin et al., 2019); (Pezoulas et al., 2017); |

| | | |
|---|---|---|
| | | (Fukushima & Sporns, 2018); (C. Zhang et al., 2016); (Liao et al., 2017); (T. Chen et al., 2016); (Lloyd, 2020); (Jacob et al., 2016); (Y. Zhang et al., 2022); (Tipnis et al., 2020); (Kim et al., 2018); (Ma & Zhang, 2017); (Bolt et al., 2017); (Najafi et al., 2016); (Thompson & Fransson, 2015); (Khodaei et al., 2023) |
| GrandMeanScale | | (Belden et al., 2020); (Agrawal et al., 2019); (Danti et al., 2018); (Ketchabaw et al., 2022); (Chong et al., 2019); (Schlesinger et al., 2017); (Zhong et al., 2014); (Sato et al., 2015); (H. Zhang et al., 2012); (Satterthwaite et al., 2012); (Gratton et al., 2016); (Westphal et al., 2017); (Bolt et al., 2017); (Kim et al., 2018) |
| Coregister | | (Moussa et al., 2012); (Masuda et al., 2018); (Servaas et al., 2017); (Rzucidlo et al., 2013); (Geerligs et al., 2014); (Ray et al., 2020); (Geerligs et al., 2015); (Alavash, Doebler, et al., 2015); (Reddy et al., 2018); (de Paula et al., 2017); (Quante et al., 2018); (Feng et al., 2015); (Bailey et al., 2018); (Le et al., 2020); (X. Liu et al., 2018); (Iordan et al., 2018); (Finc et al., 2017); (Alluri et al., 2017); (Koba et al., 2021); (Breckel et al., 2013); (Vatansever et al., 2015b); (Hearne et al., 2017); (Sreenivasan et al., 2017); (Suprano et al., 2019); (Finc et al., 2020); (Crossley et al., 2013); (Borchardt et al., 2015); (Shine et al., 2016); (Cao et al., 2019); (Huang et al., 2018); (Y. Liu et al., 2017); (Kobayashi et al., 2020); (Su et al., 2021); (Zheng et al., 2021); (C. Yan & He, 2011); (X. Fan et al., 2021); (J. Wang et al., 2017); (Yue et al., 2017); (Cohen & D'Esposito, 2016); (Westphal et al., 2017); (Pamplona et al., 2015); (S. Zhang et al., 2022); (Jarrahi & Kollias, 2020); (Deng et al., 2016); (Sheppard et al., 2011); (Beaty et al., 2015); (Gopinath et al., 2015); (Lunsford-Avery et al., 2020); (Smith et al., 2018); (Xiao et al., 2016); (C. Wang et al., 2022); (Jung et al., 2018); (Sun et al., 2017); (C. Wang et al., 2020); (Göttlich et al., 2017); (Lin et al., 2022); (Brandl et al., 2018); (Berroir et al., 2017); (Gratton et al., 2016); (Franzmeier et al., 2018); (Martial et al., 2023); (Dan et al., 2023); (Ghiles et al., 2023); (H. Wang et al., 2023); (Invernizzi et al., 2023); (Lee et al., 2022); (Setton et al., 2022); (Farahani et al., 2022); (Pan et al., 2018); (Xia et al., 2019); (J. Liu et al., 2017); (Y. Gu et al., 2022); (Polanía et al., 2011); (J. Zhang et al., 2021); (Jin et al., 2020); (Ebrahimi et al., 2019); (Mancini et al., 2017); (Du et al., 2015); (X. Xu et al., 2015); (DeSalvo et al., 2014); (Arnold et al., 2014); (X. Chen et al., 2023); (J. Song et al., 2014); (Parhizi et al., 2018); (Gozdas et al., 2019); (F. Fan et al., 2021); (Y. Fan et al., 2019); (Tooley et al., 2022); (de Pasquale et al., 2017); (Richards et al., 2018); (Han et al., 2023); (Malagurski et al., 2020); (Evensmoen et al., 2021); (Long et al., 2017); (Monge et al., 2017); (Neyland et al., 2021); (Reineberg & Banich, 2016); (Shang et al., 2017); (Liang et al., 2016); (S. Gu et al., 2015); (Liang et al., 2013); (Choi et al., 2023); (Varangis et al., 2021); (Markett et al., 2018); (Santarnecchi et al., 2014); (Kolskår et al., 2018); (Ketchabaw et al., 2022); (Y. Gao et al., 2023); (S. Gao et al., 2021); (Fujimoto et al., 2020); (Fukushima et al., 2017); (Zuo et al., 2018); (Ogawa, 2021); (Kruschwitz et al., 2018); (Yin et al., 2019); (Pezoulas et al., 2017); (Varangis et al., 2019); (Fukushima & Sporns, 2018); (Chong et al., 2019); (C. Zhang et al., 2016); (Liao et al., 2017); (Pindus et al., 2020); (Schlesinger et al., 2017); (T. Chen et al., 2016); (Lloyd, 2020); (Sato et al., 2015); (Manza et al., 2020); (Jacob et al., 2016); (Y. Zhang et al., 2022); (Tipnis et al., 2020); (Amlien et al., 2019); (Kim et al., 2018); (Zamroziewicz et al., 2017); (Ma & Zhang, 2017); (Bolt et al., 2017); (Hilger et al., 2017a); (Najafi et al., 2016); (Thompson & Fransson, 2015); (Khodaei et al., 2023); (Gracia-Tabuenca et al., 2021); (Vriend et al., 2020); (Orwig et al., 2021); (Satterthwaite et al., 2012); (Betzel et al., 2020); (Tooley et al., 2020) |
| FuncSpaNorm | MNI | (Kobayashi et al., 2020); (Y. Wang et al., 2021); (Zhou et al., 2021); (Zheng et al., 2021); (Masuda et al., 2018); (Hayasaka, 2013); (X. Liu et al., 2020); (X. Fan et al., 2021); (Tian et al., 2011); (Farahani et al., 2022); (Hilger et al., 2017b); (L. Song et al., 2020); (Feng et al., 2015); (Bartholomew et al., 2019); (Bailey et al., 2018); (Xia et al., 2019); (Westphal et al., 2017); (He et al., 2019); (S. Zhang et al., 2022); (Jarrahi & Kollias, 2020); (Rzucidlo |

| | | |
|---|---|---|
| | | et al., 2013); (Taruffi et al., 2017); (Braun et al., 2012); (Ding et al., 2011); (Finc et al., 2017); (Vatansever et al., 2015a); (Liang et al., 2012); (Duan et al., 2014); (Farah & Horowitz-Kraus, 2019); (Neyland et al., 2021); (Alluri et al., 2017); (Ray et al., 2020); (H. Zhang et al., 2012); (Santarnecchi et al., 2014); (Sato et al., 2015); (Gopinath et al., 2015); (Alavash, Doebler, et al., 2015); (Smith et al., 2018); (P. Xu et al., 2014); (J. Liu et al., 2017); (Kolskår et al., 2018); (Rubin et al., 2017); (Vatansever et al., 2015b); (Qin et al., 2016); (Sun et al., 2017); (Y. Gu et al., 2022); (Alnæs et al., 2015); (L. Wang et al., 2010); (Lin et al., 2022); (J. Zhang et al., 2021); (Betzel et al., 2020); (Jin et al., 2020); (Moorthigari et al., 2020); (Cao et al., 2019); (Brandl et al., 2018); (Huang et al., 2018); (Knyazeva et al., 2018); (de Pasquale et al., 2017); (Prčkovska et al., 2016); (de Paula et al., 2017); (Anderson et al., 2017); (Hilger et al., 2017a); (Alavash et al., 2016); (S. Gu et al., 2015); (Du et al., 2015); (Alavash, Hilgetag, et al., 2015); (Wen et al., 2015); (X. Xu et al., 2015); (Cocchi et al., 2015); (Taya et al., 2014); (Arnold et al., 2014); (Koelsch & Skouras, 2014); (Spreng et al., 2013); (Moussa et al., 2012); (Franzmeier et al., 2018); (Martial et al., 2023); (Dan et al., 2023); (Breedt et al., 2022); (Choi et al., 2023); (X. Chen et al., 2023); (Han et al., 2023); (Y. Gao et al., 2023); (Ryu et al., 2022); (Ghiles et al., 2023); (H. Wang et al., 2023); (Invernizzi et al., 2023); (Lee et al., 2022) |
| | ICBM | (Liang et al., 2016); (Liang et al., 2013) |
| | StudySpecificS | (Madden et al., 2020); (Yang et al., 2023) |
| | Talairach | (Danti et al., 2018); (Yue et al., 2017); (Huckins et al., 2019); (Amlien et al., 2019); (Gratton et al., 2016) |
| | Not reported | (Shen et al., 2013); (S. Gao et al., 2021); (Fujimoto et al., 2020); (Fukushima et al., 2017); (Gracia-Tabuenca et al., 2021); (Foo et al., 2021); (Quante et al., 2018); (Su et al., 2021); (Tooley et al., 2020); (C. Yan & He, 2011); (Zuo et al., 2018); (Vriend et al., 2020); (Varangis et al., 2021); (Ogawa, 2021); (Kruschwitz et al., 2018); (Yin et al., 2019); (G. Zhang & Liu, 2021); (Malagurski et al., 2020); (J. Song et al., 2014); (Setton et al., 2022); (Evensmoen et al., 2021); (Pezoulas et al., 2017); (Parhizi et al., 2018); (Gozdas et al., 2019); (Stevens et al., 2012); (Markett et al., 2018); (Varangis et al., 2019); (Le et al., 2020); (Fukushima & Sporns, 2018); (J. Wang et al., 2017); (Pan et al., 2018); (Long et al., 2017); (Kawagoe et al., 2017); (F. Fan et al., 2021); (Servaas et al., 2017); (Chong et al., 2019); (Cohen & D'Esposito, 2016); (Monge et al., 2017); (X. Liu et al., 2018); (C. Zhang et al., 2016); (Liao et al., 2017); (Pamplona et al., 2015); (Pindus et al., 2020); (Iordan et al., 2018); (Schlesinger et al., 2017); (Geib et al., 2017); (T. Chen et al., 2016); (Deng et al., 2016); (Belden et al., 2020); (Cole et al., 2015); (Zhong et al., 2014); (Sheppard et al., 2011); (Lloyd, 2020); (Geerligs et al., 2014); (Beaty et al., 2015); (Bottino et al., 2021); (Q. Li et al., 2019); (Meunier et al., 2014); (H. Yan et al., 2022); (Koba et al., 2021); (Breckel et al., 2013); (Y. Fan et al., 2019); (Sheppard et al., 2012); (Orwig et al., 2021); (Bueichekú et al., 2019); (Lunsford-Avery et al., 2020); (Geerligs et al., 2015); (Reineberg & Banich, 2016); (Markett et al., 2013); (Wu et al., 2013); (Xiao et al., 2016); (C. Wang et al., 2022); (Hearne et al., 2017); (Sreenivasan et al., 2017); (Ginestet & Simmons, 2011); (Suprano et al., 2019); (X. Li et al., 2020); (Jung et al., 2018); (Finc et al., 2020); (Spielberg et al., 2015); (Tooley et al., 2022); (Polanía et al., 2011); (Marek et al., 2015); (Crossley et al., 2013); (Manza et al., 2020); (C. Wang et al., 2020); (D. Liu et al., 2022); (Jacob et al., 2016); (Ketchabaw et al., 2022); (Göttlich et al., 2017); (Borchardt et al., 2015); (Ekman et al., 2012); (Burdette et al., 2010); (Shine et al., 2016); (X. Zhang et al., 2015); (Y. Zhang et al., 2022); (Yi et al., 2023); (Y. Fan et al., 2021); (Tipnis et al., 2020); (Ebrahimi et al., 2019); (Xi et al., 2019); (Agrawal et al., 2019); (Huskey et al., 2018); (Kim et al., 2018); (Reddy et al., 2018); (Aggarwal et al., 2017); (Zamroziewicz et al., 2017); (Ma & Zhang, 2017); (Y. Liu et al., 2017); (Berroir et al., |

| | | |
|---|---|---|
| | | 2017); (Mancini et al., 2017); (Bolt et al., 2017); (Shang et al., 2017); (Zhao et al., 2017); (File et al., 2016); (Najafi et al., 2016); (Thompson & Fransson, 2015); (DeSalvo et al., 2014); (Sami & Miall, 2013); (Satterthwaite et al., 2012); (Messé et al., 2012); (Sheppard et al., 2011); (Richards et al., 2018); (Khodaei et al., 2023) |
| SpaSmooth | 2mm | (Pezoulas et al., 2017); (de Pasquale et al., 2017) |
| | 3mm | (Pindus et al., 2020); (Zamroziewicz et al., 2017) |
| | 4mm | (Y. Wang et al., 2021); (Yin et al., 2019); (J. Song et al., 2014); (Farahani et al., 2022); (Le et al., 2020); (Pan et al., 2018); (F. Fan et al., 2021); (Yue et al., 2017); (He et al., 2019); (J. Liu et al., 2017); (Hearne et al., 2017); (Tooley et al., 2022); (Sun et al., 2017); (Lin et al., 2022); (Bolt et al., 2017); (Shang et al., 2017); (Du et al., 2015); (Franzmeier et al., 2018) |
| | 5mm | (Foo et al., 2021); (Su et al., 2021); (Masuda et al., 2018); (Vriend et al., 2020); (G. Zhang & Liu, 2021); (Bailey et al., 2018); (Monge et al., 2017); (Pamplona et al., 2015); (Schlesinger et al., 2017); (Gopinath et al., 2015); (Ebrahimi et al., 2019); (Agrawal et al., 2019); (Breedt et al., 2022) |
| | 6mm | (Shen et al., 2013); (Fujimoto et al., 2020); (Zhou et al., 2021); (Zheng et al., 2021); (Stevens et al., 2012); (Markett et al., 2018); (Hilger et al., 2017b); (L. Song et al., 2020); (Feng et al., 2015); (Danti et al., 2018); (Chong et al., 2019); (Cohen & D’Esposito, 2016); (Xia et al., 2019); (Westphal et al., 2017); (S. Zhang et al., 2022); (Jarrahi & Kollias, 2020); (Geib et al., 2017); (Taruffi et al., 2017); (Huckins et al., 2019); (Cole et al., 2015); (Koba et al., 2021); (Orwig et al., 2021); (Bueichekú et al., 2019); (Lunsford-Avery et al., 2020); (Markett et al., 2013); (Kolskår et al., 2018); (Qin et al., 2016); (Xiao et al., 2016); (C. Wang et al., 2022); (Suprano et al., 2019); (Marek et al., 2015); (Alnæs et al., 2015); (C. Wang et al., 2020); (Ketchabaw et al., 2022); (Y. Fan et al., 2021); (Jin et al., 2020); (Prčkovska et al., 2016); (Hilger et al., 2017a); (Gratton et al., 2016); (File et al., 2016); (Liang et al., 2016); (Cocchi et al., 2015); (Koelsch & Skouras, 2014); (Liang et al., 2013); (Spreng et al., 2013); (Satterthwaite et al., 2012); (Martial et al., 2023); (Choi et al., 2023); (X. Chen et al., 2023); (Han et al., 2023); (Yang et al., 2023); (H. Wang et al., 2023) |
| | 8mm | (Kobayashi et al., 2020); (Quante et al., 2018); (X. Liu et al., 2020); (Gozdas et al., 2019); (Kawagoe et al., 2017); (Servaas et al., 2017); (X. Liu et al., 2018); (Belden et al., 2020); (Ding et al., 2011); (Vatansever et al., 2015a); (Geerligs et al., 2014); (Beaty et al., 2015); (Q. Li et al., 2019); (Farah & Horowitz-Kraus, 2019); (Meunier et al., 2014); (H. Yan et al., 2022); (Alluri et al., 2017); (H. Zhang et al., 2012); (Sato et al., 2015); (Geerligs et al., 2015); (Smith et al., 2018); (Vatansever et al., 2015b); (Sreenivasan et al., 2017); (X. Li et al., 2020); (Göttlich et al., 2017); (X. Zhang et al., 2015); (J. Zhang et al., 2021); (Cao et al., 2019); (Brandl et al., 2018); (Huang et al., 2018); (Reddy et al., 2018); (Aggarwal et al., 2017); (Y. Liu et al., 2017); (Berroir et al., 2017); (de Paula et al., 2017); (Anderson et al., 2017); (Zhao et al., 2017); (Taya et al., 2014); (Arnold et al., 2014); (Sheppard et al., 2011); (Invernizzi et al., 2023); (Lee et al., 2022) |
| | Not reported | (Parhizi et al., 2018); (T. Chen et al., 2016); (Jung et al., 2018); (Y. Zhang et al., 2022); (Yi et al., 2023); (Moorthigari et al., 2020); (X. Xu et al., 2015); (Richards et al., 2018) |
| RegressorsExtract | | (Shen et al., 2013); (Cohen & D’Esposito, 2016); (Quante et al., 2018); (Tooley et al., 2022) |
| FuncSurface | | (DeSalvo et al., 2014); (Sheppard et al., 2011); (Sheppard et al., 2012); (Zhong et al., 2014); (S. Gao et al., 2021); (Zuo et al., 2018); (Ogawa, 2021); (Kruschwitz et al., 2018); (Pezoulas et al., 2017); (Lloyd, 2020); (Jacob et al., 2016); (Y. Zhang et al., 2022); (Tipnis et al., 2020); (Kim et al., 2018); (Ma & Zhang, 2017); (Najafi et al., 2016); (Khodaei et al., 2023); (Fujimoto et al., 2020); (Tooley et al., 2022) |

| | | |
|---|---|---|
| RETROICOR | | (Anderson et al., 2017); (Y. Fan et al., 2019); (Choi et al., 2023) |
| RVHRCOR | | (Y. Fan et al., 2019) |
| ANATICOR | | (Cole et al., 2015) |
| PESTICA | | (de Pasquale et al., 2017) |
| CompCor | | (Bottino et al., 2021); (Vatansever et al., 2015a); (Beaty et al., 2015); (Xi et al., 2019); (Xiao et al., 2016); (Hearne et al., 2017); (Kobayashi et al., 2020); (Vatansever et al., 2015b) |
| tCompCor | | (Finc et al., 2020) |
| aCompCor | | (Finc et al., 2020); (Arnold et al., 2014); (Yang et al., 2023); (Bailey et al., 2018); (Farah & Horowitz-Kraus, 2019); (J. Wang et al., 2017); (Iordan et al., 2018); (Finc et al., 2017); (Gozdas et al., 2019); (Smith et al., 2018); (Lee et al., 2022); (Le et al., 2020); (Ebrahimi et al., 2019); (Dan et al., 2023); (Jarrahi & Kollias, 2020) |
| ME_ICA | | (Setton et al., 2022) |
| ArtRepair | | (de Paula et al., 2017); (Lee et al., 2022) |
| TempDetrend | Linear | (Tooley et al., 2020); (G. Zhang & Liu, 2021); (X. Liu et al., 2020); (X. Fan et al., 2021); (Parhizi et al., 2018); (Hilger et al., 2017b); (Bartholomew et al., 2019); (Le et al., 2020); (Pan et al., 2018); (Long et al., 2017); (F. Fan et al., 2021); (X. Liu et al., 2018); (Liao et al., 2017); (Jarrahi & Kollias, 2020); (Deng et al., 2016); (Vatansever et al., 2015a); (Bottino et al., 2021); (Duan et al., 2014); (Santarnecchi et al., 2014); (Sheppard et al., 2012); (Bueichekú et al., 2019); (Markett et al., 2013); (J. Liu et al., 2017); (Kolskår et al., 2018); (C. Wang et al., 2022); (Hearne et al., 2017); (Jung et al., 2018); (Finc et al., 2020); (Alnæs et al., 2015); (Lin et al., 2022); (X. Zhang et al., 2015); (J. Zhang et al., 2021); (Ebrahimi et al., 2019); (Xi et al., 2019); (Liang et al., 2016); (Wen et al., 2015); (X. Xu et al., 2015); (Cocchi et al., 2015); (Liang et al., 2013); (Sheppard et al., 2011); (X. Chen et al., 2023); (H. Wang et al., 2023); (Lee et al., 2022) |
| | Quadratic | (Fukushima et al., 2017); (Danti et al., 2018); (Chong et al., 2019); (He et al., 2019); (Geerligs et al., 2014); (Orwig et al., 2021); (Ekman et al., 2012); (Knyazeva et al., 2018); (Berroir et al., 2017); (Hilger et al., 2017a); (Messé et al., 2012); (Yang et al., 2023) |
| | Spline | (Alluri et al., 2017) |
| | Polynomial | (Yue et al., 2017); (Sato et al., 2015); (Spielberg et al., 2015); (Borchardt et al., 2015); (Bolt et al., 2017) |
| | Not reported | (Shen et al., 2013); (Quante et al., 2018); (Zheng et al., 2021); (Yin et al., 2019); (Fukushima & Sporns, 2018); (Huckins et al., 2019); (Y. Fan et al., 2019); (Gopinath et al., 2015); (Y. Gu et al., 2022); (Ketchabaw et al., 2022); (Jin et al., 2020); (Huskey et al., 2018); (Gratton et al., 2016); (Thompson & Fransson, 2015); (Du et al., 2015); (Y. Gao et al., 2023) |
| GlobalSigRegress | | (Koelsch & Skouras, 2014); (Sheppard et al., 2011); (Hayasaka, 2013); (Rzucidlo et al., 2013); (Cole et al., 2015); (Sheppard et al., 2012); (Bueichekú et al., 2019); (Geerligs et al., 2015); (Spielberg et al., 2015); (Burdette et al., 2010); (Sami & Miall, 2013); (Markett et al., 2018); (Servaas et al., 2017); (Taruffi et al., 2017); (Braun et al., 2012); (Liang et al., 2012); (X. Zhang et al., 2015); (Shang et al., 2017); (Zhou et al., 2021); (Masuda et al., 2018); (Ding et al., 2011); (Q. Li et al., 2019); (P. Xu et al., 2014); (Shine et al., 2016); (Moussa et al., 2012); (H. Wang et al., 2023); (Y. Liu et al., 2017); (Cocchi et al., 2015); (Gracia-Tabuenca et al., 2021); (Zheng et al., 2021); (Stevens et al., 2012); (Feng et al., 2015); (Huckins et al., 2019); (Lunsford-Avery et al., 2020); (Markett |

| | | |
|---|---|---|
| | | et al., 2013); (Betzel et al., 2020); (Gratton et al., 2016); (DeSalvo et al., 2014); (Spreng et al., 2013); (Long et al., 2017); (Jarrahi & Kollias, 2020); (Wu et al., 2013); (Cao et al., 2019); (Satterthwaite et al., 2012); (Malagurski et al., 2020); (Hilger et al., 2017b); (He et al., 2019); (Qin et al., 2016); (Y. Gu et al., 2022); (C. Wang et al., 2020); (Liang et al., 2013); (Tooley et al., 2020); (Chong et al., 2019); (J. Liu et al., 2017); (Tooley et al., 2022); (Betzel et al., 2020); (F. Fan et al., 2021); (Jung et al., 2018); (Ketchabaw et al., 2022); (Tipnis et al., 2020); (Berroir et al., 2017); (Hilger et al., 2017a); (Du et al., 2015); (X. Xu et al., 2015); (Sato et al., 2015); (Kim et al., 2018); (Fukushima et al., 2017); (Yin et al., 2019); (Fukushima & Sporns, 2018) |
| ICA_FIX | | (Huskey et al., 2018); (Kolskår et al., 2018); (Alnæs et al., 2015); (Foo et al., 2021); (Betzel et al., 2020); (Fujimoto et al., 2020); (Kruschwitz et al., 2018); (Ma & Zhang, 2017); (Najafi et al., 2016); (Thompson & Fransson, 2015); (Ogawa, 2021); (Pezoulas et al., 2017); (T. Chen et al., 2016) |
| ICA_AROMA | | (Breedt et al., 2022); (Invernizzi et al., 2023); (Vriend et al., 2020); (Kolskår et al., 2018); (Khodaei et al., 2023) |
| TempFilter | High | Foo et al., 2021); (Quante et al., 2018); (Su et al., 2021); (Vriend et al., 2020); (Ogawa, 2021); (G. Zhang & Liu, 2021); (Pezoulas et al., 2017); (Bartholomew et al., 2019); (Bailey et al., 2018); (Monge et al., 2017); (X. Liu et al., 2018); (Westphal et al., 2017); (Schlesinger et al., 2017); (T. Chen et al., 2016); (Taruffi et al., 2017); (Finc et al., 2017); (Farah & Horowitz-Kraus, 2019); (Meunier et al., 2014); (H. Zhang et al., 2012); (Lunsford-Avery et al., 2020); (Geerligs et al., 2015); (Kolskår et al., 2018); (Vatansever et al., 2015b); (C. Wang et al., 2022); (Alnæs et al., 2015); (C. Wang et al., 2020); (Jacob et al., 2016); (L. Wang et al., 2010); (Yi et al., 2023); (Betzel et al., 2020); (Agrawal et al., 2019); (Huskey et al., 2018); (Knyazeva et al., 2018); (de Pasquale et al., 2017); (Y. Liu et al., 2017); (Liang et al., 2016); (Wen et al., 2015); (DeSalvo et al., 2014); (Franzmeier et al., 2018) |
| | Band | (Shen et al., 2013); (Kobayashi et al., 2020); (Fukushima et al., 2017); (Gracia-Tabuenca et al., 2021); (Y. Wang et al., 2021); (Tooley et al., 2020); (Masuda et al., 2018); (Varangis et al., 2021); (Hayasaka, 2013); (Yin et al., 2019); (X. Liu et al., 2020); (X. Fan et al., 2021); (Malagurski et al., 2020); (J. Song et al., 2014); (Madden et al., 2020); (Gozdas et al., 2019); (Stevens et al., 2012); (Markett et al., 2018); (Tian et al., 2011); (Farahani et al., 2022); (Hilger et al., 2017b); (L. Song et al., 2020); (Varangis et al., 2019); (Feng et al., 2015); (Fukushima & Sporns, 2018); (J. Wang et al., 2017); (Pan et al., 2018); (Long et al., 2017); (Kawagoe et al., 2017); (F. Fan et al., 2021); (Servaas et al., 2017); (Chong et al., 2019); (Yue et al., 2017); (Cohen & D'Esposito, 2016); (Xia et al., 2019); (Liao et al., 2017); (He et al., 2019); (Pamplona et al., 2015); (Pindus et al., 2020); (Iordan et al., 2018); (Jarrahi & Kollias, 2020); (Rzucidlo et al., 2013); (Deng et al., 2016); (Belden et al., 2020); (Huckins et al., 2019); (Braun et al., 2012); (Ding et al., 2011); (Cole et al., 2015); (Zhong et al., 2014); (Vatansever et al., 2015a); (Liang et al., 2012); (Beaty et al., 2015); (Bottino et al., 2021); (Duan et al., 2014); (Q. Li et al., 2019); (Neyland et al., 2021); (Koba et al., 2021); (Breckel et al., 2013); (Santarnecchi et al., 2014); (Sato et al., 2015); (Y. Fan et al., 2019); (Sheppard et al., 2012); (Bueichekú et al., 2019); (Reineberg & Banich, 2016); (Alavash, Doebler, et al., 2015); (Markett et al., 2013); (Wu et al., 2013); (Smith et al., 2018); (P. Xu et al., 2014); (J. Liu et al., 2017); (Rubin et al., 2017); (Qin et al., 2016); (Xiao et al., 2016); (Suprano et al., 2019); (X. Li et al., 2020); (Jung et al., 2018); (Finc et al., 2020); (Tooley et al., 2022); (Sun et al., 2017); (Polanía et al., 2011); (Marek et al., 2015); (Manza et al., 2020); (D. Liu et al., 2022); (Borchardt et al., 2015); (Ekman et al., 2012); (Burdette et al., 2010); (Shine et al., 2016); (Lin et al., 2022); (X. Zhang et al., 2015); (J. Zhang et al., 2021); (Y. Fan et al., 2021); (Tipnis et al., 2020); (Ebrahimi et al., 2019); (Cao et al., 2019); (Kim et al., 2018); (Aggarwal et al., 2017); (Zamroziewicz et al., 2017); (Bolt et al., 2017); (Prčkovska et al., 2016); (Shang et al., 2017); (Hilger et al., 2017a); (Gratton et al., 2016); (File et al., 2016); (Alavash et al., 2016); (S. Gu et al., 2015); (Du et al., |

| | | |
|---|---|---|
| | | 2015); (X. Xu et al., 2015); (Cocchi et al., 2015); (Taya et al., 2014); (Arnold et al., 2014); (Sami & Miall, 2013); (Liang et al., 2013); (Moussa et al., 2012); (Satterthwaite et al., 2012); (Martial et al., 2023); (Dan et al., 2023); (X. Chen et al., 2023); (Han et al., 2023); (Y. Gao et al., 2023); (Khodaei et al., 2023); (Ryu et al., 2022); (Ghiles et al., 2023); (Yang et al., 2023); (H. Wang et al., 2023); (Lee et al., 2022) |
| | Low | (Danti et al., 2018); (Gopinath et al., 2015); (Spreng et al., 2013) |
| | Gaussian | (S. Gao et al., 2021); (Alluri et al., 2017) |
| | FilterCombine | (Zheng et al., 2021) |
| | Not reported | (Parhizi et al., 2018); (Orwig et al., 2021); (Hearne et al., 2017); (Y. Gu et al., 2022); (Crossley et al., 2013); (Ketchabaw et al., 2022); (Moorthigari et al., 2020); (Richards et al., 2018) |
| MotionRegress | 6p | (Zhou et al., 2021); (Masuda et al., 2018); (J. Song et al., 2014); (Stevens et al., 2012); (Hilger et al., 2017b); (Feng et al., 2015); (Bartholomew et al., 2019); (Danti et al., 2018); (Chong et al., 2019); (Iordan et al., 2018); (Deng et al., 2016); (Ding et al., 2011); (Zhong et al., 2014); (Geerligs et al., 2014); (Duan et al., 2014); (Farah & Horowitz-Kraus, 2019); (Neyland et al., 2021); (Meunier et al., 2014); (Alluri et al., 2017); (Koba et al., 2021); (Santarnecchi et al., 2014); (Sato et al., 2015); (Y. Fan et al., 2019); (Bueichekú et al., 2019); (Alavash, Doebler, et al., 2015); (Markett et al., 2013); (Smith et al., 2018); (P. Xu et al., 2014); (J. Liu et al., 2017); (Qin et al., 2016); (Hearne et al., 2017); (Borchardt et al., 2015); (Brandl et al., 2018); (Huang et al., 2018); (Aggarwal et al., 2017); (Prčkovska et al., 2016); (Shang et al., 2017); (Hilger et al., 2017a); (Alavash et al., 2016); (Liang et al., 2016); (Alavash, Hilgetag, et al., 2015); (Wen et al., 2015); (X. Xu et al., 2015); (Cocchi et al., 2015); (DeSalvo et al., 2014); (Liang et al., 2013); (Moussa et al., 2012); (Satterthwaite et al., 2012); (Breedt et al., 2022); (X. Chen et al., 2023); (Khodaei et al., 2023); (Ryu et al., 2022); (Invernizzi et al., 2023) |
| | 12p | (Kobayashi et al., 2020); (Quante et al., 2018); (Su et al., 2021); (Gozdas et al., 2019); (Markett et al., 2018); (Bailey et al., 2018); (Le et al., 2020); (Kawagoe et al., 2017); (Servaas et al., 2017); (Yue et al., 2017); (Pamplona et al., 2015); (Schlesinger et al., 2017); (T. Chen et al., 2016); (Cole et al., 2015); (Vatansever et al., 2015a); (Orwig et al., 2021); (Geerligs et al., 2015); (Vatansever et al., 2015b); (C. Wang et al., 2022); (Marek et al., 2015); (C. Wang et al., 2020); (Ketchabaw et al., 2022); (Ebrahimi et al., 2019); (Zamroziewicz et al., 2017); (Spreng et al., 2013); (Martial et al., 2023); (Dan et al., 2023); (Han et al., 2023); (Lee et al., 2022) |
| | 24p | (S. Gao et al., 2021); (Fukushima et al., 2017); (Y. Wang et al., 2021); (Tooley et al., 2020); (G. Zhang & Liu, 2021); (X. Liu et al., 2020); (X. Fan et al., 2021); (Malagurski et al., 2020); (Madden et al., 2020); (Farahani et al., 2022); (L. Song et al., 2020); (Fukushima & Sporns, 2018); (J. Wang et al., 2017); (Long et al., 2017); (F. Fan et al., 2021); (Xia et al., 2019); (C. Zhang et al., 2016); (Liao et al., 2017); (He et al., 2019); (Q. Li et al., 2019); (Rubin et al., 2017); (Xiao et al., 2016); (Ginestet & Simmons, 2011); (Jung et al., 2018); (Finc et al., 2020); (Tooley et al., 2022); (Sun et al., 2017); (Y. Gu et al., 2022); (Shine et al., 2016); (Lin et al., 2022); (J. Zhang et al., 2021); (Jin et al., 2020); (Cao et al., 2019); (Bolt et al., 2017); (File et al., 2016); (Thompson & Fransson, 2015); (Choi et al., 2023); (Y. Gao et al., 2023); (Yang et al., 2023); (H. Wang et al., 2023) |
| | 36p | (Huckins et al., 2019); (S. Gu et al., 2015) |
| | Not reported | (Gracia-Tabuenca et al., 2021); (Hayasaka, 2013); (Yin et al., 2019); (Pan et al., 2018); (Monge et al., 2017); (X. Liu et al., 2018); (Westphal et al., 2017); (Jarrahi & Kollias, 2020); (Rzucidlo et al., 2013); (Geib et al., 2017); (Taruffi et al., 2017); (Bottino et al., 2021); (Breckel et al., 2013); (Lunsford-Avery et al., 2020); (Wu et al., 2013); (Suprano et al., 2019); (X. Li et al., 2020); (Spielberg et al., 2015); (Burdette et al., 2010); (X. Zhang et |

| | | |
|---|---|---|
| | | al., 2015); (Moorthigari et al., 2020); (Berroir et al., 2017); (Anderson et al., 2017); (Gratton et al., 2016); (Arnold et al., 2014); (Koelsch & Skouras, 2014); (Sami & Miall, 2013); (Richards et al., 2018) |
| ConfoundRegress | | (Braun et al., 2012); (Cole et al., 2015); (Marek et al., 2015); (Xi et al., 2019); (Sami & Miall, 2013); (Y. Gao et al., 2023); (Shen et al., 2013); (Markett et al., 2018); (Alavash, Doebler, et al., 2015); (X. Li et al., 2020); (Anderson et al., 2017); (Shang et al., 2017); (Sheppard et al., 2011); (Choi et al., 2023); (Zhou et al., 2021); (He et al., 2019); (Rzucidlo et al., 2013); (Ding et al., 2011); (Sheppard et al., 2012); (Bueichekú et al., 2019); (Reineberg & Banich, 2016); (Spielberg et al., 2015); (Crossley et al., 2013); (D. Liu et al., 2022); (Burdette et al., 2010); (Koelsch & Skouras, 2014); (Breedt et al., 2022); (Ryu et al., 2022); (Ghiles et al., 2023); (Gracia-Tabuenca et al., 2021); (Varangis et al., 2021); (Hayasaka, 2013); (G. Zhang & Liu, 2021); (L. Song et al., 2020); (Varangis et al., 2019); (Servaas et al., 2017); (Pindus et al., 2020); (Schlesinger et al., 2017); (Zhong et al., 2014); (Duan et al., 2014); (Geerligs et al., 2015); (Rubin et al., 2017); (Finc et al., 2020); (Aggarwal et al., 2017); (Zamroziewicz et al., 2017); (Prčkovska et al., 2016); (Y. Wang et al., 2021); (Su et al., 2021); (Zheng et al., 2021); (Vriend et al., 2020); (Madden et al., 2020); (Bartholomew et al., 2019); (Danti et al., 2018); (Kawagoe et al., 2017); (Pamplona et al., 2015); (Huckins et al., 2019); (Q. Li et al., 2019); (Lunsford-Avery et al., 2020); (Hearne et al., 2017); (Shine et al., 2016); (Alavash et al., 2016); (Alavash, Hilgetag, et al., 2015); (Wen et al., 2015); (Arnold et al., 2014); (Spreng et al., 2013); (Moussa et al., 2012); (H. Wang et al., 2023); (Kobayashi et al., 2020); (Masuda et al., 2018); (X. Liu et al., 2020); (X. Fan et al., 2021); (Cohen & D'Esposito, 2016); (Xia et al., 2019); (Orwig et al., 2021); (C. Wang et al., 2020); (Lin et al., 2022); (X. Zhang et al., 2015); (Y. Liu et al., 2017); (File et al., 2016); (Cocchi et al., 2015); (Satterthwaite et al., 2012); (Martial et al., 2023); (Dan et al., 2023); (X. Chen et al., 2023); (Malagurski et al., 2020); (J. Song et al., 2014); (Stevens et al., 2012); (Farahani et al., 2022); (Feng et al., 2015); (Koba et al., 2021); (Y. Fan et al., 2019); (Markett et al., 2013); (Mancini et al., 2017); (Gratton et al., 2016); (DeSalvo et al., 2014); (Stevens et al., 2012); (Pan et al., 2018); (Long et al., 2017); (Chong et al., 2019); (X. Liu et al., 2018); (Deng et al., 2016); (Belden et al., 2020); (Neyland et al., 2021); (Wu et al., 2013); (J. Liu et al., 2017); (Borchardt et al., 2015); (J. Zhang et al., 2021); (Liang et al., 2016); (Richards et al., 2018); (Han et al., 2023); (Invernizzi et al., 2023); (Lee et al., 2022); (Stevens et al., 2012); (Hilger et al., 2017b); (Smith et al., 2018); (C. Wang et al., 2022); (Suprano et al., 2019); (Jung et al., 2018); (Y. Gu et al., 2022); (Manza et al., 2020); (Ketchabaw et al., 2022); (Jin et al., 2020); (Berroir et al., 2017); (Hilger et al., 2017a); (Du et al., 2015); (Liang et al., 2013); (S. Gao et al., 2021); (Quante et al., 2018); (Sato et al., 2015); (Tooley et al., 2022); (Yin et al., 2019); (F. Fan et al., 2021); (X. Xu et al., 2015); (Fujimoto et al., 2020); (Liao et al., 2017); (Santarnecchi et al., 2014); (Fukushima et al., 2017); (Khodaei et al., 2023); (Fukushima & Sporns, 2018); (Bolt et al., 2017) |
| PhysIO | | |
| RegressorsFilter | | (S. Gu et al., 2015); (Tooley et al., 2020); (Tooley et al., 2022); (Le et al., 2020); (C. Zhang et al., 2016) |
| ModularFilterMotion | | |
| WaveletDecompose | | (Reddy et al., 2018); (Sami & Miall, 2013); (Alavash, Hilgetag, et al., 2015); (S. Gu et al., 2015); (Mancini et al., 2017); (Thompson & Fransson, 2015) |
| AtlasDefine | Power | (Gracia-Tabuenca et al., 2021); (Su et al., 2021); (Zuo et al., 2018); (Varangis et al., 2021); (G. Zhang & Liu, 2021); (J. Song et al., 2014); (Varangis et al., 2019); (Bailey et al., 2018); (Le et al., 2020); (Pan et al., 2018); (Liao et al., 2017); (Westphal et al., 2017); (He et al., 2019); (Iordan et al., 2018); (T. Chen et al., 2016); (Huckins |

| | | |
|---|---|---|
| | | et al., 2019); (Finc et al., 2017); (Cole et al., 2015); (Vatansever et al., 2015a); (Q. Li et al., 2019); (Ray et al., 2020); (Geerligs et al., 2015); (Reineberg & Banich, 2016); (Hearne et al., 2017); (Y. Gu et al., 2022); (Marek et al., 2015); (Manza et al., 2020); (D. Liu et al., 2022); (Huskey et al., 2018); (Brandl et al., 2018); (Bolt et al., 2017); (Gratton et al., 2016); (S. Gu et al., 2015); (Thompson & Fransson, 2015); (Franzmeier et al., 2018); (Ryu et al., 2022) |
| | AAL1 | (Zhou et al., 2021); (Zheng et al., 2021); (C. Yan & He, 2011); (Hayasaka, 2013); (X. Liu et al., 2020); (X. Fan et al., 2021); (Tian et al., 2011); (Feng et al., 2015); (Kawagoe et al., 2017); (X. Liu et al., 2018); (Pamplona et al., 2015); (S. Zhang et al., 2022); (Rzucidlo et al., 2013); (Geib et al., 2017); (Deng et al., 2016); (Belden et al., 2020); (Braun et al., 2012); (Liang et al., 2012); (Duan et al., 2014); (Santarnecchi et al., 2014); (Wu et al., 2013); (P. Xu et al., 2014); (Rubin et al., 2017); (Sreenivasan et al., 2017); (Ginestet & Simmons, 2011); (Sun et al., 2017); (L. Wang et al., 2010); (Yi et al., 2023); (J. Zhang et al., 2021); (Y. Fan et al., 2021); (Xi et al., 2019); (Aggarwal et al., 2017); (Anderson et al., 2017); (Shang et al., 2017); (X. Xu et al., 2015); (Taya et al., 2014); (Martial et al., 2023); (Ghiles et al., 2023) |
| | AAL2 | (C. Zhang et al., 2016); (H. Yan et al., 2022); (Borchardt et al., 2015); (Jin et al., 2020); (Sami & Miall, 2013) |
| | AAL3 | (Choi et al., 2023) |
| | MMP | (Tooley et al., 2020); (Ogawa, 2021) |
| | Brainnetome | (Vriend et al., 2020); (X. Li et al., 2020); (Breedt et al., 2022) |
| | Craddock | (Gozdas et al., 2019); (J. Wang et al., 2017); (Pindus et al., 2020); (Sato et al., 2015); (Spielberg et al., 2015); (Göttlich et al., 2017); (Zamroziewicz et al., 2017); (Cocchi et al., 2015) |
| | ICA | (Kruschwitz et al., 2018); (Stevens et al., 2012); (Xia et al., 2019); (Ding et al., 2011); (Zhong et al., 2014); (Geerligs et al., 2014); (Bottino et al., 2021); (Knyazeva et al., 2018); (de Paula et al., 2017); (Zhao et al., 2017); (Najafi et al., 2016); (Messé et al., 2012); (Yang et al., 2023) |
| | Voxel_wise | (Shen et al., 2013); (Fujimoto et al., 2020); (Kobayashi et al., 2020); (Pezoulas et al., 2017); (Markett et al., 2018); (Hilger et al., 2017b); (L. Song et al., 2020); (Taruffi et al., 2017); (Lloyd, 2020); (Beaty et al., 2015); (Meunier et al., 2014); (Alluri et al., 2017); (Breckel et al., 2013); (H. Zhang et al., 2012); (Y. Fan et al., 2019); (Orwig et al., 2021); (Bueichekú et al., 2019); (Gopinath et al., 2015); (Lunsford-Avery et al., 2020); (Alavash, Doebler, et al., 2015); (Markett et al., 2013); (Smith et al., 2018); (J. Liu et al., 2017); (Kolskår et al., 2018); (Vatansever et al., 2015b); (Qin et al., 2016); (Xiao et al., 2016); (Jung et al., 2018); (Tooley et al., 2022); (Polanía et al., 2011); (Ekman et al., 2012); (Burdette et al., 2010); (Lin et al., 2022); (X. Zhang et al., 2015); (Moorthigari et al., 2020); (Amlien et al., 2019); (Agrawal et al., 2019); (Huang et al., 2018); (Ma & Zhang, 2017); (Y. Liu et al., 2017); (Prčkovska et al., 2016); (Hilger et al., 2017a); (Liang et al., 2016); (Du et al., 2015); (Arnold et al., 2014); (Koelsch & Skouras, 2014); (Liang et al., 2013); (Spreng et al., 2013); (Moussa et al., 2012); (Richards et al., 2018); (X. Chen et al., 2023); (H. Wang et al., 2023) |
| | Schaefer100 | (Foo et al., 2021) |
| | Schaefer200 | (Malagurski et al., 2020); (Farahani et al., 2022) |
| | Schaefer400 | (Setton et al., 2022) |
| | Schaefer500 | (Koba et al., 2021) |

| | | |
|---|---|---|
| | Brodmann | (Parhizi et al., 2018); (Yue et al., 2017) |
| | Desikan_Killiany | (Fukushima et al., 2017); (Fukushima & Sporns, 2018); (Sheppard et al., 2011); (Sheppard et al., 2012); (Suprano et al., 2019); (Berroir et al., 2017); (Sheppard et al., 2011) |
| | Fun160_Dosenbach | (Bartholomew et al., 2019); (C. Wang et al., 2022); (C. Wang et al., 2020); (Wen et al., 2015); (Satterthwaite et al., 2012); (Lee et al., 2022) |
| | Harvard_Oxford | (Madden et al., 2020); (Monge et al., 2017); (Ebrahimi et al., 2019); (Reddy et al., 2018); (File et al., 2016); (Invernizzi et al., 2023) |
| | Destrieux | (Tipnis et al., 2020) |
| | Lausanne2008 | (Schlesinger et al., 2017); (Ketchabaw et al., 2022); (DeSalvo et al., 2014) |
| | Shen | (S. Gao et al., 2021); (Khodaei et al., 2023) |
| | BrainNetwork | (F. Fan et al., 2021); (Chong et al., 2019); (Neyland et al., 2021); (Betzel et al., 2020) |
| | MultipleA | (Y. Wang et al., 2021); (Yin et al., 2019); (Servaas et al., 2017); (Cohen & D'Esposito, 2016); (Finc et al., 2020); (Jacob et al., 2016); (Shine et al., 2016); (Cao et al., 2019); (Kim et al., 2018); (de Pasquale et al., 2017); (Mancini et al., 2017); (Alavash et al., 2016); (Alavash, Hilgetag, et al., 2015); (Dan et al., 2023); (Han et al., 2023); (Y. Gao et al., 2023) |
| | Not reported | (Quante et al., 2018); (Masuda et al., 2018); (Evensmoen et al., 2021); (Danti et al., 2018); (Long et al., 2017); (Jarrahi & Kollias, 2020); (Farah & Horowitz-Kraus, 2019); (Crossley et al., 2013); (Alnæs et al., 2015); (Y. Zhang et al., 2022) |
| ROITSDefine | MeanTS | (S. Gao et al., 2021); (Kobayashi et al., 2020); (Fukushima et al., 2017); (Gracia-Tabuenca et al., 2021); (Su et al., 2021); (Tooley et al., 2020); (Zhou et al., 2021); (C. Yan & He, 2011); (Zuo et al., 2018); (Ogawa, 2021); (Hayasaka, 2013); (Yin et al., 2019); (G. Zhang & Liu, 2021); (X. Liu et al., 2020); (X. Fan et al., 2021); (Malagurski et al., 2020); (J. Song et al., 2014); (Madden et al., 2020); (Pezoulas et al., 2017); (Stevens et al., 2012); (Tian et al., 2011); (Farahani et al., 2022); (Varangis et al., 2019); (Feng et al., 2015); (Bartholomew et al., 2019); (J. Wang et al., 2017); (Pan et al., 2018); (Kawagoe et al., 2017); (F. Fan et al., 2021); (Servaas et al., 2017); (X. Liu et al., 2018); (C. Zhang et al., 2016); (Liao et al., 2017); (Westphal et al., 2017); (He et al., 2019); (Pamplona et al., 2015); (Pindus et al., 2020); (Iordan et al., 2018); (S. Zhang et al., 2022); (Jarrahi & Kollias, 2020); (Schlesinger et al., 2017); (Geib et al., 2017); (T. Chen et al., 2016); (Deng et al., 2016); (Huckins et al., 2019); (Finc et al., 2017); (Cole et al., 2015); (Vatansever et al., 2015a); (Sheppard et al., 2011); (Liang et al., 2012); (Duan et al., 2014); (Q. Li et al., 2019); (Farah & Horowitz-Kraus, 2019); (Neyland et al., 2021); (Koba et al., 2021); (Breckel et al., 2013); (Santarnecchi et al., 2014); (Sato et al., 2015); (Sheppard et al., 2012); (Lunsford-Avery et al., 2020); (Geerligs et al., 2015); (Reineberg & Banich, 2016); (Alavash, Doebler, et al., 2015); (Markett et al., 2013); (Wu et al., 2013); (Smith et al., 2018); (P. Xu et al., 2014); (Rubin et al., 2017); (Vatansever et al., 2015b); (Qin et al., 2016); (C. Wang et al., 2022); (Hearne et al., 2017); (Sreenivasan et al., 2017); (Suprano et al., 2019); (X. Li et al., 2020); (Finc et al., 2020); (Spielberg et al., 2015); (Sun et al., 2017); (Y. Gu et al., 2022); (Marek et al., 2015); (Manza et al., 2020); (C. Wang et al., 2020); (D. Liu et al., 2022); (Ketchabaw et al., 2022); (Borchardt et al., 2015); (Ekman et al., 2012); (L. Wang et al., 2010); (Shine et al., 2016); (Yi et al., 2023); (J. Zhang et al., 2021); (Y. Fan et al., 2021); (Betzel et al., 2020); (Jin et al., 2020); (Tipnis et al., 2020); (Ebrahimi et al., 2019); (Xi et al., 2019); (Huskey et al., 2018); (Kim et al., 2018); (Reddy |

| | | |
|---|---|---|
| | | et al., 2018); (Aggarwal et al., 2017); (Zamroziewicz et al., 2017); (de Pasquale et al., 2017); (Berroir et al., 2017); (Mancini et al., 2017); (Bolt et al., 2017); (Anderson et al., 2017); (Shang et al., 2017); (Gratton et al., 2016); (File et al., 2016); (Alavash et al., 2016); (Thompson & Fransson, 2015); (Alavash, Hilgetag, et al., 2015); (Wen et al., 2015); (X. Xu et al., 2015); (Cocchi et al., 2015); (Taya et al., 2014); (DeSalvo et al., 2014); (Arnold et al., 2014); (Koelsch & Skouras, 2014); (Sami & Miall, 2013); (Spreng et al., 2013); (Messé et al., 2012); (Sheppard et al., 2011); (Richards et al., 2018); (Martial et al., 2023); (Dan et al., 2023); (Breedt et al., 2022); (Choi et al., 2023); (Han et al., 2023); (Y. Gao et al., 2023); (Khodaei et al., 2023); (Ryu et al., 2022); (Ghiles et al., 2023); (Invernizzi et al., 2023); (Lee et al., 2022) |
| | DimensionReduce | (Braun et al., 2012); (Jacob et al., 2016) |
| | Not reported | (Shen et al., 2013); (Fujimoto et al., 2020); (Foo et al., 2021); (Y. Wang et al., 2021); (Quante et al., 2018); (Zheng et al., 2021); (Masuda et al., 2018); (Vriend et al., 2020); (Varangis et al., 2021); (Kruschwitz et al., 2018); (Setton et al., 2022); (Evensmoen et al., 2021); (Parhizi et al., 2018); (Gozdas et al., 2019); (Markett et al., 2018); (Hilger et al., 2017b); (L. Song et al., 2020); (Danti et al., 2018); (Bailey et al., 2018); (Le et al., 2020); (Fukushima & Sporns, 2018); (Long et al., 2017); (Chong et al., 2019); (Yue et al., 2017); (Cohen & D'Esposito, 2016); (Monge et al., 2017); (Xia et al., 2019); (Rzucidlo et al., 2013); (Taruffi et al., 2017); (Belden et al., 2020); (Ding et al., 2011); (Zhong et al., 2014); (Lloyd, 2020); (Geerligs et al., 2014); (Beaty et al., 2015); (Bottino et al., 2021); (Meunier et al., 2014); (H. Yan et al., 2022); (Alluri et al., 2017); (Ray et al., 2020); (H. Zhang et al., 2012); (Y. Fan et al., 2019); (Orwig et al., 2021); (Bueichekú et al., 2019); (Gopinath et al., 2015); (J. Liu et al., 2017); (Kolskår et al., 2018); (Xiao et al., 2016); (Ginestet & Simmons, 2011); (Jung et al., 2018); (Tooley et al., 2022); (Polanía et al., 2011); (Crossley et al., 2013); (Alnæs et al., 2015); (Göttlich et al., 2017); (Burdette et al., 2010); (Lin et al., 2022); (X. Zhang et al., 2015); (Y. Zhang et al., 2022); (Moorthigari et al., 2020); (Amlien et al., 2019); (Agrawal et al., 2019); (Cao et al., 2019); (Brandl et al., 2018); (Huang et al., 2018); (Knyazeva et al., 2018); (Ma & Zhang, 2017); (Y. Liu et al., 2017); (Prčkovska et al., 2016); (de Paula et al., 2017); (Zhao et al., 2017); (Hilger et al., 2017a); (Najafi et al., 2016); (Liang et al., 2016); (S. Gu et al., 2015); (Du et al., 2015); (Liang et al., 2013); (Moussa et al., 2012); (Satterthwaite et al., 2012); (Franzmeier et al., 2018); (X. Chen et al., 2023); (Yang et al., 2023); (H. Wang et al., 2023) |
| ROIDefine | | (Ginestet & Simmons, 2011); (H. Yan et al., 2022); (Ray et al., 2020); (Alavash et al., 2016); (Alavash, Hilgetag, et al., 2015); (Messé et al., 2012); (Geib et al., 2017); (Sheppard et al., 2011); (Ekman et al., 2012); (L. Wang et al., 2010); (Agrawal et al., 2019); (Knyazeva et al., 2018); (Parhizi et al., 2018); (Tian et al., 2011); (Geerligs et al., 2014); (Bottino et al., 2021); (Meunier et al., 2014); (H. Zhang et al., 2012); (Wu et al., 2013); (Sreenivasan et al., 2017); (Polanía et al., 2011); (Crossley et al., 2013); (Burdette et al., 2010); (Yi et al., 2023); (Cao et al., 2019); (Huang et al., 2018); (C. Yan & He, 2011); (Evensmoen et al., 2021); (Braun et al., 2012); (Liang et al., 2012); (Alavash, Doebler, et al., 2015); (Göttlich et al., 2017); (Y. Fan et al., 2021); (Huskey et al., 2018); (Berroir et al., 2017); (Prčkovska et al., 2016); (Taya et al., 2014); (Sheppard et al., 2011); (Y. Wang et al., 2021); (Hayasaka, 2013); (Setton et al., 2022); (Bailey et al., 2018); (Ding et al., 2011); (Zhong et al., 2014); (Breckel et al., 2013); (P. Xu et al., 2014); (Rubin et al., 2017); (Qin et al., 2016); (Spielberg et al., 2015); (Xi et al., 2019); (Reddy et al., 2018); (Sami & Miall, 2013); (Franzmeier et al., 2018); (Breedt et al., 2022); (Ghiles et al., 2023); (Yang et al., 2023); (Varangis et al., 2021); (Bartholomew et al., 2019); (Danti et al., 2018); (Kawagoe et al., 2017); (Cohen & D'Esposito, 2016); (S. Zhang et al., 2022); (Duan et al., 2014); (Farah & Horowitz-Kraus, 2019); (Sheppard et al., 2012); (Bueichekú et al., 2019); (Vatansever et al., 2015b); (Suprano et al., 2019); (D. |

| | | |
|---|---|---|
| | | Liu et al., 2022); (Brandl et al., 2018); (Aggarwal et al., 2017); (de Pasquale et al., 2017); (Y. Liu et al., 2017); (Anderson et al., 2017); (S. Gu et al., 2015); (Wen et al., 2015); (Choi et al., 2023); (Ryu et al., 2022); (Zheng et al., 2021); (G. Zhang & Liu, 2021); (X. Liu et al., 2020); (X. Fan et al., 2021); (Finc et al., 2017); (Cole et al., 2015); (Vatansever et al., 2015a); (Lunsford-Avery et al., 2020); (X. Li et al., 2020); (Marek et al., 2015); (File et al., 2016); (Cocchi et al., 2015); (Arnold et al., 2014); (Spreng et al., 2013); (Invernizzi et al., 2023); (Su et al., 2021); (Vriend et al., 2020); (Madden et al., 2020); (Farahani et al., 2022); (Varangis et al., 2019); (J. Wang et al., 2017); (Servaas et al., 2017); (Yue et al., 2017); (Monge et al., 2017); (Pamplona et al., 2015); (Iordan et al., 2018); (Schlesinger et al., 2017); (Q. Li et al., 2019); (Geerligs et al., 2015); (Reineberg & Banich, 2016); (Markett et al., 2013); (Finc et al., 2020); (Sun et al., 2017); (Mancini et al., 2017); (DeSalvo et al., 2014); (Y. Gao et al., 2023); (Foo et al., 2021); (Quante et al., 2018); (Gozdas et al., 2019); (Feng et al., 2015); (Xia et al., 2019); (X. Liu et al., 2018); (Westphal et al., 2017); (He et al., 2019); (Pindus et al., 2020); (Deng et al., 2016); (Belden et al., 2020); (Koba et al., 2021); (J. Liu et al., 2017); (Shine et al., 2016); (Zamroziewicz et al., 2017); (Shang et al., 2017); (Richards et al., 2018); (Martial et al., 2023); (X. Chen et al., 2023); (Kobayashi et al., 2020); (Zuo et al., 2018); (Malagurski et al., 2020); (Stevens et al., 2012); (Le et al., 2020); (Huckins et al., 2019); (Neyland et al., 2021); (Smith et al., 2018); (C. Wang et al., 2020); (Borchardt et al., 2015); (J. Zhang et al., 2021); (Jin et al., 2020); (Ebrahimi et al., 2019); (Satterthwaite et al., 2012); (H. Wang et al., 2023); (Lee et al., 2022); (Gracia-Tabuenca et al., 2021); (Pan et al., 2018); (Chong et al., 2019); (C. Zhang et al., 2016); (Jarrahi & Kollias, 2020); (C. Wang et al., 2022); (Jung et al., 2018); (Y. Gu et al., 2022); (Manza et al., 2020); (Hilger et al., 2017a); (Dan et al., 2023); (Han et al., 2023); (Ogawa, 2021); (J. Song et al., 2014); (Pezoulas et al., 2017); (Hearne et al., 2017); (Ketchabaw et al., 2022); (Betzel et al., 2020); (Tipnis et al., 2020); (X. Xu et al., 2015); (Fujimoto et al., 2020); (Tooley et al., 2020); (T. Chen et al., 2016); (Santarnecchi et al., 2014); (Kim et al., 2018); (Gratton et al., 2016); (S. Gao et al., 2021); (F. Fan et al., 2021); (Liao et al., 2017); (Sato et al., 2015); (Tooley et al., 2022); (Thompson & Fransson, 2015); (Khodaei et al., 2023); (Yin et al., 2019); (Fukushima & Sporns, 2018); (Bolt et al., 2017); (Fukushima et al., 2017) |
| BlockCombine | Concatenated | (Zhou et al., 2021); (Varangis et al., 2021); (Ogawa, 2021); (X. Liu et al., 2018); (C. Wang et al., 2022); (J. Zhang et al., 2021); (Bolt et al., 2017); (Prčkovska et al., 2016); (Messé et al., 2012); (Sheppard et al., 2011) |
| | BlockWise | (Zuo et al., 2018); (Yue et al., 2017); (Cohen & D'Esposito, 2016); (Tipnis et al., 2020); (Liang et al., 2016); (Du et al., 2015) |
| | Not reported | |
| ConnectCompute | Pearson | (S. Gao et al., 2021); (Fujimoto et al., 2020); (Kobayashi et al., 2020); (Fukushima et al., 2017); (Gracia-Tabuenca et al., 2021); (Quante et al., 2018); (Su et al., 2021); (Tooley et al., 2020); (Zhou et al., 2021); (Zheng et al., 2021); (Zuo et al., 2018); (Varangis et al., 2021); (Ogawa, 2021); (Hayasaka, 2013); (Yin et al., 2019); (X. Liu et al., 2020); (X. Fan et al., 2021); (Malagurski et al., 2020); (J. Song et al., 2014); (Setton et al., 2022); (Madden et al., 2020); (Pezoulas et al., 2017); (Gozdas et al., 2019); (Stevens et al., 2012); (Markett et al., 2018); (Tian et al., 2011); (Farahani et al., 2022); (Hilger et al., 2017b); (L. Song et al., 2020); (Varangis et al., 2019); (Feng et al., 2015); (Bartholomew et al., 2019); (Danti et al., 2018); (Bailey et al., 2018); (Le et al., 2020); (Fukushima & Sporns, 2018); (J. Wang et al., 2017); (Pan et al., 2018); (Kawagoe et al., 2017); (F. Fan et al., 2021); (Servaas et al., 2017); (Chong et al., 2019); (Yue et al., 2017); (Cohen & D'Esposito, 2016); (Monge et al., 2017); (Xia et al., 2019); (X. Liu et al., 2018); (C. Zhang et al., 2016); (Liao et al., 2017); (Westphal et al., 2017); (He et al., 2019); (Pamplona et al., 2015); (Pindus et al., 2020); (Iordan et al., 2018); (S. Zhang et al., |

| | | |
|---|---|---|
| | | 2022); (Jarrahi & Kollias, 2020); (Rzucidlo et al., 2013); (Geib et al., 2017); (T. Chen et al., 2016); (Deng et al., 2016); (Huckins et al., 2019); (Braun et al., 2012); (Ding et al., 2011); (Finc et al., 2017); (Cole et al., 2015); (Zhong et al., 2014); (Sheppard et al., 2011); (Geerligs et al., 2014); (Bottino et al., 2021); (Duan et al., 2014); (Farah & Horowitz-Kraus, 2019); (Neyland et al., 2021); (Meunier et al., 2014); (H. Yan et al., 2022); (Alluri et al., 2017); (Koba et al., 2021); (Ray et al., 2020); (Santarnecchi et al., 2014); (Sato et al., 2015); (Y. Fan et al., 2019); (Sheppard et al., 2012); (Orwig et al., 2021); (Bueichekú et al., 2019); (Gopinath et al., 2015); (Lunsford-Avery et al., 2020); (Geerligs et al., 2015); (Reineberg & Banich, 2016); (Alavash, Doebler, et al., 2015); (Markett et al., 2013); (Wu et al., 2013); (Smith et al., 2018); (P. Xu et al., 2014); (J. Liu et al., 2017); (Rubin et al., 2017); (Vatansever et al., 2015b); (Qin et al., 2016); (Xiao et al., 2016); (C. Wang et al., 2022); (Hearne et al., 2017); (X. Li et al., 2020); (Jung et al., 2018); (Finc et al., 2020); (Spielberg et al., 2015); (Tooley et al., 2022); (Sun et al., 2017); (Y. Gu et al., 2022); (Polanía et al., 2011); (Marek et al., 2015); (Crossley et al., 2013); (Manza et al., 2020); (C. Wang et al., 2020); (D. Liu et al., 2022); (Ketchabaw et al., 2022); (Göttlich et al., 2017); (Borchardt et al., 2015); (Ekman et al., 2012); (L. Wang et al., 2010); (Shine et al., 2016); (Lin et al., 2022); (X. Zhang et al., 2015); (Yi et al., 2023); (J. Zhang et al., 2021); (Y. Fan et al., 2021); (Betzel et al., 2020); (Jin et al., 2020); (Moorthigari et al., 2020); (Tipnis et al., 2020); (Amlien et al., 2019); (Ebrahimi et al., 2019); (Xi et al., 2019); (Agrawal et al., 2019); (Cao et al., 2019); (Huskey et al., 2018); (Kim et al., 2018); (Reddy et al., 2018); (Knyazeva et al., 2018); (Zamroziewicz et al., 2017); (Ma & Zhang, 2017); (Y. Liu et al., 2017); (Berroir et al., 2017); (Mancini et al., 2017); (Bolt et al., 2017); (Prčkovska et al., 2016); (Anderson et al., 2017); (Shang et al., 2017); (Zhao et al., 2017); (Hilger et al., 2017a); (Gratton et al., 2016); (File et al., 2016); (Najafi et al., 2016); (Alavash et al., 2016); (Liang et al., 2016); (Du et al., 2015); (Alavash, Hilgetag, et al., 2015); (Wen et al., 2015); (X. Xu et al., 2015); (Cocchi et al., 2015); (Taya et al., 2014); (DeSalvo et al., 2014); (Arnold et al., 2014); (Koelsch & Skouras, 2014); (Sami & Miall, 2013); (Liang et al., 2013); (Moussa et al., 2012); (Satterthwaite et al., 2012); (Messé et al., 2012); (Sheppard et al., 2011); (Richards et al., 2018); (Martial et al., 2023); (Dan et al., 2023); (Breedt et al., 2022); (Choi et al., 2023); (X. Chen et al., 2023); (Han et al., 2023); (Y. Gao et al., 2023); (Khodaei et al., 2023); (Ryu et al., 2022); (Ghiles et al., 2023); (H. Wang et al., 2023); (Lee et al., 2022) |
| | Partial | (Foo et al., 2021); (Lloyd, 2020); (H. Zhang et al., 2012); (Jacob et al., 2016); (Burdette et al., 2010) |
| | WaveletCoherence | (Vriend et al., 2020); (Schlesinger et al., 2017); (Q. Li et al., 2019); (Breckel et al., 2013); (Suprano et al., 2019); (S. Gu et al., 2015) |
| | Spearman | (Thompson & Fransson, 2015); (Franzmeier et al., 2018) |
| | MultipleC | (Y. Wang et al., 2021); (Kruschwitz et al., 2018); (G. Zhang & Liu, 2021); (Parhizi et al., 2018); (Liang et al., 2012); (de Pasquale et al., 2017); (Spreng et al., 2013) |
| | Not reported | |
| ConnectNormalize | | (Ray et al., 2020); (Agrawal et al., 2019); (Alavash, Doebler, et al., 2015); (Crossley et al., 2013); (Ekman et al., 2012); (Göttlich et al., 2017); (Alavash et al., 2016); (Taya et al., 2014); (Setton et al., 2022); (L. Song et al., 2020); (Zhong et al., 2014); (Bartholomew et al., 2019); (Danti et al., 2018); (S. Zhang et al., 2022); (Vatansever et al., 2015a); (Duan et al., 2014); (Farah & Horowitz-Kraus, 2019); (D. Liu et al., 2022); (Amlien et al., 2019); (de Pasquale et al., 2017); (Ryu et al., 2022); (X. Liu et al., 2020); (X. Fan et al., 2021); (Finc et al., 2017); (Cole et al., 2015); (Lunsford-Avery et al., 2020); (Qin et al., 2016); (Marek et al., 2015); (File et al., 2016); (Arnold |

| | | |
|---|---|---|
| | | et al., 2014); (Madden et al., 2020); (Farahani et al., 2022); (Varangis et al., 2019); (J. Wang et al., 2017); (Pamplona et al., 2015); (Iordan et al., 2018); (Reineberg & Banich, 2016); (Markett et al., 2013); (Finc et al., 2020); (Sun et al., 2017); (Quante et al., 2018); (Gozdas et al., 2019); (He et al., 2019); (Pindus et al., 2020); (J. Liu et al., 2017); (Vatansever et al., 2015b); (Shine et al., 2016); (Zamroziewicz et al., 2017); (X. Chen et al., 2023); (Kobayashi et al., 2020); (Malagurski et al., 2020); (Le et al., 2020); (Jin et al., 2020); (H. Wang et al., 2023); (Lee et al., 2022); (Zuo et al., 2018); (Pan et al., 2018); (C. Zhang et al., 2016); (Jung et al., 2018); (Dan et al., 2023); (Han et al., 2023); (Ogawa, 2021); (Y. Fan et al., 2019); (Hearne et al., 2017); (Ketchabaw et al., 2022); (X. Xu et al., 2015); (Fujimoto et al., 2020); (Santarnecchi et al., 2014); (Gratton et al., 2016); (Liao et al., 2017); (Kim et al., 2018); (Tooley et al., 2022); (Khodaei et al., 2023); (Fukushima & Sporns, 2018); (Fukushima et al., 2017) |
| NetworkDefine | CommunityDetection | (Kobayashi et al., 2020); (Su et al., 2021); (Vriend et al., 2020); (Varangis et al., 2021); (G. Zhang & Liu, 2021); (J. Song et al., 2014); (Madden et al., 2020); (Varangis et al., 2019); (Le et al., 2020); (Servaas et al., 2017); (Chong et al., 2019); (Cohen & D'Esposito, 2016); (Monge et al., 2017); (C. Zhang et al., 2016); (Westphal et al., 2017); (He et al., 2019); (Pindus et al., 2020); (Iordan et al., 2018); (Jarrahi & Kollias, 2020); (T. Chen et al., 2016); (Huckins et al., 2019); (Ding et al., 2011); (Neyland et al., 2021); (Ray et al., 2020); (Y. Fan et al., 2019); (Lunsford-Avery et al., 2020); (Finc et al., 2020); (Tooley et al., 2022); (Marek et al., 2015); (Crossley et al., 2013); (Manza et al., 2020); (C. Wang et al., 2020); (Göttlich et al., 2017); (Borchardt et al., 2015); (Shine et al., 2016); (Y. Fan et al., 2021); (Betzel et al., 2020); (Cao et al., 2019); (Kim et al., 2018); (Reddy et al., 2018); (Knyazeva et al., 2018); (de Pasquale et al., 2017); (Ma & Zhang, 2017); (Mancini et al., 2017); (Shang et al., 2017); (File et al., 2016); (Najafi et al., 2016); (Alavash et al., 2016); (S. Gu et al., 2015); (Du et al., 2015); (Alavash, Hilgetag, et al., 2015); (Cocchi et al., 2015); (DeSalvo et al., 2014); (Liang et al., 2013); (Spreng et al., 2013); (Moussa et al., 2012); (Satterthwaite et al., 2012); (Messé et al., 2012); (Khodaei et al., 2023) |
| | 7RSNYeo | (Fukushima et al., 2017); (Foo et al., 2021); (Farahani et al., 2022); (Fukushima & Sporns, 2018); (Bueichekú et al., 2019); (Yi et al., 2023); (Tipnis et al., 2020); (Han et al., 2023); (Y. Gao et al., 2023) |
| | 17RSNYeo | (J. Wang et al., 2017) |
| | PowerNetwork | |
| | Cole | (Vatansever et al., 2015a) |
| | Shirer | (Belden et al., 2020); (Smith et al., 2018) |
| | Schlesinger | (Malagurski et al., 2020) |
| | Damoiseaux | (Alluri et al., 2017) |
| | StudySpecificN | (Vatansever et al., 2015b); (C. Wang et al., 2022); (Liang et al., 2016); (Richards et al., 2018) |
| | Not reported | |
| GraphDefine | Binary | (Fujimoto et al., 2020); (Kobayashi et al., 2020); (Zhou et al., 2021); (Zheng et al., 2021); (C. Yan & He, 2011); (Zuo et al., 2018); (Ogawa, 2021); (Hayasaka, 2013); (Parhizi et al., 2018); (Gozdas et al., 2019); (Stevens et al., 2012); (Tian et al., 2011); (Hilger et al., 2017b); (Feng et al., 2015); (Bartholomew et al., 2019); (Bailey et al., 2018); (Le et al., 2020); (Pan et al., 2018); (Kawagoe et al., 2017); (Yue et al., 2017); (Cohen & D'Esposito, 2016); (X. Liu et al., 2018); (C. Zhang et al., 2016); (Iordan et al., 2018); (Rzucidlo et al., 2013); (Deng et al., |

| | | |
|---|---|---|
| | | 2016); (Belden et al., 2020); (Braun et al., 2012); (Ding et al., 2011); (Lloyd, 2020); (Neyland et al., 2021); (Alluri et al., 2017); (Ray et al., 2020); (Breckel et al., 2013); (H. Zhang et al., 2012); (Santarnecchi et al., 2014); (Gopinath et al., 2015); (Geerligs et al., 2015); (Alavash, Doebler, et al., 2015); (Wu et al., 2013); (Smith et al., 2018); (P. Xu et al., 2014); (Xiao et al., 2016); (Suprano et al., 2019); (Y. Gu et al., 2022); (Polanía et al., 2011); (Crossley et al., 2013); (C. Wang et al., 2020); (Ekman et al., 2012); (Burdette et al., 2010); (J. Zhang et al., 2021); (Jin et al., 2020); (Amlien et al., 2019); (Huskey et al., 2018); (Ma & Zhang, 2017); (Mancini et al., 2017); (Prčkovska et al., 2016); (Anderson et al., 2017); (Shang et al., 2017); (Hilger et al., 2017a); (Najafi et al., 2016); (Liang et al., 2016); (Du et al., 2015); (Wen et al., 2015); (X. Xu et al., 2015); (Cocchi et al., 2015); (Sami & Miall, 2013); (Moussa et al., 2012); (Sheppard et al., 2011); (Martial et al., 2023); (Breedt et al., 2022); (Lee et al., 2022) |
| | Weighted | (Fukushima et al., 2017); (Gracia-Tabuenca et al., 2021); (Foo et al., 2021); (Quante et al., 2018); (Tooley et al., 2020); (Vriend et al., 2020); (Varangis et al., 2021); (G. Zhang & Liu, 2021); (X. Fan et al., 2021); (Malagurski et al., 2020); (Madden et al., 2020); (Evensmoen et al., 2021); (Pezoulas et al., 2017); (Markett et al., 2018); (L. Song et al., 2020); (Varangis et al., 2019); (Fukushima & Sporns, 2018); (J. Wang et al., 2017); (F. Fan et al., 2021); (Servaas et al., 2017); (Chong et al., 2019); (Monge et al., 2017); (Xia et al., 2019); (Liao et al., 2017); (Westphal et al., 2017); (He et al., 2019); (Pindus et al., 2020); (S. Zhang et al., 2022); (Schlesinger et al., 2017); (Geib et al., 2017); (T. Chen et al., 2016); (Huckins et al., 2019); (Cole et al., 2015); (Vatansever et al., 2015a); (Sheppard et al., 2011); (Geerligs et al., 2014); (Duan et al., 2014); (Q. Li et al., 2019); (Meunier et al., 2014); (H. Yan et al., 2022); (Koba et al., 2021); (Sato et al., 2015); (Y. Fan et al., 2019); (Lunsford-Avery et al., 2020); (Reineberg & Banich, 2016); (Markett et al., 2013); (J. Liu et al., 2017); (Rubin et al., 2017); (Vatansever et al., 2015b); (Qin et al., 2016); (Hearne et al., 2017); (Ginestet & Simmons, 2011); (Jung et al., 2018); (Finc et al., 2020); (Spielberg et al., 2015); (Sun et al., 2017); (Manza et al., 2020); (Borchardt et al., 2015); (L. Wang et al., 2010); (Lin et al., 2022); (X. Zhang et al., 2015); (Moorthigari et al., 2020); (Ebrahimi et al., 2019); (Huang et al., 2018); (Reddy et al., 2018); (Knyazeva et al., 2018); (Aggarwal et al., 2017); (Zamroziewicz et al., 2017); (de Pasquale et al., 2017); (Y. Liu et al., 2017); (Berroir et al., 2017); (de Paula et al., 2017); (Gratton et al., 2016); (File et al., 2016); (Alavash et al., 2016); (S. Gu et al., 2015); (Thompson & Fransson, 2015); (Alavash, Hilgetag, et al., 2015); (Taya et al., 2014); (DeSalvo et al., 2014); (Arnold et al., 2014); (Liang et al., 2013); (Spreng et al., 2013); (Satterthwaite et al., 2012); (Messé et al., 2012); (Dan et al., 2023); (Choi et al., 2023); (X. Chen et al., 2023); (Y. Gao et al., 2023); (Ryu et al., 2022); (Ghiles et al., 2023); (H. Wang et al., 2023) |
| | MeasureDependent | (Yin et al., 2019); (J. Song et al., 2014); (Farahani et al., 2022); (Pamplona et al., 2015); (Finc et al., 2017); (Liang et al., 2012); (Bottino et al., 2021); (Tooley et al., 2022); (Cao et al., 2019); (Bolt et al., 2017); (Khodaei et al., 2023) |
| | Not reported | (Shen et al., 2013); (S. Gao et al., 2021); (Y. Wang et al., 2021); (Su et al., 2021); (Masuda et al., 2018); (Kruschwitz et al., 2018); (X. Liu et al., 2020); (Setton et al., 2022); (Danti et al., 2018); (Long et al., 2017); (Jarrahi & Kollias, 2020); (Taruffi et al., 2017); (Zhong et al., 2014); (Beaty et al., 2015); (Farah & Horowitz-Kraus, 2019); (Sheppard et al., 2012); (Orwig et al., 2021); (Bueichekú et al., 2019); (Kolskår et al., 2018); (C. Wang et al., 2022); (Sreenivasan et al., 2017); (X. Li et al., 2020); (Marek et al., 2015); (Alnæs et al., 2015); (D. Liu et al., 2022); (Jacob et al., 2016); (Ketchabaw et al., 2022); (Göttlich et al., 2017); (Shine et al., 2016); (Y. Zhang et al., 2022); (Yi et al., 2023); (Y. Fan et al., 2021); (Betzel et al., 2020); (Tipnis et al., 2020); (Xi et al., 2019); (Agrawal et al., 2019); (Brandl et al., 2018); (Kim et al., 2018); (Zhao et al., 2017); (Koelsch & Skouras, |

| | | |
|---|---|---|
| | | 2014); (Franzmeier et al., 2018); (Richards et al., 2018); (Han et al., 2023); (Yang et al., 2023); (Invernizzi et al., 2023) |
| NegCCHandle | Keep | Fukushima et al., 2017); (Gracia-Tabuenca et al., 2021); (Tooley et al., 2020); (Fukushima & Sporns, 2018); (Pamplona et al., 2015); (Iordan et al., 2018); (T. Chen et al., 2016); (Duan et al., 2014); (Meunier et al., 2014); (Betzel et al., 2020); (Alavash et al., 2016); (Satterthwaite et al., 2012); (Dan et al., 2023) |
| | Zero | (Foo et al., 2021); (Y. Wang et al., 2021); (Quante et al., 2018); (Zhou et al., 2021); (Zheng et al., 2021); (Zuo et al., 2018); (X. Fan et al., 2021); (Malagurski et al., 2020); (Madden et al., 2020); (Pezoulas et al., 2017); (Hilger et al., 2017b); (Varangis et al., 2019); (Danti et al., 2018); (Monge et al., 2017); (X. Liu et al., 2018); (Liao et al., 2017); (Pindus et al., 2020); (Y. Fan et al., 2019); (Bueichekú et al., 2019); (Reineberg & Banich, 2016); (Markett et al., 2013); (J. Liu et al., 2017); (Xiao et al., 2016); (C. Wang et al., 2022); (Jung et al., 2018); (Finc et al., 2020); (Spielberg et al., 2015); (Sun et al., 2017); (Y. Gu et al., 2022); (Manza et al., 2020); (Shine et al., 2016); (Lin et al., 2022); (Zamroziewicz et al., 2017); (Y. Liu et al., 2017); (Bolt et al., 2017); (Prčkovska et al., 2016); (Hilger et al., 2017a); (File et al., 2016); (Najafi et al., 2016); (Liang et al., 2016); (Du et al., 2015); (DeSalvo et al., 2014); (Liang et al., 2013); (Spreng et al., 2013); (Moussa et al., 2012); (Franzmeier et al., 2018); (Choi et al., 2023); (X. Chen et al., 2023); (Khodaei et al., 2023); (H. Wang et al., 2023) |
| | LinearScale | |
| | AbsoluteN | (Hayasaka, 2013); (Farahani et al., 2022); (Feng et al., 2015); (Yue et al., 2017); (Braun et al., 2012); (Finc et al., 2017); (Suprano et al., 2019); (Tooley et al., 2022); (L. Wang et al., 2010); (Jin et al., 2020); (Knyazeva et al., 2018); (Alavash, Hilgetag, et al., 2015); (Messé et al., 2012); (Sheppard et al., 2011); (Breedt et al., 2022); (Ryu et al., 2022); (Ghiles et al., 2023) |
| | Not reported | (Shen et al., 2013); (S. Gao et al., 2021); (Fujimoto et al., 2020); (Kobayashi et al., 2020); (Su et al., 2021); (C. Yan & He, 2011); (Masuda et al., 2018); (Vriend et al., 2020); (Varangis et al., 2021); (Ogawa, 2021); (Kruschwitz et al., 2018); (Yin et al., 2019); (G. Zhang & Liu, 2021); (X. Liu et al., 2020); (J. Song et al., 2014); (Setton et al., 2022); (Evensmoen et al., 2021); (Parhizi et al., 2018); (Gozdas et al., 2019); (Stevens et al., 2012); (Tian et al., 2011); (L. Song et al., 2020); (Bartholomew et al., 2019); (Bailey et al., 2018); (Le et al., 2020); (J. Wang et al., 2017); (Pan et al., 2018); (Long et al., 2017); (Kawagoe et al., 2017); (F. Fan et al., 2021); (Servaas et al., 2017); (Chong et al., 2019); (Cohen & D'Esposito, 2016); (Xia et al., 2019); (C. Zhang et al., 2016); (Westphal et al., 2017); (He et al., 2019); (S. Zhang et al., 2022); (Jarrahi & Kollias, 2020); (Schlesinger et al., 2017); (Rzucidlo et al., 2013); (Geib et al., 2017); (Deng et al., 2016); (Taruffi et al., 2017); (Belden et al., 2020); (Huckins et al., 2019); (Ding et al., 2011); (Cole et al., 2015); (Zhong et al., 2014); (Vatansever et al., 2015a); (Sheppard et al., 2011); (Lloyd, 2020); (Liang et al., 2012); (Geerligs et al., 2014); (Beaty et al., 2015); (Bottino et al., 2021); (Q. Li et al., 2019); (Farah & Horowitz-Kraus, 2019); (Neyland et al., 2021); (H. Yan et al., 2022); (Alluri et al., 2017); (Koba et al., 2021); (Ray et al., 2020); (Breckel et al., 2013); (H. Zhang et al., 2012); (Santarnecchi et al., 2014); (Sato et al., 2015); (Sheppard et al., 2012); (Orwig et al., 2021); (Gopinath et al., 2015); (Lunsford-Avery et al., 2020); (Geerligs et al., 2015); (Alavash, Doebler, et al., 2015); (Wu et al., 2013); (Smith et al., 2018); (P. Xu et al., 2014); (Kolskår et al., 2018); (Rubin et al., 2017); (Vatansever et al., 2015b); (Qin et al., 2016); (Hearne et al., 2017); (Sreenivasan et al., 2017); (Ginestet & Simmons, 2011); (X. Li et al., 2020); (Polanía et al., 2011); (Marek et al., 2015); (Crossley et al., 2013); (Alnæs et al., 2015); (C. Wang et al., 2020); (D. Liu et al., 2022); (Jacob et al., 2016); (Ketchabaw et al., 2022); (Göttlich et al., 2017); (Borchardt et al., 2015); (Ekman et al., 2012); (Burdette et al., 2010); (X. Zhang et al., 2015); (Y. Zhang et al., 2022); (Yi et |

| | | |
|---|---|---|
| | | al., 2023); (J. Zhang et al., 2021); (Y. Fan et al., 2021); (Tipnis et al., 2020); (Amlien et al., 2019); (Ebrahimi et al., 2019); (Xi et al., 2019); (Agrawal et al., 2019); (Cao et al., 2019); (Huskey et al., 2018); (Brandl et al., 2018); (Kim et al., 2018); (Huang et al., 2018); (Reddy et al., 2018); (Aggarwal et al., 2017); (de Pasquale et al., 2017); (Ma & Zhang, 2017); (Berroir et al., 2017); (Mancini et al., 2017); (de Paula et al., 2017); (Anderson et al., 2017); (Shang et al., 2017); (Zhao et al., 2017); (Gratton et al., 2016); (S. Gu et al., 2015); (Thompson & Fransson, 2015); (Wen et al., 2015); (X. Xu et al., 2015); (Cocchi et al., 2015); (Taya et al., 2014); (Arnold et al., 2014); (Koelsch & Skouras, 2014); (Sami & Miall, 2013); (Richards et al., 2018); (Martial et al., 2023); (Han et al., 2023); (Y. Gao et al., 2023); (Yang et al., 2023); (Invernizzi et al., 2023); (Lee et al., 2022) |
| SparseControl | AbsoluteT | (Kobayashi et al., 2020); (Parhizi et al., 2018); (Stevens et al., 2012); (L. Song et al., 2020); (Bailey et al., 2018); (J. Wang et al., 2017); (Pan et al., 2018); (F. Fan et al., 2021); (Chong et al., 2019); (X. Liu et al., 2018); (Pamplona et al., 2015); (S. Zhang et al., 2022); (Deng et al., 2016); (Belden et al., 2020); (Ding et al., 2011); (Sheppard et al., 2011); (Lunsford-Avery et al., 2020); (Smith et al., 2018); (J. Liu et al., 2017); (Xiao et al., 2016); (Lin et al., 2022); (Du et al., 2015); (Alavash, Hilgetag, et al., 2015); (Arnold et al., 2014); (Sami & Miall, 2013); (Franzmeier et al., 2018); (X. Chen et al., 2023); (Y. Gao et al., 2023); (Ghiles et al., 2023); (H. Wang et al., 2023) |
| | Density | (Fujimoto et al., 2020); (Y. Wang et al., 2021); (Su et al., 2021); (Zhou et al., 2021); (Zheng et al., 2021); (C. Yan & He, 2011); (Zuo et al., 2018); (Ogawa, 2021); (Kruschwitz et al., 2018); (Tian et al., 2011); (Farahani et al., 2022); (Hilger et al., 2017b); (Varangis et al., 2019); (Feng et al., 2015); (Le et al., 2020); (Kawagoe et al., 2017); (Servaas et al., 2017); (Yue et al., 2017); (Cohen & D'Esposito, 2016); (Xia et al., 2019); (C. Zhang et al., 2016); (Liao et al., 2017); (Westphal et al., 2017); (Iordan et al., 2018); (Huckins et al., 2019); (Braun et al., 2012); (Finc et al., 2017); (Cole et al., 2015); (Lloyd, 2020); (Bottino et al., 2021); (H. Yan et al., 2022); (Koba et al., 2021); (Ray et al., 2020); (Santarnecchi et al., 2014); (Gopinath et al., 2015); (Geerligs et al., 2015); (Reineberg & Banich, 2016); (Alavash, Doebler, et al., 2015); (Wu et al., 2013); (P. Xu et al., 2014); (Vatansever et al., 2015b); (C. Wang et al., 2022); (Hearne et al., 2017); (Sreenivasan et al., 2017); (Suprano et al., 2019); (X. Li et al., 2020); (Spielberg et al., 2015); (Tooley et al., 2022); (Sun et al., 2017); (Y. Gu et al., 2022); (Marek et al., 2015); (C. Wang et al., 2020); (D. Liu et al., 2022); (Göttlich et al., 2017); (Borchardt et al., 2015); (Ekman et al., 2012); (L. Wang et al., 2010); (Shine et al., 2016); (J. Zhang et al., 2021); (Amlien et al., 2019); (Cao et al., 2019); (Huskey et al., 2018); (Brandl et al., 2018); (Kim et al., 2018); (Huang et al., 2018); (Ma & Zhang, 2017); (Mancini et al., 2017); (Bolt et al., 2017); (de Paula et al., 2017); (Anderson et al., 2017); (Shang et al., 2017); (Hilger et al., 2017a); (Gratton et al., 2016); (Najafi et al., 2016); (Alavash et al., 2016); (Liang et al., 2016); (Thompson & Fransson, 2015); (X. Xu et al., 2015); (Cocchi et al., 2015); (Taya et al., 2014); (DeSalvo et al., 2014); (Ryu et al., 2022); (Lee et al., 2022) |
| | SignificantEdges | (X. Fan et al., 2021); (Gozdas et al., 2019); (Bartholomew et al., 2019); (Pindus et al., 2020); (Jarrahi & Kollias, 2020); (Zhong et al., 2014); (Duan et al., 2014); (Q. Li et al., 2019); (Rubin et al., 2017); (Crossley et al., 2013); (Zamroziewicz et al., 2017); (de Pasquale et al., 2017); (Y. Liu et al., 2017); (Berroir et al., 2017); (Prčkovska et al., 2016); (Wen et al., 2015) |
| | GraphCost | Hayasaka, 2013); (Yin et al., 2019); (J. Song et al., 2014); (Rzucidlo et al., 2013); (Neyland et al., 2021); (Alluri et al., 2017); (Breckel et al., 2013); (Sheppard et al., 2012); (Polanía et al., 2011); (Burdette et al., 2010); (X. Zhang et al., 2015); (Jin et al., 2020); (Ebrahimi et al., 2019); (Zhao et al., 2017); (Moussa et al., 2012); (Sheppard et al., 2011) |

| | | |
|---|---|---|
| | Full | (Fukushima et al., 2017); (Gracia-Tabuenca et al., 2021); (Quante et al., 2018); (Tooley et al., 2020); (Vriend et al., 2020); (Varangis et al., 2021); (G. Zhang & Liu, 2021); (Malagurski et al., 2020); (Pezoulas et al., 2017); (Markett et al., 2018); (Danti et al., 2018); (Fukushima & Sporns, 2018); (Monge et al., 2017); (Geib et al., 2017); (T. Chen et al., 2016); (Vatansever et al., 2015a); (Geerligs et al., 2014); (Markett et al., 2013); (File et al., 2016); (Liang et al., 2013); (Spreng et al., 2013); (Satterthwaite et al., 2012); (Messé et al., 2012); (Dan et al., 2023); (Breedt et al., 2022) |
| | Not reported | (Shen et al., 2013); (S. Gao et al., 2021); (Foo et al., 2021); (Masuda et al., 2018); (X. Liu et al., 2020); (Setton et al., 2022); (Madden et al., 2020); (Evensmoen et al., 2021); (Long et al., 2017); (He et al., 2019); (Schlesinger et al., 2017); (Taruffi et al., 2017); (Liang et al., 2012); (Beaty et al., 2015); (Farah & Horowitz-Kraus, 2019); (Meunier et al., 2014); (H. Zhang et al., 2012); (Sato et al., 2015); (Y. Fan et al., 2019); (Orwig et al., 2021); (Bueichekú et al., 2019); (Kolskår et al., 2018); (Qin et al., 2016); (Ginestet & Simmons, 2011); (Jung et al., 2018); (Finc et al., 2020); (Manza et al., 2020); (Alnæs et al., 2015); (Jacob et al., 2016); (Ketchabaw et al., 2022); (Y. Zhang et al., 2022); (Yi et al., 2023); (Y. Fan et al., 2021); (Betzel et al., 2020); (Moorthigari et al., 2020); (Tipnis et al., 2020); (Xi et al., 2019); (Agrawal et al., 2019); (Reddy et al., 2018); (Knyazeva et al., 2018); (Aggarwal et al., 2017); (S. Gu et al., 2015); (Koelsch & Skouras, 2014); (Richards et al., 2018); (Martial et al., 2023); (Choi et al., 2023); (Han et al., 2023); (Khodaei et al., 2023); (Yang et al., 2023); (Invernizzi et al., 2023) |
| GraphMeasDefine | | (Ginestet & Simmons, 2011); (Zhao et al., 2017); (Taruffi et al., 2017); (Beaty et al., 2015); (H. Yan et al., 2022); (Agrawal et al., 2019); (Koelsch & Skouras, 2014); (Masuda et al., 2018); (Evensmoen et al., 2021); (Geib et al., 2017); (Sheppard et al., 2011); (H. Zhang et al., 2012); (Kolskår et al., 2018); (Sreenivasan et al., 2017); (Yi et al., 2023); (C. Yan & He, 2011); (Parhizi et al., 2018); (Tian et al., 2011); (Rzucidlo et al., 2013); (Liang et al., 2012); (Geerligs et al., 2014); (Bottino et al., 2021); (Meunier et al., 2014); (Ray et al., 2020); (Gopinath et al., 2015); (Polanía et al., 2011); (L. Wang et al., 2010); (Burdette et al., 2010); (Y. Fan et al., 2021); (Moorthigari et al., 2020); (Xi et al., 2019); (Knyazeva et al., 2018); (de Paula et al., 2017); (Setton et al., 2022); (Ekman et al., 2012); (Huskey et al., 2018); (Huang et al., 2018); (Messé et al., 2012); (Y. Wang et al., 2021); (Braun et al., 2012); (Zhong et al., 2014); (Farah & Horowitz-Kraus, 2019); (Breckel et al., 2013); (Sheppard et al., 2012); (Alavash, Doebler, et al., 2015); (P. Xu et al., 2014); (Rubin et al., 2017); (Crossley et al., 2013); (Göttlich et al., 2017); (Reddy et al., 2018); (Aggarwal et al., 2017); (Taya et al., 2014); (Sami & Miall, 2013); (Hayasaka, 2013); (L. Song et al., 2020); (Bailey et al., 2018); (Kawagoe et al., 2017); (Ding et al., 2011); (Alluri et al., 2017); (Bueichekú et al., 2019); (Wu et al., 2013); (X. Li et al., 2020); (Spielberg et al., 2015); (D. Liu et al., 2022); (Brandl et al., 2018); (Prčkovska et al., 2016); (Anderson et al., 2017); (Alavash, Hilgetag, et al., 2015); (Wen et al., 2015); (Moussa et al., 2012); (Sheppard et al., 2011); (Franzmeier et al., 2018); (Breedt et al., 2022); (Ghiles et al., 2023); (Yang et al., 2023); (Zhou et al., 2021); (X. Liu et al., 2020); (Danti et al., 2018); (S. Zhang et al., 2022); (Qin et al., 2016); (X. Zhang et al., 2015); (Amlien et al., 2019); (Cao et al., 2019); (Y. Liu et al., 2017); (Alavash et al., 2016); (S. Gu et al., 2015); (Choi et al., 2023); (Su et al., 2021); (Zheng et al., 2021); (Varangis et al., 2021); (Kruschwitz et al., 2018); (G. Zhang & Liu, 2021); (Cohen & D'Esposito, 2016); (Schlesinger et al., 2017); (Cole et al., 2015); (Vatansever et al., 2015a); (Duan et al., 2014); (Q. Li et al., 2019); (Geerligs et al., 2015); (Marek et al., 2015); (Lin et al., 2022); (de Pasquale et al., 2017); (Berroir et al., 2017); (Cocchi et al., 2015); (Arnold et al., 2014); (Richards et al., 2018); (Ryu et al., 2022); (Invernizzi et al., 2023); (Vriend et al., 2020); (X. Fan et al., 2021); (Hilger et al., 2017b); (Servaas et al., 2017); (Deng et al., 2016); (Belden et al., 2020); (Finc et al., 2017); (Koba et al., 2021); (Lunsford-Avery et al., 2020); (Xiao et al., 2016); (Spreng et al., |

| | | |
|---|---|---|
| | | 2013); (Madden et al., 2020); (Gozdas et al., 2019); (Stevens et al., 2012); (Markett et al., 2018); (Feng et al., 2015); (J. Wang et al., 2017); (Yue et al., 2017); (Monge et al., 2017); (Westphal et al., 2017); (He et al., 2019); (Pamplona et al., 2015); (Reineberg & Banich, 2016); (Markett et al., 2013); (Suprano et al., 2019); (Finc et al., 2020); (Sun et al., 2017); (Ebrahimi et al., 2019); (Ma & Zhang, 2017); (Mancini et al., 2017); (Shang et al., 2017); (File et al., 2016); (Liang et al., 2016); (DeSalvo et al., 2014); (Liang et al., 2013); (Martial et al., 2023); (Y. Gao et al., 2023); (Foo et al., 2021); (Quante et al., 2018); (Farahani et al., 2022); (Varangis et al., 2019); (X. Liu et al., 2018); (Iordan et al., 2018); (Jarrahi & Kollias, 2020); (Huckins et al., 2019); (Neyland et al., 2021); (Smith et al., 2018); (J. Liu et al., 2017); (Vatansever et al., 2015b); (C. Wang et al., 2020); (Ketchabaw et al., 2022); (Borchardt et al., 2015); (Shine et al., 2016); (J. Zhang et al., 2021); (Zamroziewicz et al., 2017); (X. Chen et al., 2023); (S. Gao et al., 2021); (Kobayashi et al., 2020); (Gracia-Tabuenca et al., 2021); (Le et al., 2020); (Chong et al., 2019); (Pindus et al., 2020); (Jung et al., 2018); (Y. Gu et al., 2022); (Manza et al., 2020); (Betzel et al., 2020); (Jin et al., 2020); (Tipnis et al., 2020); (Du et al., 2015); (Satterthwaite et al., 2012); (H. Wang et al., 2023); (Lee et al., 2022); (Zuo et al., 2018); (Malagurski et al., 2020); (J. Song et al., 2014); (Pezoulas et al., 2017); (Pan et al., 2018); (C. Zhang et al., 2016); (Sato et al., 2015); (C. Wang et al., 2022); (Hearne et al., 2017); (Najafi et al., 2016); (X. Xu et al., 2015); (Dan et al., 2023); (Fujimoto et al., 2020); (Tooley et al., 2020); (Ogawa, 2021); (F. Fan et al., 2021); (Santarnecchi et al., 2014); (Y. Fan et al., 2019); (Han et al., 2023); (T. Chen et al., 2016); (Kim et al., 2018); (Gratton et al., 2016); (Yin et al., 2019); (Liao et al., 2017); (Thompson & Fransson, 2015); (Tooley et al., 2022); (Bolt et al., 2017); (Khodaei et al., 2023); (Fukushima & Sporns, 2018) |
| ResultAggregate | Average | (Hilger et al., 2017b); (Kawagoe et al., 2017); (Deng et al., 2016); (Breckel et al., 2013); (Marek et al., 2015); (Cao et al., 2019); (Hilger et al., 2017a) |
| | Optimal | (Servaas et al., 2017); (Bottino et al., 2021); (Koba et al., 2021); (Brandl et al., 2018); (Alavash et al., 2016) |
| | AUC | (Chong et al., 2019); (Westphal et al., 2017); (S. Zhang et al., 2022); (Liang et al., 2012); (Rubin et al., 2017); (Sreenivasan et al., 2017); (X. Li et al., 2020); (Spielberg et al., 2015); (D. Liu et al., 2022); (J. Zhang et al., 2021); (Y. Fan et al., 2021); (Jin et al., 2020); (Bolt et al., 2017) |
| | Not reported | (Spreng et al., 2013) |
